# N6-Methyladenine DNA modification modulates pathogen virulence in nematodes

**DOI:** 10.1101/2025.09.03.674121

**Authors:** Dadong Dai, Shurong Zhang, Boyan Hu, Yayi Zhou, Simeng Cui, Jia Sun, Yali Zhang, Xueyu Wang, Shahid Siddique, Dexin Bo, Min Zhang, Valerie M Williamson, Haozhe Yao, Xinya Duan, Wentao Wu, Donghai Peng, Jinshui Zheng, Ming Sun

## Abstract

Understanding the global regulatory mechanisms that control pathogen virulence gene expression is essential for elucidating the molecular basis of pathogenicity. N6-methyladenine (6mA) plays a crucial role in regulating gene expression in response to various environmental stresses; however, its role in pathogen virulence remains largely unexplored. Here, we report the widespread occurrence of 6mA across 17 nematode isolates and map its genomic landscape in six notorious agriculturally important pathogen root-knot nematodes (RKNs). We demonstrated that 6mA is characterized by a conserved GAG motif across nematodes, but exhibits species-specific distribution patterns and distinct effects on gene expression. In particular, its enrichment in transposable elements differs between polyploid and diploid nematodes, suggesting lineage-specific epigenetic regulation potentially associated with polyploidy. We further identified two functional 6mA demethylases, MiNMAD-1 and MiNMAD-2, and confirmed their catalytic activity and active sites. Host-induced gene silencing of minmad-1 significantly increased plant resistance to three polyploid RKN species. A detailed functional analysis revealed that minmad-1 knockdown disrupted virulence gene expression during the parasitic stage, thereby reducing nematode infectivity. Together, our findings suggest 6mA demethylase as a key epigenetic regulator of RKNs’ virulence, providing new insights into nematode biology and offering promising targets for the development of sustainable control strategies.

**Significance Statement:** 6mA methyltransferase regulates the expression of virulence genes in certain pathogenic bacteria and plays a critical role in their infectivity. However, the regulation of virulence gene expression in eukaryotic pathogens, particularly plant pathogens, remains poorly understood. Most studies on pathogen virulence have focused on individual effectors, with little insight into global regulatory mechanisms. Here, we demonstrate that 6mA broadly shapes virulence gene expression patterns in polyploid RKNs. Moreover, transgenic tobacco, tomato, and rice plants expressing dsRNA against nematode demethylases showed significantly enhanced resistance to RKNs. These findings establish 6mA demethylase as a promising epigenetic target for controlling plant-parasitic nematodes.

## Introduction

The regulation of pathogen virulence gene expression is crucial to understanding pathogenicity mechanisms. While current research has largely focuses on the functional analysis of individual virulence genes, little is known about the broader mechanisms underlying global virulence regulation in pathogens. DNA methylation, one of the earliest discovered DNA modifications, influences chromatin structure, DNA conformation, stability, and protein interactions, thereby modulating gene expression(1, 2). The role of 6mA in bacterial virulence is well established: DNA adenine methylase (Dam) is essential for *Salmonella* virulence(3), and Dam(-) mutants exhibit multiple virulence-related defects(4).

Dam also regulates effectors such as SpoA and SopB, the latter possessing phosphoinositide phosphatase activity(5, 6), and has also been reported to globally modulate the expression of cariogenicity-associated genes in *Streptococcus mutans*(7). Additionally, 6mA methyltransferase have been shown to regulate virulence gene expression in pathogenic bacteria that cause pediatric osteoarticular infections and enterocolitis(8, 9), as well as in the opportunistic pathogen *Pseudomonas aeruginosa*, which induces both acute and chronic infections(10). In *Streptococcus pyogenes*, 6mA also modulates the expression of Mga, a major transcriptional regulator of multiple virulence genes (11). In eukaryotes, evidence remains limited, with only a few studies reporting 6mA-regulated virulence in fungi. For example, 6mA methyltransferase is critical for the pathogenicity of the plant fungal pathogen *Cryphonectria parasitica*(12). However, most investigations linking 6mA to virulence have focused on animal pathogens, leaving its role in plant pathogens largely unexplored. Furthermore, while functional studies have predominantly examined 6mA methyltransferases, the roles of demethylases remain poorly understood.

Recently, DNA 6mA modification has been reported in an increasing number of eukaryotes, including *Caenorahabiditis elegans*(13), *Bursaphelenchus xylophilus*(14), *Drosophila*(15), *Chlamydomonas reinhardtii*(16), fungi(17, 18), plant(19–21), and mammal(22). In eukaryotes, 6mA methylation can occur in both symmetric and asymmetric contexts, while asymmetric 6mA methylation predominates in nematodes. Eukaryotic 6mA has been shown to be transmitted via a semiconservative mechanism(23) and incorporated into mammalian genomes by DNA polymerase(24, 25). Functionally, 6mA has been linked to disease onset(26–28), embryonic development(15), and environmental stress responses(20, 29). As a key epigenetic modulator, 6mA demethylases influences diverse biological processes, including eproduction(13), ovarian development(15), DNA replication and repair(30), vascular calcification(31), fatty acid metabolism in humans and mice(28), and neurodevelopment(32). The role of 6mA demethylase in pathogen infection has also been preliminary reported in *Mucor lusitanicus*(33). However, it is worth noting that several earlier studies reporting high 6mA levels in eukaryotes—especially mammals—were later shown to be confounded by bacterial DNA contamination(34). Thus, stringent control of bacterial contamination is critical for accurate detection of 6mA in eukaryotes.

Nevertheless, the mechanism by which 6mA regulates parasitism in eukaryotic pathogens—particularly the link between demethylases and virulence—remains unclear. Plant-parasitic nematodes (PPNs) cause devastating agricultural losses estimated to exceed $100 billion annually(35). Among them, root-knot nematodes (RKNs) are the most destructive(36), drawing increasing attention due to their broad host range and complex ploidy (37). Infective second-stage juveniles (J2s) penetrate host roots near the tip, migrate intercellularly, and establish giant cells within the vascular cylinder. These giant cells are extensively modified and serve as the sole nutrient source for nematodes throughout their lifecycle. A diverse suite of nematode effectors facilitate the formation and maintenance of giant cells(38). However, the epigenetic regulation of these effector genes remains largely unexplored. Previous studies of RKN epigenetics have mainly focused on histone modifications(39–41) and microRNA(42). Homology-based analysis of methyltransferases and demethylases suggested that the *Meloidogyne incognita* (*M. incognita*, Mi) lacks DNA 6mA modification machinery(43), raising doubts about the presence of 6mA in the RKN genomes. Alternatively, these enzymes may have diverged substantially from known homologs, warranting further investigation.

In this study, after rigorously eliminating bacterial DNA contamination, we investigated the presence of 6mA in 17 nematode isolates using LC-MS/MS and confirmed DNA 6mA modification in two diploids (*M. graminicola* and *M. hapla*), two allotriploids (*M. incognita* and *M. arenaria* 3n), and two allotetraploids (*M. arenaria* 4n and *M. javanica*) through SMRT-seq, N6-Methyladenine DNA immunoprecipitation sequencing (6mA-DIP-seq), and DNA Dot blot assays. We found that 6mA is enriched in actively transcribed genes and exhibits distinct distribution patterns across RKN species, particularly within transposable elements (TEs), where striking differences were observed between diploid and allopolyploid RKNs. In M. incognita, low-methylation-level sites were concentrated in the A2 subgenome and associated with genes critical for cellular function, such as G protein-coupled receptors (GPCR). Importantly, we identified two putative 6mA demethylases (MiNMAD-1 and MiNMAD-2) and pinpointed their catalytic sites through structural prediction and biochemical validation. Remarkably, host-induced gene silencing of *minmad-1* significantly enhanced resistance of transgenic host plants to three polyploid RKN species. Further analyses revealed that MiNMAD-1 globally regulates the expression pattern of numerous virulence genes. Collectively, our findings establish a direct link between epigenetic regulation and virulence in a eukaryotic pathogen.

## Results

### 6mA widespread in nematode genome

To investigate the presence of DNA 6mA in nematodes, we analyzed 17 isolates, including seven RKNs (*M. graminicola*, Mi, *M. arenaria* 3n, *M. arenaria* 4n, *M. javanica*, *M. enterolobii*, and *M. hapla*), four isolates of *Heterodera glycines*, three migratory PPNs (*Aphelenchus avenae*, *Ditylenchus destructor*, and *Bursaphelenchus xylophilus*), one animal-parasitic nematodes (*Haemonchus contortus*), and two free-living nematodes (*Caenorhabditis elegans* and *Caenorhabditis nigoni*). To minimize bacterial DNA contamination during liquid chromatography coupled with tandem mass spectrometry (LC-MS/MS) detection, we extracted DNA from sterilized eggs and verified the absence of bacterial contamination using 16S rRNA primers amplification(27F, 1492R) (see Method, *SI Appendix*, Fig. S1). Finally, a clear 6mA signal was detected in all tested species (Fig. 1*A*).

**Figure 1.**
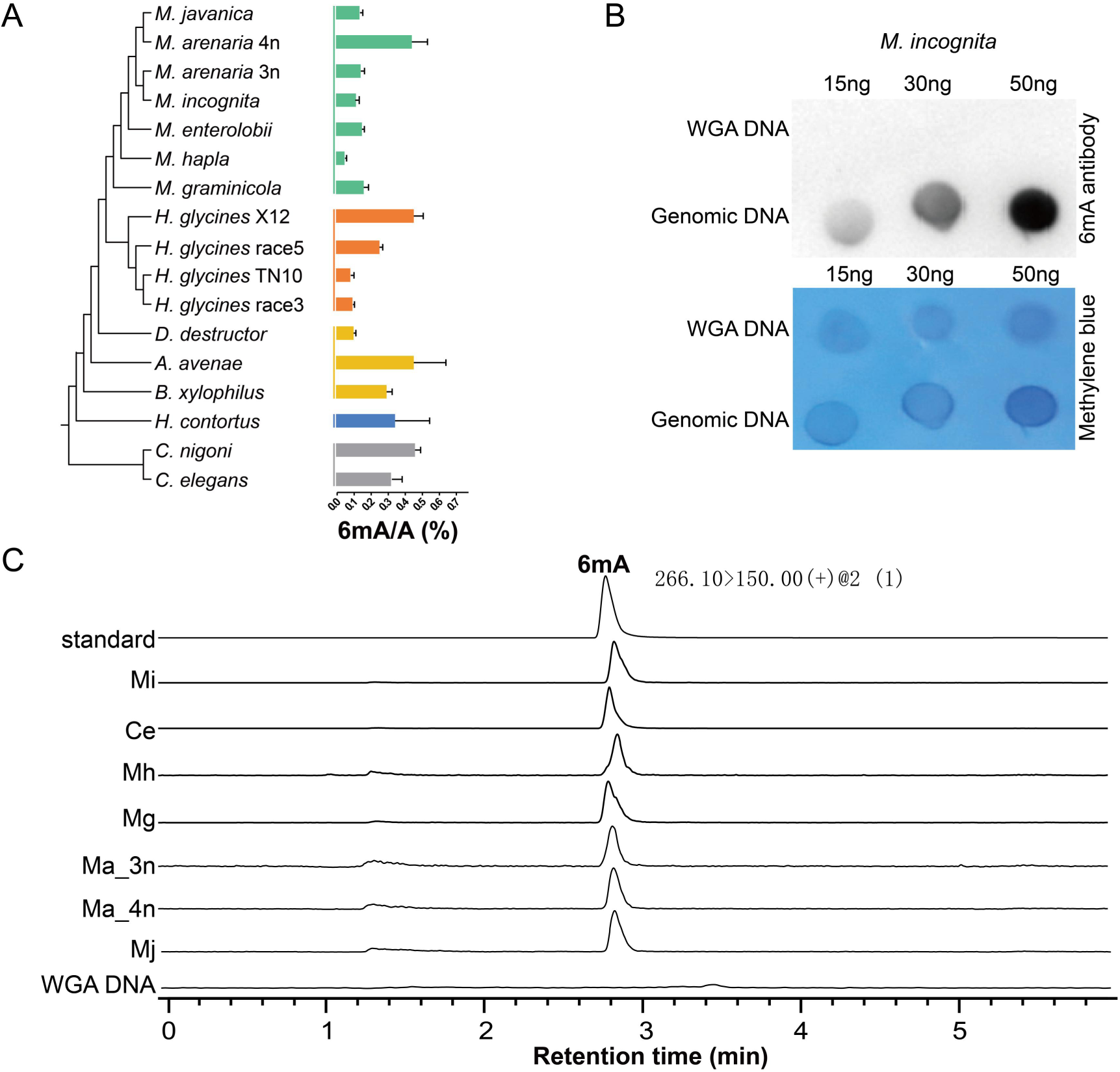
6mA DNA methylation is widespread in nematode. (*A*), 6mA levels identified by LC-MS/MS in different nematodes. The green bar plot means 7 species of RKNs, the orange bar plot means 4 isolates of cyst nematodes, the yellow bar plot means migratory nematodes, the blue bar plot means animal-parasitic nematode, and the gray bar plot means free-living nematode. Three biological replicates per species. (*B*), The DNA 6mA Dot blot of *M. incognita,* the figure below shows the membrane after antibody hybridization and chemiluminescence, and then methylene blue staining. Whole genome amplified (WGA) DNA of the *M. incognita* genome, generated using Phi29 polymerase, was used as a negative control. (*C*), LC-MS/MS for 6mA standard and 6mA levels in genomic DNA purified from *M. incognita*, *C. elegans*, *M. hapla* (Mh), *M. graminicola* (Mg), *M. arenaria* 3n (Ma 3n), *M. arenaria* 4n (Ma 4n), and *M. javanica* (Mj), respectively. The WGA DNA was used as a negative control. All DNAs were extracted from nematode eggs.

To further understand the role of 6mA methylation in the life history of PPNs, we focused on several species of RKNs (*SI Appendix*, Tables S1 and 2). PacBio sequencing is widely used to detect 6mA modifications at single-base resolution; however, false positives caused by bacterial contamination remain a major concern. To address this, we applied the 6mASCOPE approach (34) to quantify potential contamination by aligning PacBio sequel I reads to the NCBI nt database. The analysis showed that only 0.0004%-0.11% of reads were of bacterial origin (*SI Appendix*, Tables S3). To further validate these estimates, we generated HiFi sequencing data for Mi, Ma, and Mj and analyzed them with 6mASCOPE. The results revealed that the proportions of fungal and bacterial reads were 0.018%, 0.011%, and 0.011%, respectively (*SI Appendix*, Tables S4-6). The 6mASCOPE evaluation confirmed that the contamination rate in our data was extremely low. We then analyzed the 6mA content in our previously assembled genomes of *M. graminicola* (Mg), Mi, *M. arenaria* 3n (Ma 3n), *M. arenaria* 4n (Ma 4n), and *M. javanica* (44) using PacBio sequel I data, and the results were consistent with the LC-MS/MS measurements. In addition, we generated a chromosome-level genome assembly for the diploid *M. hapla* (Mh, *SI Appendix*, Tables S7) using Pacbio, Illumina, and Hi-C data (*SI Appendix*, Tables S8). The genome size of this assembly is consistent with the previously reported version (45) (53.1 Mb vs. 53.6 Mb), but demonstrates markedly higher quality, as reflected by a substantially improved scaffold N50 (2.8 Mb vs. 0.08 Mb) and a BUSCO completeness of 66.8%. Furthermore, we identified 6mA modifications at single-base resolution in Mh using PacBio sequel I data. To minimize false positives, we excluded 6mA sites with low sequencing coverage, using different minimum coverage thresholds (ranging from 15× to 50×) depending on the sequencing depth of each sample, as previously described(13, 17). After this filtering step, numerous 6mA sites were identified across the six genomes (*SI Appendix*, Tables S9). Subsequently, to further minimize false positives in 6mA analysis, we performed whole-genome amplification (WGA) on the genomic DNA of Mi, Ma, Mj, Mg, and Mh. To evaluate the reliability of 6mA site detection, we used WGA samples sequenced by PacBio HiFi as negative controls. The number of 6mA sites identified in these WGA samples, which should not contain true methylation, reflects the false-positive rate (*SI Appendix*, Tables S9 and S10). We observed that samples with minimal bacterial contamination—such as Ma 4n—had very few false-positive 6mA sites. In contrast, WGA samples with higher contamination levels yielded substantially more false-positive sites (*SI Appendix*, Tables S10 and S11). These findings highlight the importance of controlling for contamination when interpreting 6mA methylation data.

To further validate the presence of 6mA in RKNs genome, we first excluded potential RNA m6A contamination (*SI Appendix*, Fig. S2, see Method). We then performed 6mA Dot blot assays on egg genomic DNA from Mi, Ma 3n, Ma 4n, Mj, Mg, and Mh, which all displayed clear 6mA signals (Fig. 1*B*, *SI Appendix*, Fig. S3). In contrast, no signal was detected in the negative control, in which nematode DNA was subjected to whole-genome amplification (WGA) using phi29 polymerase (Fig. 1*B*) or 18S amplification (*SI Appendix*, Fig. S3). We next applied LC-MS/MS to discriminate between 1mA from 6mA in these six genomic DNAs, and only 6mA signals were detected (Fig. 1*C*). Additionally, we performed 6mA-DIP-seq on Mi genomic DNA and Mi-WGA DNA (*SI Appendix*, Fig. S4*A*). Pearson correlation analysis confirmed high reproducibility between replicates (*SI Appendix*, Fig. S4*B*). The results verified the presence of 6mA in the Mi genome, with 53% of the peaks overlapping those identified by SMRT sequencing (*SI Appendix*, Fig. S4*C*, *D*). By contrast, only a few peaks were identified in the Mi-WGA 6mA-DIP-seq data (*SI Appendix*, Tables S12). Collectively, these results provide strong evidence for the presence of 6mA in RKN genomes.

### 6mA distribution landscape in nematodes genome

DNA methylation site distribution has been reported to be associated with gene expression regulation (46). To comprehensively examine the regulatory effects of 6mA methylation on gene expression in RKNs and the free-living nematode *C. elegans* (Ce, data download form GEO database, GSE66504), we categorized all 6mA-modified genes according to whether the sites were located in promoter, gene body, or distal intergenic regions. Most 6mA sites were found in gene bodies and promoters, with Ce showing the highest proportion of 6mA distribution in promoters (59.93%) (Fig. 2*A*; *SI Appendix*, Fig. S5). We then extracted 6mA sites from each of these regions and analyzed the 8-bp sequence flanking the modification sites for motif enrichment. Within a given species, the motifs of 6mA sites in promoters, gene bodies, and intergenic regions were largely similar (*SI Appendix*, Fig. S6). By contrast, cross-species comparisons revealed that motifs were highly conserved among RKNs but markedly different between RKNs and Ce (*SI Appendix*, Fig. S6). Specifically, the core motif in RKNs was GAG, whereas in Ce it was ApT, consistent with previous observations in fungi. This finding suggest that 6mA may regulate gene expression through distinct contexts in RKNs and Ce.

**Figure 2.**
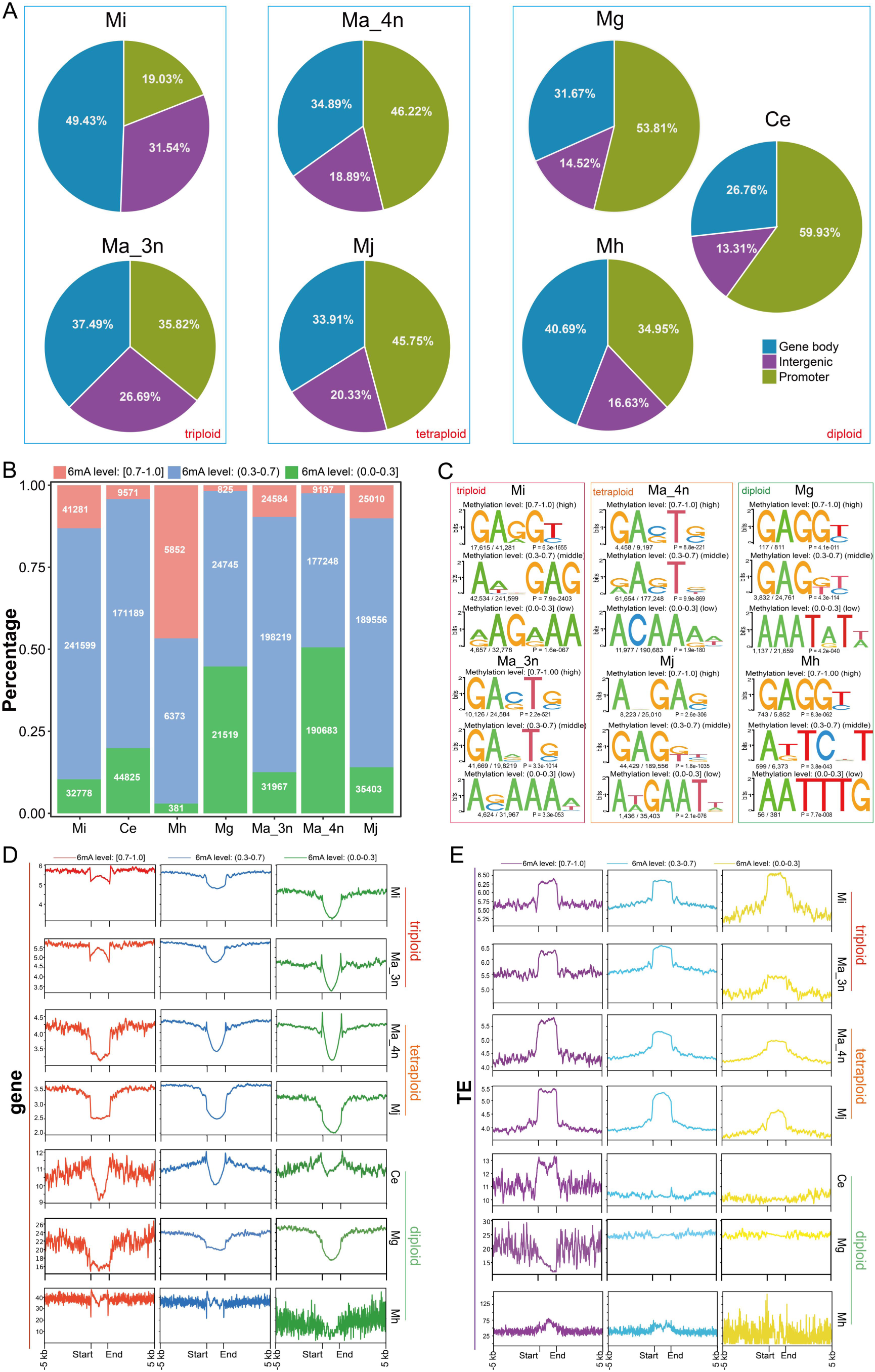
The distribution landscape of 6mA in RKNs genome. (*A*), The distribution ratio of 6mA in three nematodes promoter, gene body, and intergenic. *M. incognita* (Mi), *M. arenaria* 3n (Ma 3n), *M. arenaria* 4n (Ma 4n), *M. javanica* (Mj), *M. graminicola* (Mg), *M. hapla* (Mh), and *C. elegans*. The 1 kb region before the start codon is defined as the promoter. (*B*), Percentage of the different modification level categories of 6mA in Mi, Ma 3n, Ma 4n, Mj, Mg, and Mh, respectively. (*C*), The motif of different methylation categories of Mi, Ma 3n, Ma 4n, Mj, Mg, and Mh. (*D*) The distribution density of 6mA on gene regions within 100bp genome window (Mi, Ma 3n, Ma 4n, Mj, Ce, Mg, and Mh). (*E*) The distribution density of 6mA on transposable element (TE) region within 100bp genome window. *p* values were calculated by two-sided Wilcoxon rank sum tests. The y-axis represents the average number of 6mA per 100 bp genome window.

To investigate the functional implications of different 6mA methylation levels, we classified 6mA sites into three categories: low (at a specific locus, <30% of reads methylated across all sequencing data), moderate (30%-70%), and high (70%-100%). Moderate-methylation sites were the most abundant in Ce, Mi, Ma 3n, and Mj, accounting for ∼76% of all 6mA sites. In Ma 4n, Mg, and Mh, approximately 50% of the sites showed moderate methylation, whereas only a small fraction exhibited high or low methylation (Fig. 2*B*). Motif analysis of the 8-bp flanking sequences revealed three distinct motifs corresponding to the methylation categories in each nematode (Fig. 2*C*). Among them, the GAG motif was most enriched at high-methylation sites in the four RKNs (Mi, Mj, Mg, and Mh), consistent with previous findings in *C. elegans*(13) and rice(19, 20). By contrast, motifs associated with moderate and low methylation levels varied considerably among species (Fig. 2*C*), suggesting that different levels of 6mA methylation may be regulated by distinct modifying enzymes and may serve diverse biological functions.

Subsequently, we analyzed the distribution density of 6mA sites in gene regions and transposable element (TE) regions across these nematode genomes. We found that the gene regions generally exhibited a low density of 6mA in all nematodes except Mh (Fig. 2*D*), a pattern reminiscent of the distribution characteristics of 5mC(19). By contrast, TE regions showed significant differences in the distribution density of 6mA sites between polyploidy and the diploid species (Ce, Mg, and Mh) (Fig. 2*E*). These observations suggest potential lineage-specific patterns of 6mA–TE distribution, which may warrant further investigation in the context of polyploidization.

To further investigate the distribution patterns of 6mA in TE regions of polyploid and diploid nematodes, we examined 6mA enrichment across different TE classes. In polyploid nematodes, 6mA was densely distributed in DNA transposons, MITEs, and LTRs, but showed a low density in low-complexity regions (*SI Appendix*, Fig. S7-10). We noticed that both the content and composition of TE differed significantly between polyploid and diploid RKNs (*SI Appendix*, Tables S13). In polyploid RKNs, TEs accounted for 28%-33% of the genome, with 8%-10% classified as Low-complexity elements (including simple repeat). While in diploid RKNs, only 8%-10% of the genome consisted of TEs, of which 41%-81% were low-complexity elements. However, differences in TE composition alone cannot explain the pronounced discrepancy in 6mA abundance between polyploid and diploid nematodes. For instance, although diploid nematodes harbor a substantial proportion of DNA transposons, MITE-type Telements in these species still lacked the high-density 6mA modification observed in polyploids (*SI Appendix*, Fig. S11-13). For example, although *C. elegans* exhibits a TE composition similar to that of polyploid RKNs (*SI Appendix*, Tables S13), its 6mA distribution across TEs resembled that of diploid nematodes. To verify the authenticity of our observations, we repeated the analysis using false-positive 6mA sites predicted from WGA sequencing data. These false-positive sites showed neither increased density in the TE regions nor a decreased density in gene regions (*SI Appendix*, Fig. S14), but instead appeared randomly distributed across the genome. To further confirm the enrichment of 6mA in TE regions, we compared 6mA-DIP-seq peaks with the annotated TE regions in Mi. The results revealed that 74.8% of merged 6mA-DIP-seq peaks overlapped with the TE regions, whereas TEs comprise only 28.18% of the Mi genome. Together, these results indicate that the high-density 6mA feature of TEs in polyploid RKNs is not directly associated with TE type. Instead, it may reflect broader genomic processes, such as TE activity or mechanisms contributing to genome stability.

### 6mA regulates gene expression of RKNs

To a<u>sses</u>s the impact of 6mA methylation on gene expression, we examined the relationship between methylation status and gtranscrip levels from multiple perspectives. First, we compared the expression levels of 6mA-methylated and unmethylated genes in the egg stage (same stage as collecting SMRT sequencing data) across six RKN species. The results showed that in nearly all RKNs, 6mA-methylated genes exhibited significantly higher expression levels than unmethylated genes, whereas the opposite trend was observed in Mh (Fig. 3*A*). In polyploid RKNs, unmethylated genes were predominantly expressed at low levels (TPM<1), while a larger proportion of methylated genes displayed higher expression levels (TPM>5) (Fig. 3*B*). By contrast, in diploid RKNs, most genes—regardless of methylation status—showed expression levels exceeding TPM>5. In the free-living nematode Ce, however, the majority of genes exhibited low expression (TPM < 1) (Fig. 3*B*). Collectively, these findings suggest that 6mA methylation is associated with transcriptional activation in polyploid species, while no such trend is evident in diploid nematodes.

**Figure 3.**
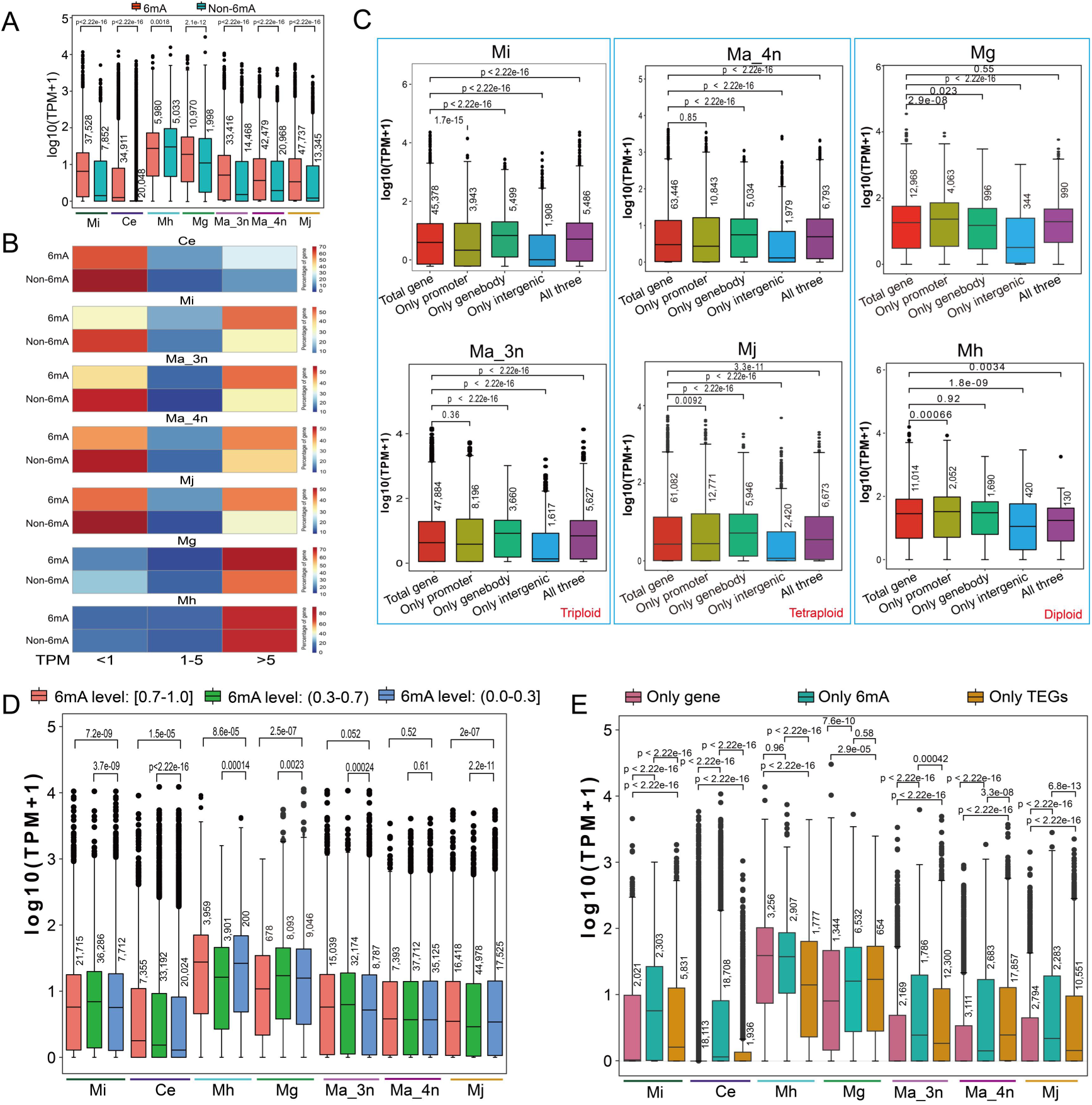
6mA influences the genes expression level of nematodes. (*A*) The comparison of TPM value between 6mA-methylated (6mA) and unmethylated (Non-6mA) in Ce, Mi, Ma, Mj, Mg, and Mh. The gene with 6mA modification in the promoter, gene body, or intergenic region was defined as a 6mA-modified gene; otherwise, it was classified as an unmethylated gene. (*B*) The percentage of 6mA-methylated (6mA) and unmethylated (Non-6mA) genes at a given TPM level (<1, 1–5, and >5). (*C*) The comparison of expression levels between the genes modified by 6mA at different positions and total genes. (*D*) The comparison of genes expression level of different 6mA modification level categories in Mi, Ma 3n, Ma 4n, Mj, Ce, Mg, and Mh. (*E*) Gene expression in Mi, Mh, Mg, Ma 3n, Ma 4n, and Mj categorized into three groups: genes modified only by 6mA without nearby TE (“only 6mA”), genes associated only with nearby TE but lacking 6mA (“only TE”), and genes with neither 6mA modification nor nearby TE (“only gene”). In box plots, the central line represents the median, the box represents the 25% and 75% percentiles, and the whiskers represent 1.5 times the interquartile range beyond the box. N numbers were shown upon the box. *p* values were calculated by two-sided Wilcoxon rank sum tests.

Subsequently, we performed motif enrichment analysis of 6mA sites in genes with TPM<1, 1<TPM<5, and TPM>5. The results showed that within the same species, genes across different expression levels exhibited similar motif enrichment patterns, and motifs among RKNs were relatively conserved, characterized by a core GAG motif (*SI Appendix*, Fig. S15). In contrast, a marked divergence was observed between Ce and RKNs, with the core motif in Ce being ApT (*SI Appendix*, Fig. S15). Notably, the motif patterns associated with genes of different expression levels are similar to those observed for 6mA sites located in different genomic regions, such as promoters, gene bodies, and distal intergenic regions (*SI Appendix*, Fig. S6).

Second, we examined genes modified by 6mA at only one of the three positions (promoter, gene body, or intergenic) as well as those simultaneously modified at all three positions (*SI Appendix*, Fig. S16) to explore expression differences among these categories. The analysis revealed that in polyploid RKNs (Mi, Ma 3n, Ma 4n, and Mj), genes methylated in the gene body showed higher expression levels than the average of total genes (Fig. 3*C*). In contrast, in diploid RKNs, promoter-methylated genes exhibited higher expression levels than the average of total genes (Fig. 3*C*). Moreover, in Ce, both promoter- and gene body-methylated genes were expressed at higher levels than the average of total genes (*SI Appendix*, Fig. S17), which differs from the patterns in RKNs. These results suggest that 6mA exerts distinct regulatory effects on gene expression in polyploid versus diploid RKNs, which may reflect differences in ploidy level and reproductive strategy.

Third, we analyzed the expression levels of genes associated with different 6mA methylation levels (low, moderate, and high). The results showed that in Mi, Ma 3n, and Mg, genes with moderate methylation exhibited the highest expression levels, whereas in Ce and Mh, genes with high methylation displayed the highest levels (Fig. 3*D*). These finding indicate that the impact on gene expression differs among nematodes species.

Finally, we compared the expression levels of transposable element-related genes (TEGs) and 6mA-marked genes, as previously described(19). Genes unmodified by either 6mA or TE shows the lowest expression levels in all six nematodes except Mh. The triploid RKNs (Mi and Ma 3n) displayed similar patterns, whereas diploid and tetraploid RKNs did not (Fig. 3*E*). In Ma 3n, Ma 4n, Mj, Mg, and Ce, 6mA-modified genes with nearby TE showed higher expression than those without, while the opposite trend was observed in Mi and Mh (*SI Appendix*, Fig. S18). Moreover, 6mA-marked TEGs consistently exhibited significantly higher expression than unmarked TEGs (*SI Appendix*, Fig. S18). These findings indicate that 6mA exerts context-dependent effects on gene expression.

### Low-methylation-level 6mA sites play a crucial role in Mi

We examined the genomic distribution of 6mA sites at different methylation levels in RKNs (*SI Appendix*, Fig. S19*A*). Overall, sites of all three methylation levels were evenly distributed across chromosome, except for low-methylation-level sites in Mi, which showed a distinct enrichment (green circle in *SI Appendix*, Fig. S19*A*). Specifically, 72% of high-density-regions (HDRs, >60 sites per 50 kb genome window) of low-methylation-level 6mA were located in the A_2_ subgenome. We then extracted genes from these HDRs in the A_2_ subgenome, along with orthologous regions in A_1_, and B_1_, to compare their expression level in egg stage. Notably, the genes from A_2_ consistently exhibited higher expression than those from A1 and B1 (*SI Appendix*, Fig. S20). Moreover, these A_2_ genes maintained higher expression not only in egg stage (the stage used for 6mA profiling), but also across other developmental stages of Mi (*SI Appendix*, Fig. S19*B*), suggesting a subgenome-specific regulatory role of 6mA.

To explore the functional differences of the genes associated with HDRs of low-methylation-level 6mA sites between A_2_ and the A_1_/B_1_ subgenome, we compared their enriched Pfam domain. The results showed that the genes from A_2_ subgenome were enriched in GPCR (*SI Appendix*, Fig. S19*C*), consistent with previous GO enrichment results of the genes with high methylation density in human(22). The GPCR superfamily represents one of the largest chemoreceptor families; in nematodes, the absence of GPCR renders them “blind” and “deaf” (47). GPCRs are also critical for fungal infection(48, 49) and may play essential roles in host recognition and effector secretion in RKNs(50). Furthermore, GO analysis revealed that A_2_-associated genes were predominatly enriched in key molecule function, such as pharyngeal gland morphogenesis in Mi (*SI Appendix*, Fig. S19*D*). These finding suggest that low-methylation 6mA–modified genes in the A_2_ subgenome participate in vital biological processes and exert a stronger regulatory influence in A_2_ than in A_1_ or B_1_ subgenome.

### MiNMAD-1/2 are involved in 6mA demethylation

A previous study identified three 6mA methyltransferases from Mi, but the corresponding demethylase remained unknown (43). To test whether Mi possesses 6mA demethylase activity, we synthesized a biotin-labeled oligo DNA containing 6mA and measured its methylation level after incubation with a suspension of disrupted Mi eggs. Compared with the heat-inactivated egg suspension control and the BSA control, the 6mA content in the treatment group was significantly reduced (*SI Appendix*, Fig. S21*A*), indicating the presence of one or more putative 6mA demethylases in Mi.

To identify putative demethylase genes in Mi, we collected six genes containing 2OG-FeII_Oxy_2 domain based on Pfam annotation, together with three genes showing the highest sequence similarity to the *C. elegans* demethylase gene *nmad-1*. Phylogenetic analysis revealed that these nine putative demethylase genes (including three alleles) clustered into the *alkbh1*, *alkbh6,* and *alkbh7* families (Fig. 4*A*; *SI Appendix*, Fig. S22). To assess whether these genes function as 6mA demethylases, we measured genomic DNA 6mA levels after RNAi-mediated knockdown of each gene in Mi eggs using LC-MS/MS. The results showed that knockdown of *Mi_14752.1* and *Mi_00010.1* significantly increased 6mA levels, whereas knockdown of *Mi_23849.1* had no effect (Fig. 4*B*; SI Appendix, Fig. S21*B*).

**Figure 4.**
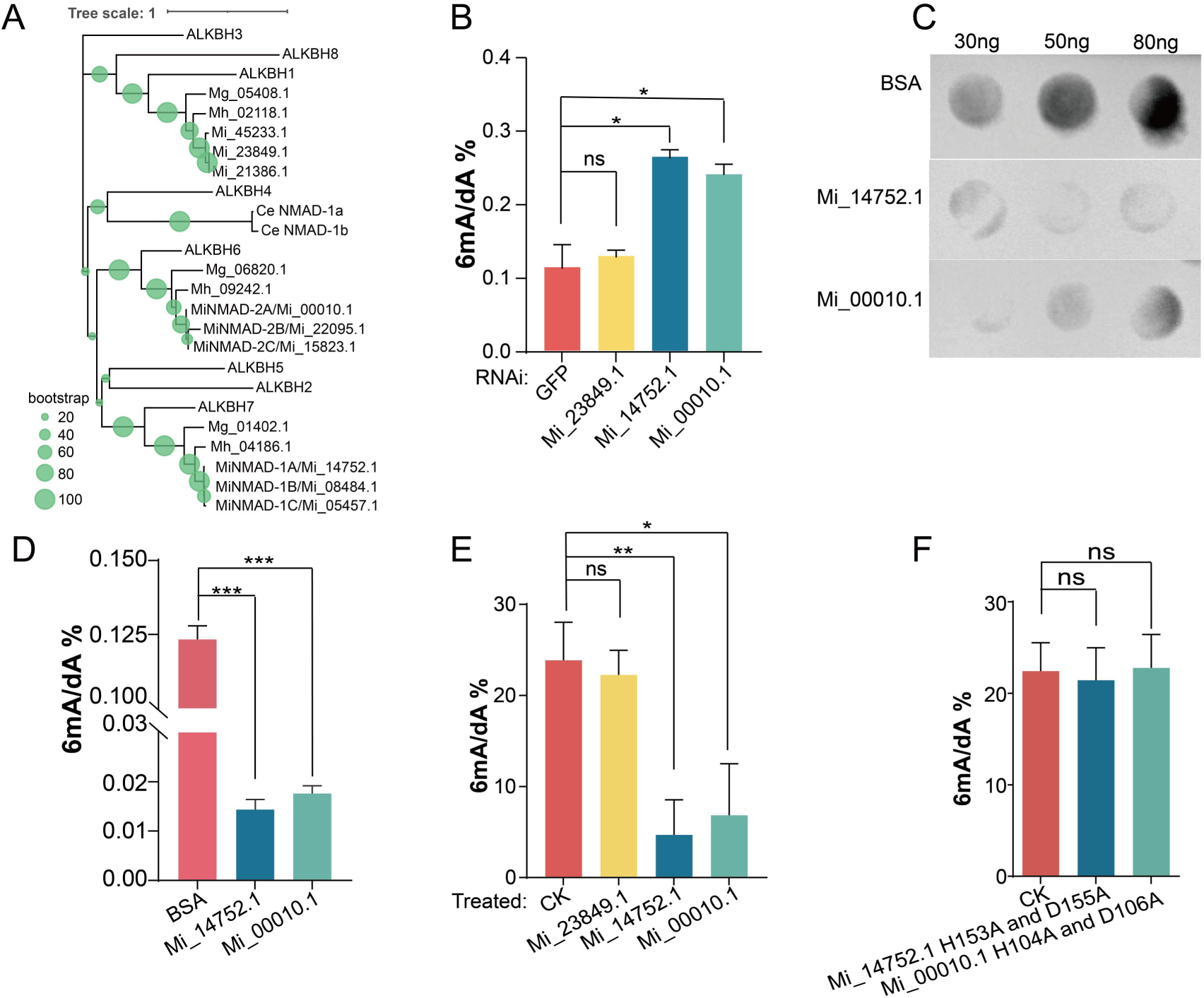
MiNMAD-1/2 are involved in 6mA demethylation in *M. incognita*. (*A*), The maximum-likelihood tree is constructed from the protein sequence of RKNs, *C. elegans* demethylase, and human ALKBH family members, bootstraps are shown in green dot. (*B*), Knockdown of 3 ALKBH family members present a significant increase of 6mA level in two of them (Mi_14752.1, Mi_00010.1). (*C*), In vitro demethylation assay showing that purified Mi_14752.1 and Mi_00010.1 proteins catalyze the removal of DNA methylation from Mi WHF4-1 egg genomic DNA, with BSA included as a negative control. (*D*), The 6mA contents of (*C*) were determined by LC-MS/MS. (*E*), LC-MSMS detected the oligo DNA with 6mA modified (G**A**G) was demethylated by purified MiNMAD-1/2 proteins *in vitro*. (*F*), Mutation of the catalytic domain of Mi_14752.1 and Mi_00010.1 abrogates the ability of them to demethylate 6mA premethylated oligos. *p* values were calculated by two-tailed unpaired Student’s *t*-test. One star * means *p* < 0.05, two means *p* < 0.01, three means *p* < 0.001.

We purified these three candidate proteins and tested their demethylation activity *in vitro.* Both Mi_14752.1 and Mi_00010.1 were able to demethylate Mi genomic DNA (Fig. 4*C*, *D*; *SI Appendix,* Fig. S23). To further validate their activity, we treated synthetic oligo DNA carrying 6mA modifications with the purified proteins, and again observed significant demethylation activity for Mi_14752.1 and Mi_00010.1 (Fig. 4*E*). In addition, comparative sequence analysis revealed that although demethylases from different species show substantial divergence at the overall sequence level, their catalytic active sites are highly conserved (*SI Appendix*, Fig. S24*A*). We predicted the 3D structure of Mi_14752.1, Mi_00010.1, Mi_23849.1, and Ce_NMAD-1 using Alphafold2(51). Structural comparison revealed that Mi_14752.1 closely resembles human ALKBH7, with a root-mean-square deviations (RMSD) of 0.067 Å (*SI Appendix*, Fig. S24*B*). Moreover, the catalytic sites of Ce_NMAD-1 overlap with those of Mi_14752.1 and Mi_00010.1 but not with Mi_23849.1 (*SI Appendix*, Fig. S24*C*-*E*). The RMSD values between Ce_NMAD-1 and Mi_14752.1, Mi_00010.1, and Mi_23849.1 were 2.392, 1.066, and 12.625 Å, respectively (SI Appendix, Fig. S24*C*-*E*). To determine whether the demethylase activity is intrinsic to Mi_14752.1 and Mi_00010.1, we introduced point mutations (H153A and D155A in Mi_14752.1; H104A and D106A in Mi_00010.1). These mutations completely abolition their demethylation activity (Fig. 4*F*). Accordingly, we designated Mi_14752.1 as Mi N6-methyladenine demethylase 1 (MiNMAD-1) and Mi_00010.1 as MiNMAD-2, reflecting their newly identified demethylation functions.

### Host-induced gene silencing (HIGS) of *minmad-1*/*2* reduces infectivity of MIG

Considering that most stages of the RKN life cycle occur within plants, we generated transgenic tobacco lines expressing dsRNA targeting *minmad-1*/*2, Mi_23849.1* (to verify its demethylase activity during infection), and *gfp* as a control, to investigate the regulatory role of 6mA in parasitic stage of the *M. incognita* group (MIG, including Mi, Mj, and Ma). In three-week pot experiments, *minmad-1*/*2*-HIGS tobacco plants exhibited significantly enhanced resistance to Mi compared with *gfp*-HIGS and CK (wild-type) controls, whereas *Mi_23849.1*-HIGS plants showed no significant difference (Fig. 5*A*). The eight-week pot experiment showed that the gall number in the high-expression line of *minmad-1/2*-HIGS tobacco was reduced by approximately tenfold compared with that in *gfp-*HIGS and CK plants (Fig. 5*B*; *SI Appendix*, Fig. S25*A*, *B*). As anticipated, Mi genomic DNA exhibited a significant increase in 6mA levels upon infestation of *minmad-1/2*-HIGS tobacco relative to the control (Fig. 5C), confirming the demethylase activity of MiNMAD-1/2 in Mi. Subsequently, according to galls number, we divided the nematode-infested *minmad-1*/*2-*, and *Mi_23849.1-*HIGS tobacco plants into three groups including high-, medium-, low-galls groups (*SI Appendix*, Fig. S26*A*-*C*). After confirming the absence of bacterial contamination (see Methods), DNA from females and eggs of the three groups was extracted for 6mA quantification. The results showed a negative correlation between 6mA content and gall number (Fig. 5*D*), indicating that demethylase silencing reduces the virulence of Mi. Additionally, the 6mA level of females was much higher than that of eggs (Fig. 5*D*), indicating a stage-specific difference in 6mA levels during development. To further explore the resistance conferred by demethylase silencing in different hosts, we generated *minmad-1*-HIGS tomato lines. Pot experiments demonstrated that RNAi targeting the demethylase enhanced tomato resistance to Mi (*SI Appendix*, Fig. S27*A*-*C*), and the 6mA level in Mi feeding on transgenic plants increased significantly (*SI Appendix*, Fig. S27*D*). These results highlight the feasibility of using transgenic plants expressing Mi demethylase dsRNA to confer resistance against Mi.

**Figure 5.**
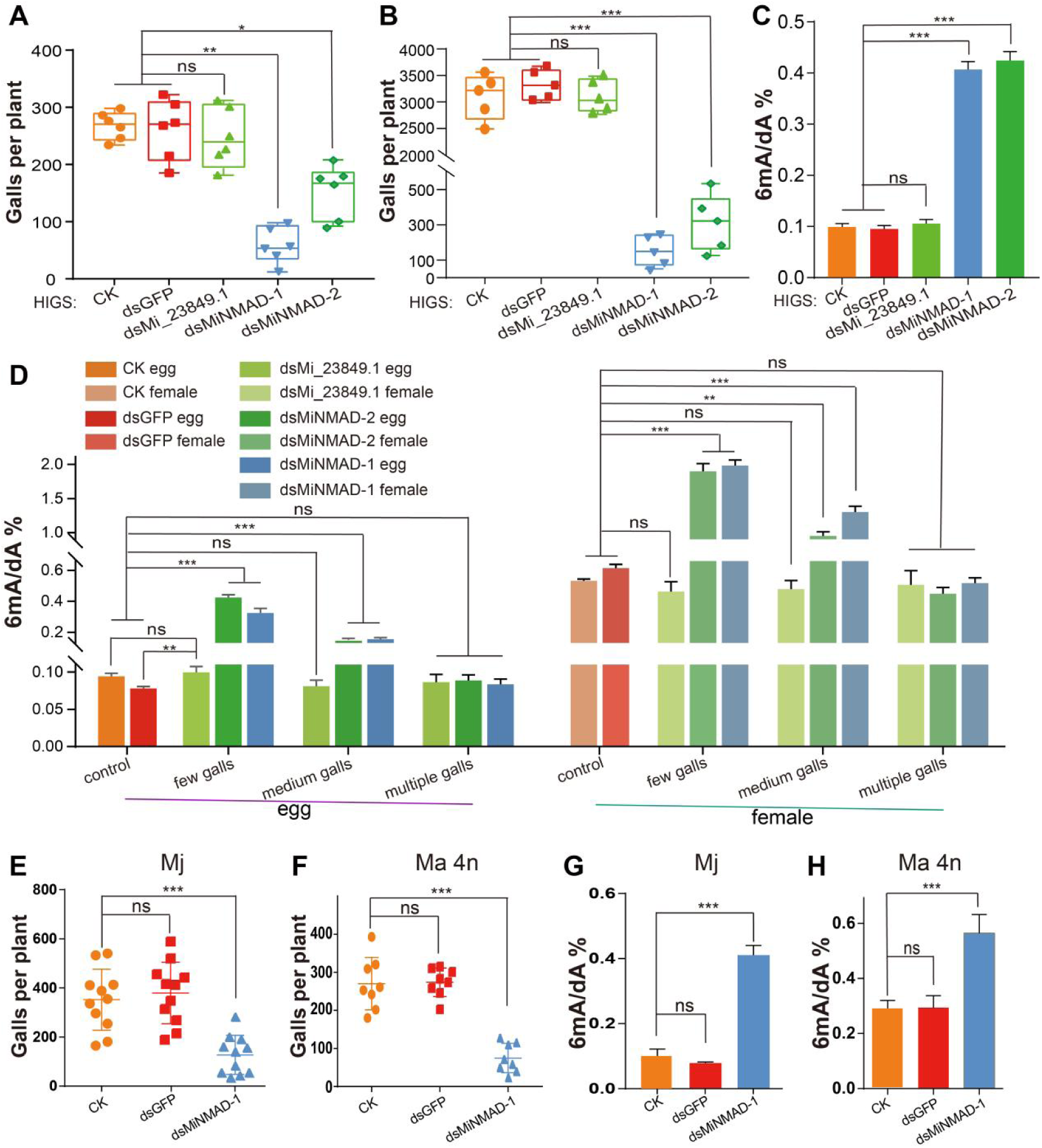
Host-induced gene silencing of *minmad-1* reduces the infectivity of polyploid RKNs. (*A*) Three weeks pot experiment of transgenic plants show a resistance of Mi for *minmad-1/2* RNAi, but no difference for *Mi_23849.1* RNAi. (*B*) Eight weeks pot experiment show an extremely significant resistance of expression *MiNMAD-1/2* dsRNA transgenic tobacco to Mi. (*C*) The 6mA level of Mi is significantly increased after infection of transgenic tobacco expression *minmad-1/2* dsRNA. (*D*) The number of galls was proportional to the methylation content in the corresponding Mi, and the level of 6mA in females was much higher than in eggs. (*E*) The number of galls after 1000 *M. javanica* (Mj) J2 infection two months of transgenic tobacco expressing *minmad-1* dsRNA. (*F*) The number of root-knots after 1000 M. arenaria 4n (Ma 4n) J2 infection of two months of transgenic tobacco expressing minmad-1 dsRNA. (*G*) The genomic DNA 6mA content of the Mj after infecting the *minmad-1* transgenic tobaccos was significantly higher than that of the control. (*H*) The genomic DNA 6mA content of the Ma 4n after infecting the *minmad-1* transgenic tobaccos was significantly higher than that of the control. *p* values were calculated by two-tailed unpaired Student’s *t*-test. One star * means *p* < 0.05, two means *p* < 0.01, three means *p* < 0.001.

Additionally, *minmad-1* sequences are completely conserved among polyploid RKNs (*SI Appendix*, Fig. S28), suggesting that the transgenic plants expressing Mi demethylase dsRNA could potentially confer resistance to multiple polyploid RKNs. To test this hypothesis, we evaluated the infection ability of other polyploid RKNs on transgenic tobacco. The results showed that *minmad-1*-HIGS tobacco plants displayed significant resistance to the tetraploid RKNs Mj and Ma 4n compared with CK and *gfp*-HIGS controls (Fig. 5*E*, *F*; *SI Appendix*, Fig. S29*A*, *B*). Consistently, the 6mA levels of genomic DNA from Mj and Ma 4n were significantly increased in *minmad-1-*HIGS group (Fig. 5*G*, *H*). Finally, to assess the role of demethylases in diploid RKNs, we generated transgenic rice expressing Mg putative demethylase dsRNA. However, these plants did not exhibit resistance to Mg, despite the significant elevation of 6mA levels in nematodes feeding on them (*SI Appendix*, Fig. S29*C*-*F*). Together, these results suggest that demethylases regulate the virulence of polyploid RKNs, whereas their functional impact on diploid RKNs remains to be further explored.

### 6mA regulates the expression pattern of the Mi virulence gene

To further investigate the regulatory effect of the 6mA demethylase on Mi, we grew 80 *minmad-1*-HIGS tobacco plants inoculated with Mi. Based on gall formation, we divided the 80 tobacco plants into three categories: high-gall plants (groups 1–2, with numerous galls but fewer than those in the control), medium-gall plants (groups 3–6), and low-gall plants (groups 7–8). Infective third/fourth-stage juveniles (J3/J4) were then collected from these three categories for RNA-seq analysis (*SI Appendix*, Tables S14). We identified 7,198, 8,978, and 8,660 DEGs in high-, medium-, and low-gall plants, respectively (*SI Appendix*, Fig. S30). Among them, 4,667 DEGs were shared across all three groups (*SI Appendix*, Fig. S31*A*) and exhibited two distinct expression patterns (*SI Appendix*, Fig. S31*B*). The numbers of DEGs unique to the high-, medium-, and low-gall plants were 832, 1685, and 1491, respectively (*SI Appendix*, Fig. S31*A*). Functional enrichment analysis revealed that a large proportion of downregulated DEGs unique to the medium-gall plants were ribosome genes (*SI Appendix*, Fig. S31*C*), suggesting that demethylase activity may influence ribosomal function, thereby modulating mRNA translation efficiency and ultimately reducing Mi infectivity.

To explore the primary cause of the reduced infectivity of Mi in *minmad-1-*HIGS tobacco plants, we focused on the nematode’s virulence gene (effectors), which constitute its key molecular weapons for host parasitism. From the 4,667 DEGs shared by the three groups (high-, medium-, and low-gall plants), we identified 520 putative effector genes (*SI Appendix*, Fig. S31*A*). The putative effector genes exhibited consistently lower expression levels across all three gall groups compared with the control (*SI Appendix*, Fig. S31*D*), with the low-gall group showing the most pronounced downregulation (*SI Appendix*, Fig. S31*E*). These results indicate a negative correlation between transgenic plant resistance and the expression levels of Mi effectors.

Nematode invasion requires diverse effectors, and their proper expression is essential for host infestation (52). Expression profiling revealed that all putative secreted protein genes clustered into 11 groups, with RNAi plants displaying an expression pattern nearly opposite to the control (Fig. 6*A*). These results indicate that demethylases globally alter effector expression, thereby reducing Mi infectivity. To further examine the regulatory effect of demethylase on effectors, we measured the expression of several reported nematode effectors in *minmad-1-*HIGS and *gfp-*HIGS control tobacco plants by qRT-PCR. The results showed that, compared with the *gfp-*HIGS control, the expression levels of Mi-XYl1, Mi-CRT, Mi8D05, MiPFN3, 16D10, MiSGCR1, and MiIDL1 in female stage were markedly reduced, by approximately 22-fold (log₂FC = –4.46), 126-fold (log₂FC = –6.98), 10-fold (log₂FC = –3.32), 28-fold (log₂FC = –4.81), 27-fold (log₂FC = –4.75), 5-fold (log₂FC = –2.32), and 13-fold (log₂FC = –3.70), respectively (Fig. 6*B*; *SI Appendix*, Fig. S32; *SI Appendix*, Tables S15). To determine whether effector reduction also occurred at the protein level, we generated a specific antibody against Mi-CRT. Western blot analysis revealed that *minmad-1* RNAi significantly decreased Mi-CRT protein accumulation compared with the gfp-HIGS and CK controls (Fig. 6*C*). Moreover, Mi-CRT expression was down-regulated in both female and egg stages (Fig. 6*D*), suggesting that 6mA DNA methylation may affect effector expression primarily at the transcriptional level, thereby leading to reduced protein abundance.

**Figure 6.**
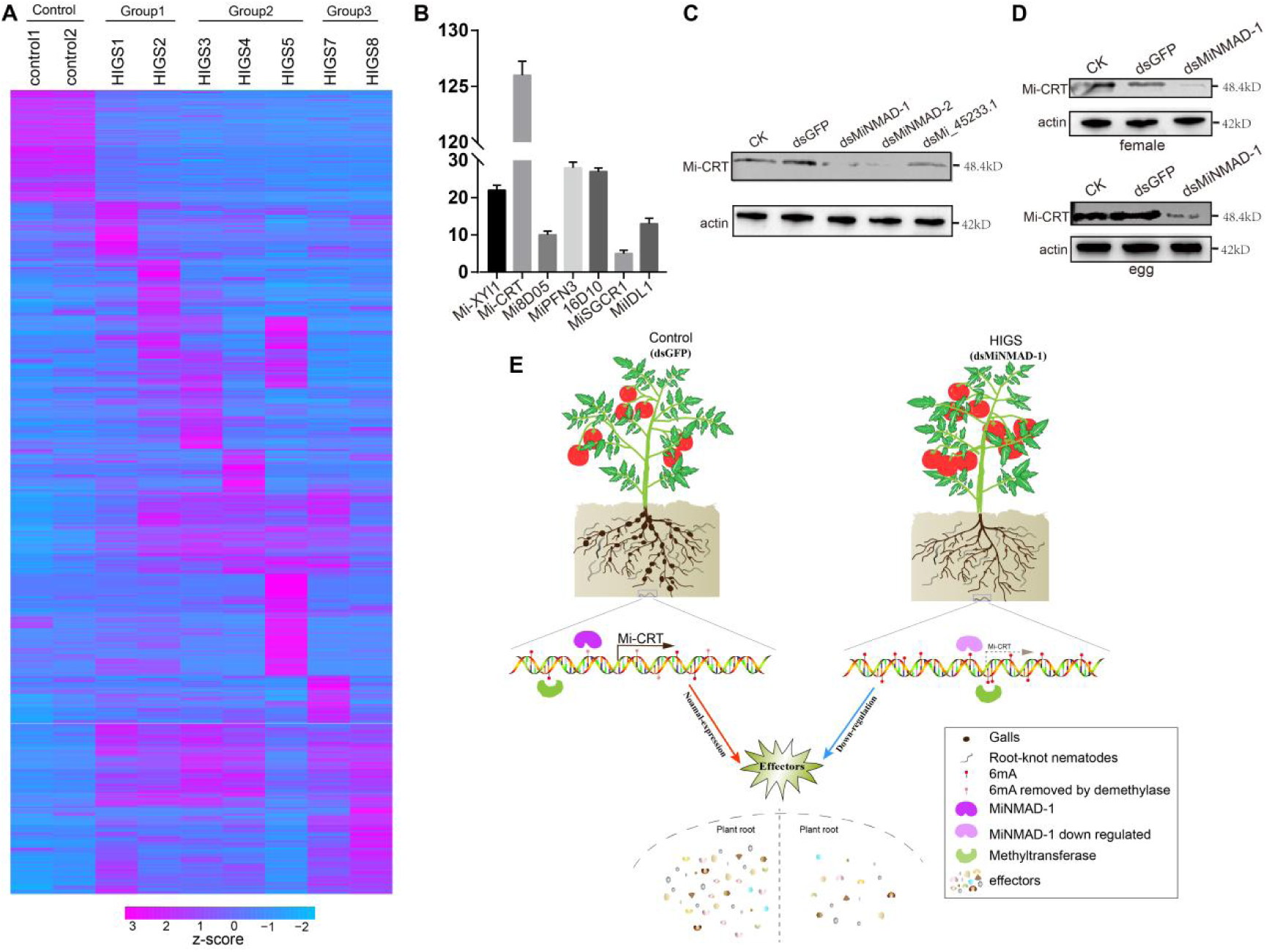
6mA regulation of the expression module of *M. incognita* effector. (*A*) The Expression patterns of all secreted proteins at Mi J3J4 stage in dsGFP control group and *minmad-1/2* RNAi group. Group1: multiple-galls. Group2: medium-galls. Group3: low-galls. (*B*) Relative fold down-regulation of some validated effectors after HIGS of *minmad-1*, assay was performed by qRT-PCR in Mi female. (*C*) Western blot of the highest down-regulate effector show that Mi-CRT was also significantly down-regulated at the protein level after *minmad-1/2* RNAi. (*D*) Mi-CRT protein level down-regulate both at egg and female stage. (*E*) Schematic illustration of the regulation of Mi virulence factor expression patterns by 6mA to affect the ability of nematode infectivity.

We generated a specific antibody against the demethylase and performed ChIP-seq analysis on infective J2s to investigate whether the enzyme directly binds to and regulates effectors in polyploid RKNs (*SI Appendix*, Fig. S33*A*). The results revealed that 40.4%–51.7% of the candidate effector genes were bound by the demethylase (*SI Appendix*, Fig. S33*B*-*D*). We next performed 6mA-DIP-seq on Mi J2s to further assess whether 6mA directly marks the candidate effector genes. Approximately 40% of the candidate effector genes carried 6mA modifications, and notably, 47.4% of these overlapped with demethylase ChIP-seq targets (*SI Appendix*, Fig. S34). Together, these findings demonstrate that the enzyme directly acts on effector genes and may influence their transcriptional regulation.

Together, these results support a model in which 6mA methyltransferases and demethylases dynamically regulate effector methylation in a spatiotemporal manner, thereby shaping their expression across developmental stages (Fig. 6*E*). Such regulation likely represents a major epigenetic mechanism underlying the parasitic ability of RKNs. Importantly, whether 6mA also acts through transcription factors to exert broader control over effector networks remains to be determined, and will be an important direction for future research.

## Discussion

In recent years, the existence of 6mA in eukaryotes has been debated, largely due to concerns about bacterial DNA contamination. To address this, we implemented multiple strategies to minimize false positives: (i) DNA was extracted from axenic eggs collected after nematode infection of axenic tobacco cultures; (ii) All tissues (egg, J3J4, female) were surface-sterilized with sodium hypochlorite and plated on LB and PDA media to confirm the absence of bacteria; (iii) 16S rRNA amplification showed no bacterial DNA bands; (iv) SMRT-seq reads were first aligned to nematode genomes and sites with ≥15× coverage were retained; (v) false positives were further excluded by sequencing WGA samples using PacBio and 6mA-DIP-seq; (vi) HiFi data analyzed by 6mASCOPE indicated bacterial/fungal contamination levels as low as ∼0.01%; and (vii) LC-MS/MS distinguished DNA-derived 6mA from RNA-derived m6A. These results confirm that contamination in our datasets is negligible, strengthening the reliability of our findings. Nevertheless, when studying other nematodes—especially migratory parasites that feed on fungi or free-living species feeding on bacteria—greater caution is warranted. Our data further indicate that the egg stage, lacking intestinal tissues, is least prone to contamination. Importantly, WGA HiFi data also showed that samples with higher bacterial contamination yielded more false-positive 6mA calls, which may explain earlier misinterpretations of 6mA in eukaryotes. Our analyses verified that the contamination ratio was as low as 0.01%, supporting the high reliability of our results.

Methylation is an epigenetic modification that influences gene expression and genome stability. The enrichment of 6mA sites in transposable element (TE) regions may affect their stability, expression, and chromatin structure, thereby shaping the three-dimensional organization of the genome and modulating chromatin accessibility. Because TE regions contain highly repetitive sequences prone to recombination and insertion, they often contribute to genomic instability and mutation(53). However, 6mA methylation may enhance TE stability and reduce such structural changes, thereby preserving essential genetic information and supporting species evolution and adaptability. Notably, while high-density TE methylation has been associated with reduced expression of neighboring TE-related genes (TEGs)(54), our results show that 6mA-marked TEGs exhibit significantly higher expression than unmarked ones.

Our results show that TE regions in polyploid nematodes harbor high-density 6mA methylation, whereas diploid nematodes lack such enrichment. Given that TEs are often associated with phenotypic diversity and stress responses (55), a comparative analysis of 6mA distribution between polyploid and diploid RKNs may help uncover the mechanistic driving these differences. Moreover, investigating the association between infection ability and 6mA across RKNs of different ploidy could provide further biological insights. While we observed distinct genomic and transcriptomic features between polyploid and diploid RKNs, these differences may not be attributable to polyploidy alone. Polyploid RKNs included here are also more closely related phylogenetically, raising the possibility that some features reflect shared ancestry rather than polyploidy per se. Disentangling polyploidy effects from lineage-specific adaptations will require broader taxon sampling, including both closely and distantly related diploid and polyploid species. Future studies incorporating independent polyploidization events and well-resolved phylogenies will be essential to clarify the respective roles of genome duplication and evolutionary history.

The AlkB family of dealkylating enzymes is conserved from bacteria to humans(56, 57). Humans encode nine AlkB proteins (AlkB1-8 and FTO), among which ALKBH1 and ALKBH4 are most strongly associated with demethylase activity. ALKBH1 has been confirmed as a demethylase in human(22), mice(58), and rice(19), while NMAD-1 in *C. elegans,* a homolog of ALKBH4, performs a similar role(13). In this study, we identified three proteins containing the 2OG-FeII_Oxy_2 domain from Mi. Our *in vivo* and *in vitro* assays demonstrated that two proteins, homologous to ALKBH6 and ALKBH7, act as putative 6mA demethylases in triploid Mi. Moreover, we found that genomic regions with different methylation levels corresponded to distinct sequence motifs, suggesting that these motifs may be recognized by different modifying enzyme systems.

Finally, our data suggest that 6mA can directly affect the expression of nematode virulence genes, but it is also likely to influence virulence indirectly through modulating of key transcription factors. Importantly, 6mA is a genome-wide epigenetic mark, and its role in nematode biology extends beyond virulence regulation to broader essential processes such as development, reproduction, and stress responses, consistent with its known effects on shaping chromatin accessibility and transcriptional dynamics. Moreover, *minmad-1*-HIGS in tomato and tobacco significantly enhanced the resistance to polyploid RKNs (Mi, Ma, and Mj), whereas rice expressing Mg-demethylase dsRNA showed no resistance to Mg, and demethylase knockout in *C. elegans* only modestly reduced egg-laying(13). These findings suggest functional divergence of demethylase between polyploid and diploid nematodes, potentially reflecting the greater importance of epigenetic regulation in parthenogenetic compared to sexually reproducing organisms. Further studies of 6mA analysis in additional parthenogenetic and sexual RKNs will be essential to further clarify its role across reproductive modes and ploidy levels. In summary, this study establishes a link between epigenetic regulation and pathogen virulence, and highlights 6mA as a promising target for nematode control, with broader implications for the management of other agricultural pathogens.

## Materials and Methods

### Nematode materials

Our previously work has constructed the chromosome-level genome assemblies for *M. incognita* WHF4-1, *M. arenria* SYMaF1-2, and *M. javanica* LongYF2-1., In addition, a telomere-to-telomere assembly of *M. graminicola* (Mg) and a newly assembled genome of *M. hapla* (Mh) were used in this study for subsequent 6mA DNA methylation analyses. Nematodes were propagated on Rutgers tomato under greenhouse conditions at 25°C, and eggs, J3J4 juveniles, and females were collected after six weeks for downstream experiments. DNA used for SMRT-seq, LC-MS/MS and Dot blot assays was extracted from surface-sterilized J2s inoculated into aseptically cultured tobacco grown in MS medium solidified with plant gel instead of agar. After 35 days, sterile eggs were harvested for subsequent analyses. Native DNA and WGA DNA used for HiFi sequencing and Mi-WGA 6mA-DIP-seq were obtained from eggs collected from nematodes infecting potted host plants. Prior to DNA extraction, eggs were disinfected with 10% sodium hypochlorite to minimize bacterial contamination. The protocol for stage-specific nematode collection is described in *SI Appendix*, Materials and Methods.

### Nematode gDNA Extraction

Genomic DNA was extracted from eggs of RKNs and cyst nematodes, as well as from mixed-stage individuals of *D. destructor*, *A. avenae*, *B. xylophilus*, *H. contortus*, *C. elegans*, and *C. nigoni* for subsequent experiments. Before DNA extraction, sodium hypochlorite was used to sterilize the nematodes and pass through a 3 µm filter to remove contamination such as bacteria. Genomic DNA isolation was performed as described in *SI Appendix*, Materials and Methods.

### Eliminate bacterial DNA contamination and RNA m6A contamination

To ensure the accuracy of 6mA detection in nematode DNA, multiple precautions were taken to eliminate potential contamination from bacterial DNA and RNA-derived m6A. For all initial tests, genomic DNA was extracted from nematode eggs. In root-knot nematodes, eggs were collected from parasitized roots maintained under sterile tobacco culture conditions. To further minimize bacterial contamination on the egg surface, eggs were disinfected with sodium hypochlorite, then ground and plated on LB and PDA media to assess the presence of cultivable endophytes. In addition, PCR amplification using universal 16S rRNA primers (27F and 1492R) was performed on the extracted DNA to screen for bacterial DNA contamination.

To eliminate RNA contamination in downstream 6mA detection assays such as dot blot and 6mA-DIP-seq, genomic DNA was treated with RNase A following extraction. RNA fragment sizes were monitored to confirm efficient digestion, with residual fragments reduced to less than 100 bp. The DNA was then sonicated and size-selected with magnetic beads to retain fragments larger than 200 bp. Finally, liquid chromatography–tandem mass spectrometry (LC-MS/MS) was employed to distinguish DNA-derived 6mA (m/z 266.1 → 150.0) from RNA-derived m6A (m/z 282.1 → 150.0), thereby confirming that the detected 6mA signals originated exclusively from DNA.

### 6mA-DIP-seq and analysis

The 6mA-DIP-seq was performed following the published protocols. Briefly, we fragmented 7 ug WHF4-1 RNA removed gDNA to 200-400 bp. After end repaired and adapter ligation by using Vazyme ND607 VAHTS Universal DNA Library Prep Kit for Illumina V3, DNA was heat-denatured and immunoprecipitated overnight at 4 °C with anti-6mA antibodies purchased from Abcam (ab151-230). Input and 6mA immunoprecipitated DNA was PCR amplified using Vazyme ND607 to construction the next generation sequencing library. The raw data were filtered by using Cutadapt (v3.3)(59) and aligned to WHF4-1 genome by using bowtie2 (v2.3.0)(60). The low quality and duplicate reads were removed by using SAMtools (v1.7)(61) and Picard (https://broadinstitute.github.io/picard/). Then, macs2 (v2.1.1)(62) was introduced to give the enriched regions to call 6mA peaks by comparing the reads between 6mA-IP and Input sample, with the following options: -B -SPMR -nomodel. The post file was further transformed into bigwig format for the profile visualization by using deeptools (v3.5.1)(63) and IGV (v2.5.2)(64). The MEME-ChIP(65) (https://meme-suite.org/meme/tools/meme-chip) was used to identify the conserved motifs enriched in 6mA peaks.

### DNA/RNA library construction and sequencing

Approximately 100 µl eggs were purified and collected for DNA extraction, and the genomic DNA was isolated using the CTAB method as described in previously. The DNA was sent to the ANNOROAD Gene Technology Co., Ltd. (Beijing, China) for PacBio library construction followed by sequel I platform sequencing. RNA extracting was using the TransZol Up Plus RNA Kit (TransGen Biotech, Beijinng, China) and RNA Nano 6000 Assay Kit of the Bioanalyzer 2100 system (Agilent Technologies, CA, USA) was using to assess the RNA integrity and concentration. The RNA library preparation was followed the protocol of VAHTS^®^ Universal V8 RNA-seq Library Prep Kit for Illumina (Vazyme, Nanjing, China). Finally, the library was pooled and sequenced on Illunima NovaSeq platform with 150 bp paired-end reads. Each treat of RNA sequencing is performed by using three biological repetitions.

### Hi-C library construction and sequencing

The eggs of Mh were used to construct the Hi-C library according to previously described method with appropriate modifications(44). Briefly, approximately 50 thousand newly collected eggs were pelleted and cross-linked in a 1.5% final concentration (v/v) formaldehyde (Sigma) for 30 min, followed de-crosslinking, eggs were homogenized and digested by using MboI (New England Biolabs). DNA was extracted after the addition of biotin, end-filling and ligation. Subsequently, the DNA was sheared and removed from the unligated fragment ends biotin mark by using T4 DNA polymerase. After biotinylated fragment pull-down, the target DNA was used to construct the Hi-C library by using End Prep Mix 4 from VAHTSTM Universal DNA Library Prep Kit for Illumina^®^ ND607 (Vazyme, Nanjing, China), and then sequenced on Illunima NovaSeq platform with 150 bp paired-end reads.

### Chromosome-level genome assembly

To obtain a high quality Mh genome, through the Pacbio data, we performed de novo genome assembly by using Canu (v1.9)(66) with default parameters. The initial assembled contigs were polished with Illumina short reads for 5 rounds by using BWA (v0.7.17)(67), SAMtools, sambamba (v0.7.1)(68), and pilon (v1.23)(69). Then, the Hi-C reads were aligned into the corrected contigs by using Juicer (v1.5.7) pipeline(70). The 3D-DNA (v180419) pipeline(71) was used to construct the chromosome-level scaffolds with parameter of -r 0, and we finally obtained the MhF1-2 genome with 99.18% sequences clustered into 16 chromosomes.

### Genome annotation

The egg, J2, J3J4, and female stages of Mh were collected to performed RNA-seq and subsequently use to genome annotation. RepeatModeler(72) and RepeatMasker(73) (http://www.repeatmasker.org) were used to soft-mask repetitive region of the assembled genome. The RNA-seq data was aligned to the masked genome by using HISAT2 (v2.2.1)(74), then, the BAM files were utilized to annotation based on ab initio prediction evidence by using AUGUSTUS (v3.3.3)(75), GeneMark-ES (v4.64)(76), and BRAKER2 (v2.1.6)(77). On the other hand, the RNA-seq data were de novo assembled using Trinity (v2.8.5)(78), and the assembled transcripts were used to the following integrated by PASA (v2.4.1)(79). The file augustus.hints.gff3 from Braker2 and the file pasa_assemblies.gff3 from PASA were used to integrate all evidence to obtain a final gene structure information by using EVidenceModeler (v1.1.1)(79). For the functional annotation, HMMER(80) was used to scan all the predicted proteins conserve domains based on Pfam database, and eggNOG-mapper (v2.0.1)(81) was used to perform the KEGG and GO annotation. For secreted protein prediction, we considered the proteins with signal peptide but without transmembrane domain were secreted proteins. The signal peptides were predicted by SignalP (v5.0)(82), and the transmembrane domains were identified by using TMHMM (2.0c)(83). The transposable elements (TEs) were predicted by using EDTA (v2.0.0) pipeline(84).

### 6mA identification and analysis by SMRT sequencing

We use SMRTLink-7.0.1 (http://www.pacb.com/support/software-downloads/) to identify the 6mA modification in WHF4-1, Mg, and Mh genome. The SMRT sequencing reads were aligned to the genome by pbmm2, and the 6mA modification sites were identified by ipdSummary with the following options: --methylFraction, --minCoverage 3, --identify m6A, --bigwig, --csv. The 6mA sites with less than 50 or 15 coverage were filtered according to the sequencing depth of each RKNs as described before(13, 17). For motifs identification, the 6mA sites were separated into three groups based on their methylation level (0-30%, 30%-70%, 70%-100%). We extracted 4 bp from the upstream and downstream sequences of each 6mA site and used BEDtools (v2.29.2)(85) to acquire the fasta file and followed by using MEME-ChIP to identify motifs for each group. 6mA peaks were loading to R package ChIPseeker (v1.24.0)(86) for associated genes annotation. The genome-wide distribution of 6mA sites with different methylation levels was visualized using Circos (v0.69)(87). To calculate the reads contamination rate of SMRT sequencing, we aligned the PacBio reads to bacteria, fungi, and plant NCBI nt library, the aligned length more than 90% of the query reads were considered as a valid alignment.

### 6mA distribution density analysis

The 6mA sites with different methylation levels were extracted into 6mAfilterHigh.bed, 6mAfilterMiddle.bed, and 6mAfilterLow.bed. We extended 4bp upstream and downstream of the 6mA site to obtain the *_8bp.bed files for subsequent analysis. BEDtools bedtobam was utilized to convert the *_8bp.bed files to *_8bp.bam files, and deeptools bamCoverage was used to generate *_8bp.bw files from *_8bp.bam files. Finally, the BED files containing exon and TE information, along with the *_8bp.bw files, were used to calculate the distribution density of 6mA in the gene or TE region using computeMatrix, and plotProfile was employed to visualize the results.

### DNA Dot blot

The Hybond+ membranes were fully soaked in double distilled water, then soaked in 20 SSC for 1h, sandwiched between two layers of filter paper and dried at 37 °C for 30 min. The purified genomic DNA was heated at 95 °C for 5 min to denature DNA and immediately placed on ice for 5 min and then 15 ng, 30 ng, and 50 ng DNA were loaded per dot on Hybond+ membranes. Membranes were air dry and placed them in the oven at 80 °C for 1 h, and transferred into boxes with damp paper towels. DNA was then auto-crosslinked twice for 6 min in an Ultraviolet crosslinker (CL-100). The membrane was blocked for 2 h in 5% skim milk TTBS. Membranes were incubated overnight at 4 °C with 6mA primary antibody (ab151-230, Abcam) in 5% skim milk TTBS. After three times wash for 5 min with TTBS, the blots were probed with secondary antibody in 5% skim milk TTBS for 1 h at room temperature. Blots were washed three times for 5 min with TTBS and then performed chemiluminescence detection. After the chemiluminescence is over, the membrane is directly used for methylene blue staining. (I) Immerse the membrane in 10mL of 1× methylene blue solution, stain for 3min on a shaker at room temperature, and wash the membrane twice with double distilled water for 5 minutes each time; stain again with methylene blue solution for 3min and wash the membrane twice with double distilled water for 5 minutes each time. Soak in TBST overnight at 4℃; (II) Stain with methylene blue solution for 5min, wash three times with double distilled water for 5 minutes each time. Take photos in the gel imaging instrument/camera.

### LC–MS/MS analysis of 6mA

The LC-MS/MS was performed as described before(13). Briefly, the nematodes genomic DNA was denatured at 100 °C for 3 min, and chilled in ice for 2 min immediately. Aspirate 26 µl of denatured DNA, add 1 µl of S1 nuclease, 3.2 µl of buffer, and treated at 37 °C overnight. One-tenth volume of 1M NH_4_HCO_3_ and 1 µl venom phosphodiesterase I (Sigma) was added to the solution and incubation at 37 °C for 2h. Finally, 1 U of alkaline phosphatase from *E. coli* (Sigma) was added to the solution at 37 °C for 2h. The digested genomic DNA was diluted two-fold with nuclease-free H_2_O and filtered through 0.22 µm filter. To determine the peak position of each nucleoside, the nucleoside standards including N6-methyldeoxyadenosine and deoxyadenosine nucleoside were used to perform the reverse-phase ultra-performance liquid chromatography on a C18 column (Agilent), with online mass spectrometry detection using triple-quadrupole tandem mass spectrometer (Shimadzu, LC-MS 8050) set to multiple reaction monitoring (MRM) in positive electrospray ionization mode. Simultaneous detection of standards and samples using established methods base ion mass transitions of 266.1-150.0 for 6mA and 252.1-136.0 for A, and calculate the ratio of 6mA/A.

### RNA interference *in vitro*

We select about 300 bp conserved regions as RNAi target fragments. We synthesized dsRNA using the T7 RNAi Transcription Kit-Box 1 (Vazyme, China, Cat. TR102-02). We followed the protocol for dsRNA synthesis and diluted the concentration of dsRNA to 2 µg/µl. The potential 6mA demethylase in vitro interference experiment was performed using WHF4-1 eggs. About 10,000 WHF4-1 eggs were sterilized and suspended in 100 µl of nuclease-free water, and an equal volume of dsRNA solution was added to make the final concentration of dsRNA to 1 µg/µl. After 60 hours of incubation at room temperature with 15 rpm rotation, the disturbed eggs were divided into two parts, one was used for qRT-PCR to detect the interference efficiency of the target gene, and the other was used to extract DNA to detect 6mA content by mass spectrometry. Three biological replicates were performed for each experiment, dsGFP was used as control.

### RNA-seq analysis

The RNA-seq data of *C. elegans* were downloaded from SRA database (accession number SRR6125151-SRR6125153). The raw reads of RNA-seq were cleaned by Cutadapt, the PCR duplications were removed by using Picard. We use STAR [84] to align the reads to RKNs genome and use RSEM(88) to calculate the TPM and FPKM value of each gene. For differentially expressed genes (DEGs) analysis, HISAT2 was used to align the RNA-seq data to the RKNs genome, then use SAMtools to convert the SAM files to BAM files and sort them, and finally use HTSeq-count (v0.13.5)(89) to calculate the expression count number of each gene. The count number was input into DESeq2 (v1.28.1)(90) to standardize and acquire the DEGs. Differentially expressed genes (DEGs) were identified using DESeq2, applying an adjusted p-value (FDR) < 0.05 and an absolute log2foldchange > 1 as thresholds. Comparisons of gene expression (TPM) between groups were performed using the Wilcoxon rank sum test, as TPM values showed non-normal distributions and unequal variances. Each treatment was analyzed with three biological replicates. The expression module analysis of secreted proteins was performed by using Mfuzz (v2.48.0)(91) and visualization by using R package pheatmap (v1.0.12). The GO and Pfam enrichment analysis were performed by python script and visualization by using R package ggplot2 (v3.3.3).

### Demethylase assays

*In vitro* demethylase activity experiments were performed using purified potential demethylase proteins of MiNMAD-1/2. The demethylase assays were performed as previously described(13). In brief, a triplicate of a 50 µl volumes containing 1 µg WHF4-1 gDNA and 1 mg recombinant protein in a reaction mixture (50 mM HEPES (pH 7.0), 50 mM KCl, 2 mM MgCl_2_, 2 mM ascorbic acid, 1 mM α-KG, and 200 mM (NH)_4_Fe(SO)_4_) were used to perform the reaction. After 1h hold at 37 °C, the reactions were stopped by adding 5mM EDTA and denatured at 100 °C for 3 min followed by chilling on ice for 2 min. The products of reactions were used to Dot blot and LC-MS/MS to determine the 6mA contents after putative demethylase treated.

### Verification of endogenous nematode 6mA demethylase

Verification of the nematode endogenous 6mA demethylase was carried out using protein extracts from WHF4-1 eggs, followed by incubation with biotin-labeled oligo DNA and pull-down with Dynabeads MyOne Streptavidin C1. The bound DNA was purified and analyzed to assess demethylase activity. Detailed experimental procedures are provided in *SI Appendix*, Materials and Methods.

### Polyclonal antibody preparation and western blot assays

All the work of polyclonal antibody preparation was performed by Friendbio Science & Technology (Wuhan, China). Briefly, the purified demathylase/Mi-CRT protein was used to immunize rabbits intradermally for antiserum production, and gel specific purification was used to purify the antibody. For western blot, the RKNs eggs, J2, and females were collected and homogenate. The protein concentration of the suspended was determined by using the BCA Kit (Beyotime, China). Proteins were separated on a 10% SDS-PAGE followed by blotting on PVDF membranes (Biosharp, China). The membranes were blocked 2h in 5% skim milk TTBS and probed with the demathylase/Mi-CRT antibody or β-actin (Proteintech, China, Cat No.: 66009-1-Ig) overnight. The secondary antibody and detection were performed as described in Dot blot.

### Demethylase prediction

We aligned the proteins sequence of WHF4-1 to the protein NMAD-1 of *C. elegans* by using a loose e-values by using blastp (v2.10.1)(92). The first three proteins on the alignment were selected as candidate demethylases. On the other hand, we performed Pfam domain annotation on all the proteins of WHF4-1 and found that 6 of them have the 2OG-FeII_Oxy_2 domain with potential 6mA demethylase function. Every three of these 9 candidate genes are alleles of each other, the coverage between alleles is 100%, and the minimum identity is 96%, therefore, we selected one of the three alleles for further validation studies.

### Transgenic plant construct

Nicotiana benthamiana was used to generate transgenic tobacco following the method of Agrobacterium-mediated genetic transformation of leaf discs(93). We selected the same segment as the above *in vitro* RNA interference as the segment for constructing transgenic plants expressing dsRNA. The Agrobacterium EHA105 was inoculated into YEB medium and cultured at 28 °C until OD600 up to 0.6-0.8. Put the tobacco leaf discs for about 6-7 weeks into the Agrobacterium suspension and soak for 10min and culture in the dark at 28 °C for 2-4 days. The explants were transferred to the Murashige and Skoog (MS) medium (with 1.0 mg/L hormones Benzylaminopurine, 0.1 mg/L Indole acetic acid, 100 mg/L antibiotics kanamycin, and 500 mg/L cefotaxime), and cultured under a light intensity of 25 μmol m⁻² s⁻¹ at 25 °C. After 4-5 weeks of selective culture, the explants will regenerate shoot buds and then transferred to fresh selective medium. When the shoot buds grow to 3-4 cm, transfer to the rooting medium (1/2 MS medium with the same concentration of antibiotics), and the roots will grow about 2-3 weeks. Using qRT-PCR to confirm putative T0 transgenic plants with specific primers described in *SI Appendix*, Materials and Methods. Further, PCR-positive plant was cultured in green house to obtain a homozygous T1 generation. Tomato transformation was performed using the ‘Moneymaker’ cultivar, following the same method as used for tobacco transformation. Cotyledons and hypocotyls from 5-day-old tomato seedlings were used for the transformation. The rice transgenic was completed by Wuhan Towin Biotechnology Co., Ltd.

## Author Contributions

M.S., J. Z., and D. D. conceived the project. M. S., J. Z., D. D., and D. P. administration the project. M. S. acquisition the funding. D.D. designed the experiments. D. D., S. Z. performed the experiments. S. Z. purified the protein and performed the in vitro protein activity experiments. S. Z., J. L., and J. S. constructed the transgenic plants. D. D. collected the data and performed the data analysis. Y.Z. and B.H. performed the Dot blot experiments. D. D., S. Z., X. W., S. C., Y. Z., M. Z., B. H., G. W., W. W., and F. W. prepared the materials. D. D. wrote the manuscript, S. S., and V.M.W. revised the manuscript.

## Competing Interest Statement

The authors declare that they have no competing interests.

## Acknowledgements

We thank Huazhong Agricultural University for technical support; Weiguo Lu of Henan Academy of Agricultural Sciences for providing cyst nematode X12 material; Min Hu of Huazhong Agricultural University for providing the *Haemonchus contortus* species; Xiaoli Guo of Huazhong Agricultural University for providing the cyst nematodes Hg3 and Hg5 species; Fengjuan Zhang of Hubei University for providing the *Aphelenchus avenae* and *Bursaphelenchus xylophilus* species.

## Notes

### Competing Interest Statement

The authors have declared no competing interest.

### Summary of Updates

We revised and adjusted the entire article, refined the language, deleted some content that was not highly relevant to the main conclusions of this study, updated the Discussion and Introduction, and moved part of the Methods to the Supplementary File.

