## Supplemental Files for "N6-Methyladenine DNA modification modulates pathogen virulence in nematodes"

**This PDF file includes:**

Supporting Materials and Methods
Figures S1 to S35
Tables S1 to S15

### Materials and Methods

#### Nematode collection

Three stages of RKNs were used in this study. For egg collection, cut the nematode-infected tomato or tobacco roots into 0.5 cm size, put it into a 500 ml beaker, followed by adding 225 ml ddH2O and 25 ml sodium hypochlorite, and place it on a magnetic stirrer for 8 minutes at 1000 rpm. The suspension was successively passed through 18, 60, 100, 200 and 500 mesh sieves, rinsed five times with water to remove sodium hypochlorite, and the eggs were retained on the 500-mesh sieve. After sucrose gradient centrifugation with a final concentration of 35%, the sucrose was passed through a 3um filter membrane using a negative pressure suction filter, and the eggs retained on the 3 μm filter membrane were collected for the next experiment. For first-stage juveniles (J1) collection, the purified eggs were incubated for 2-3 days at 20 °C, J1 larvae were manually picked out under a microscope until most of the eggs became intertwined worms. For J2 collection, the purified eggs were incubated on a 500-mesh sieve with water for 3-5 days at 20 °C, and the J2 larvae were collected from the water under the mesh. For J3J4 and female collection, the roots after treated with sodium hypochlorite to collect the egg were put into a mortar, then use a pestle to smash the roots lightly, suspend the roots with 100 ml of water, and pass them through 18, 60, 100, and 200 mesh sieves in sequence. The J3J4 were on the 200 mesh sieves, and the female were on the 60 mesh sieves. After most plant debris was removed by sucrose centrifugation, J3J4 were manually picked under a microscope, and the females were picked visually. For early female (eFemale) collection, the tobacco roots 21 days after nematode infestation were collected, and early females before oviposition were manually picked out with forceps.

#### Nematode gDNA Extraction protocol

Genomic DNA was isolated using the CTAB method as follow: (1) eggs were suspended in 200 μl SDS-EB lysis buffer (50 mM Tris-HCl pH 8.0, 200 mM NaCl, 20 mM EDTA, 2% SDS, 1mg/ml proteinase K) and transfer into liquid nitrogen; (2) after homogenized, the powder was collected into 1.5 ml tube and add SDS-EB lysis buffer to 500 μl; (3) add 500 μl 65 °C preheat CTAB buffer (100 mM Tris-HCl pH 8.0, 20 mM EDTA, 1.4 M NaCl, 2% CTAB, 1% PVP 40000) and incubation at 65°C for 30 min; (4) equal volume phenol-chloroform-isoamyl alcohol mix (25:24:1) were added to separate the DNA and protein, mixture were spun at 13,000 rpm at 4 °C for 15 min, collect the supernatant; (5) 2 μl 10 mg/ml RNase A was added into the supernatant and incubated at 37 °C for 15 min; (6) 0.75 M final concentration ammonium acetate was added and DNA was sedimented in pre-cold isopropanol (0.9:1, v/v) and finally dissolution in 30 to 50 μl DNase-free water.

#### RNA interference primer

The PCR primers used to amplify the RNAi target fragment template are as follows: *minmad-1* F: 5'-GGTGGAATAATAAGTGGCTT-3', *minmad-1* R: 5'-CTGATTGTGGAATTATTTTC-3'; *minmad-2* F: 5'-CCTCAATTTATTTCGGAAGC-3', *minmad-2* R: 5'-CCTGTCCAGGCAAATATTCA-3'; *Mi_23849.1* F: 5'-GGATAGAAAAATGTCTGTTG-3', *Mi_23849.1* R: 5'-GCAGCATCAGGAAGGAAATC-3'; *gfp* F: 5'-GAGTGCCATGCCCGAAGGTTA-3', *gfp* R: 5'-GGTCTGCTAGTTGAACGCTTCC-3'. *Mg_06820* F: 5'-GAAACAGAAGAAGCAAAT-3', *Mg_06820* R: 5'-AGTAGAAATCATTGGAAAG-3'; *Mg_00565* F: 5'-GTATTGGATTAAATAATTTCTTC-3', *Mg_00565* R: 5'-TTTGTATAGATTCTTCACTAA-3'.

#### Protocol for verification of endogenous nematode 6mA demethylase

About 50 μl WHF4-1 eggs were collected and suspended in 200 μl nematode lysis buffer (50 mM HEPES, 150 mM NaCl, 10% glycerol, 1% Triton X-100, 1.5 mM $MgCl_2$, and 1 mM EGTA) with proteinase inhibitor. After adding 100 mg of 1 mm diameter zirconia microspheres to the suspension, the samples were crushed by a tissue grinder (JXFSTPRP-24L, Shanghaijingxin Experimental technology, China), and the oscillation frequency was 70 Hz for 30 seconds, and the repetition was repeated 5 times. Suck up 20 μl suspension into a new 1.5 ml centrifuge tube, and add 64 μl $ddH_2O$, 10 μl reaction mixture, 1 μl protease inhibitor (Roche), 5 μl 20 μM oligo DNA. The suspension was incubated with rotation at 12 rpm for 3-6 hours at 28 °C. To separate the oligo DNA from Mi's own DNA, the biotin-labeled oligo DNA was pulled down using Dynabeads MyOne Streptavidin C1 beads (Thermo Fisher Scientific). 100 μl 2× BB (10 mM Tris-HCl (pH 7.5); 1 mM EDTA; 2 M NaCl) suspended Dynabeads MyOne Streptavidin C1 beads were mixed with the suspension and incubated at room temperature for 30 min with rotation (15 rpm). The beads with biotin marked DNA were washed three times with 300 μl TWB (5 mM Tris-HCl (pH 7.5); 0.5 mM EDTA; 1 M NaCl; 0.05% Tween 20) and incubated at 55 °C for 2 min. Added 27 μl dd$H_2O$ to resuspend the beads, treat them in a dry bath at 98 °C for 10 minutes, centrifuge and collect the supernatant after magnetic stand separation. The experiment of treating oligo DNA with purified potential demethylase is the same as the above method, and the eggs suspension should be replaced with purified protein.

We noticed that the 6mA content of the control group was as high as about 40%, which was quite different from the 6mA content of the designed oligo DNA. We suspect that the increased ratio of 6mA is due to the degradation of part oligo DNA by nucleases present in the environment. Nevertheless, the experimental group and the control group still showed a significant difference in 6mA content. The method of 6mA content detection is consistent with that described above, and the oligo DNA sequence modified with 6mA, and biotin are shown as below:

| Oligo name | Oligo sequence |
|---|---|
| 6mA_motif | GGGAATTTCCCGGCGATTTGA(N6-Me-dA)GGTAATCGCCGGGAAATT(biotin)CCC |

**Transgenic plant qRT-PCR primer**

| Primer name | Primer sequence |
|---|---|
| MiNMAD-1 F | TACCTCAATAATGCGGCTTGTT |
| MiNMAD-1 R | ACTCTCTTCATTAGCCAGGATTTC |
| MiNMAD-2 F | CCTCAATTTATTTCGGAAGCAGAA |
| MiNMAD-2 R | TCTAACCACATCGGCAATTCTT |
| Mi_23849.1 F | TCCGAGACAATCTTTACCTGATGA |
| Mi_23849.1 R | AAATCGCCCAAGCCTAGACTA |

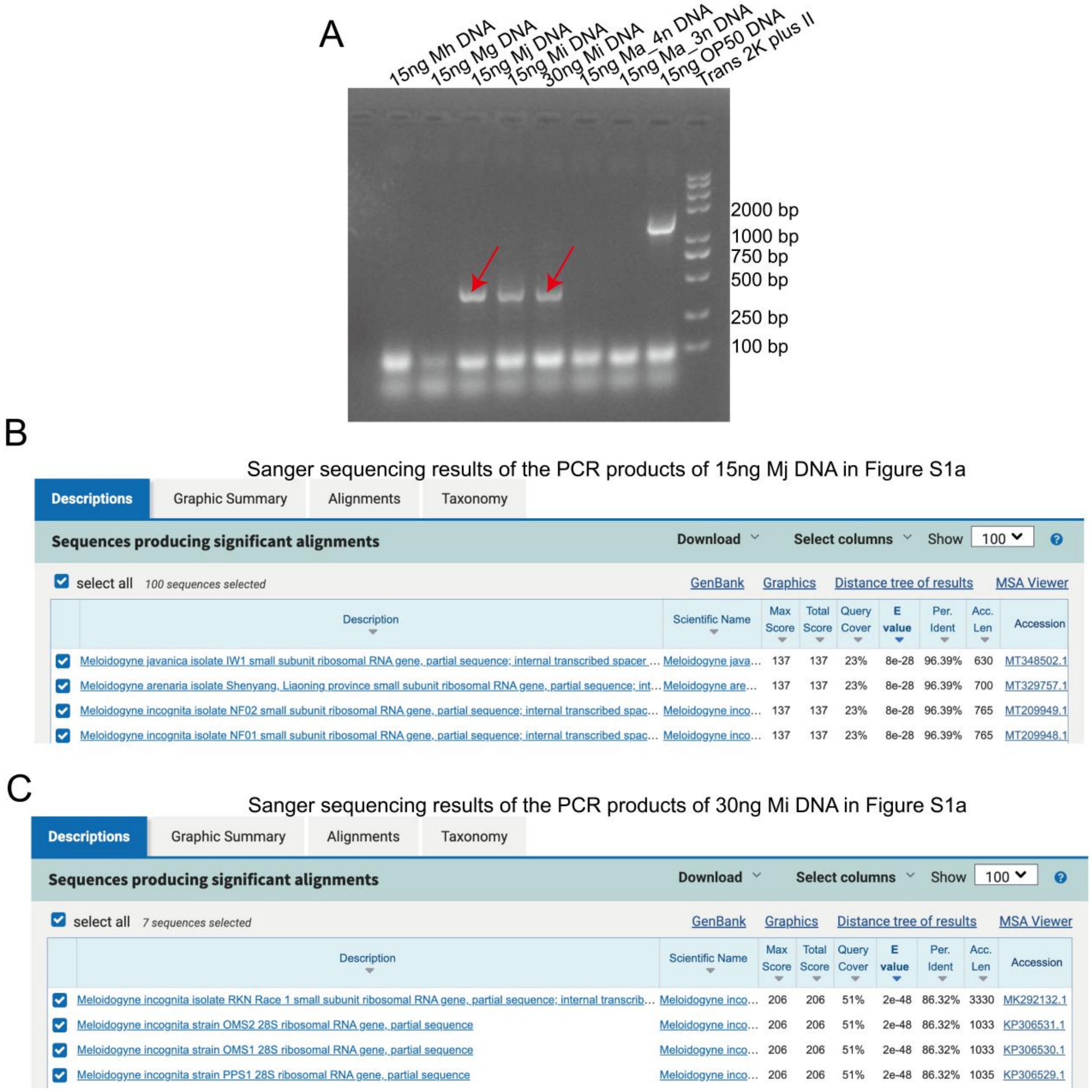


| Description | Scientific Name | Max Score | Total Score | Query Cover | E value | Per. Ident | Acc. Len | Accession |
|---|---|---|---|---|---|---|---|---|
| Meloidogyne javanica isolate IW1 small subunit ribosomal RNA gene, partial sequence; internal transcribed spacer … | Meloidogyne java… | 137 | 137 | 23% | 8e-28 | 96.39% | 630 | MT348502.1 |
| Meloidogyne arenaria isolate Shenyang, Liaoning province small subunit ribosomal RNA gene, partial sequence; int… | Meloidogyne are… | 137 | 137 | 23% | 8e-28 | 96.39% | 700 | MT329757.1 |
| Meloidogyne incognita isolate NF02 small subunit ribosomal RNA gene, partial sequence; internal transcribed spac… | Meloidogyne inco… | 137 | 137 | 23% | 8e-28 | 96.39% | 765 | MT209949.1 |
| Meloidogyne incognita isolate NF01 small subunit ribosomal RNA gene, partial sequence; internal transcribed spac… | Meloidogyne inco… | 137 | 137 | 23% | 8e-28 | 96.39% | 765 | MT209948.1 |



| Description | Scientific Name | Max Score | Total Score | Query Cover | E value | Per. Ident | Acc. Len | Accession |
|---|---|---|---|---|---|---|---|---|
| Meloidogyne incognita isolate RKN Race 1 small subunit ribosomal RNA gene, partial sequence; internal transcrib… | Meloidogyne inco… | 206 | 206 | 51% | 2e-48 | 86.32% | 3330 | MK292132.1 |
| Meloidogyne incognita strain OMS2 28S ribosomal RNA gene, partial sequence | Meloidogyne inco… | 206 | 206 | 51% | 2e-48 | 86.32% | 1033 | KP306531.1 |
| Meloidogyne incognita strain OMS1 28S ribosomal RNA gene, partial sequence | Meloidogyne inco… | 206 | 206 | 51% | 2e-48 | 86.32% | 1033 | KP306530.1 |
| Meloidogyne incognita strain PPS1 28S ribosomal RNA gene, partial sequence | Meloidogyne inco… | 206 | 206 | 51% | 2e-48 | 86.32% | 1035 | KP306529.1 |

**Fig. S1.**

**The 16S amplification indicates that no detectable bacterial DNA is present in the nematode DNA.** (*A*) The 16S primers (27F and 1492R, the target sequence was about 1500 bp) were used to amplify the DNA extracted from the nematode eggs used in the this study, and OP50 DNA was used as a positive control. The DNA template amplification of the experimental group Mj and Mi produced a non-specific band of about 350 bp. The sequence indicated by the red arrow was subjected to Sanger sequencing to determine whether it was contaminated by bacterial DNA. (*B*) The sequence blast results of the PCR product of the Mj template showed that it was aligned to the *M. javanica* small subunit ribosomal RNA gene. (*C*) The sequence blast results of the PCR product of the Mi template showed that it was aligned to the *M. incognita* small subunit ribosomal RNA gene.

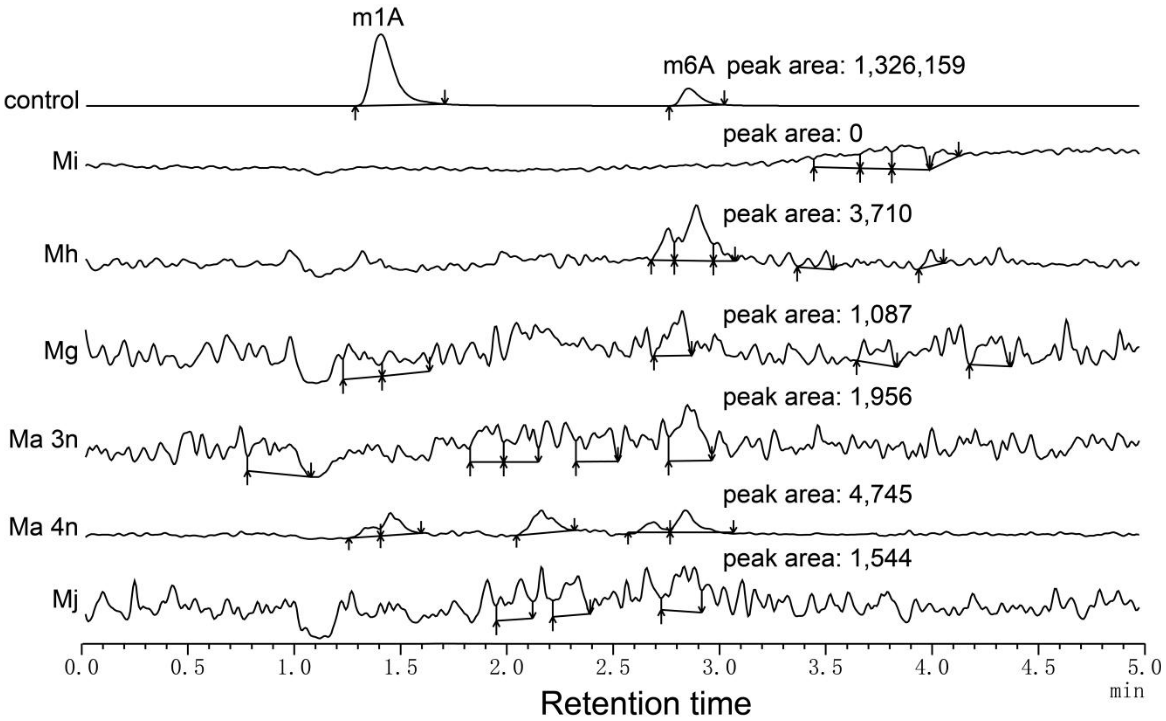


**Fig. S2.**

**Mass spectrometry is used to identify whether RNA m6A contamination has been eliminated in DNA samples, thereby ensuring that Dot blot and 6mA-DIP-seq samples reflect the true 6mA information on DNA.** Due to the different molecular weights, the molecular weight detected by mass spectrometry for RNA m6A is 282.1, and the daughter ion is 150.0; the molecular weight detected by mass spectrometry for DNA 6mA is 266.1, and the daughter ion is 150.0. Therefore, mass spectrometry can be used to determine whether the DNA sample contains RNA m6A contamination. The mass spectrometer used in this study was Shimadzu's triple quadrupole LC-MS 8050, and the detection mode was MRM. The control group consisted of DNA samples without RNA removal treatment, in which strong m1A and m6A signals were clearly detected.

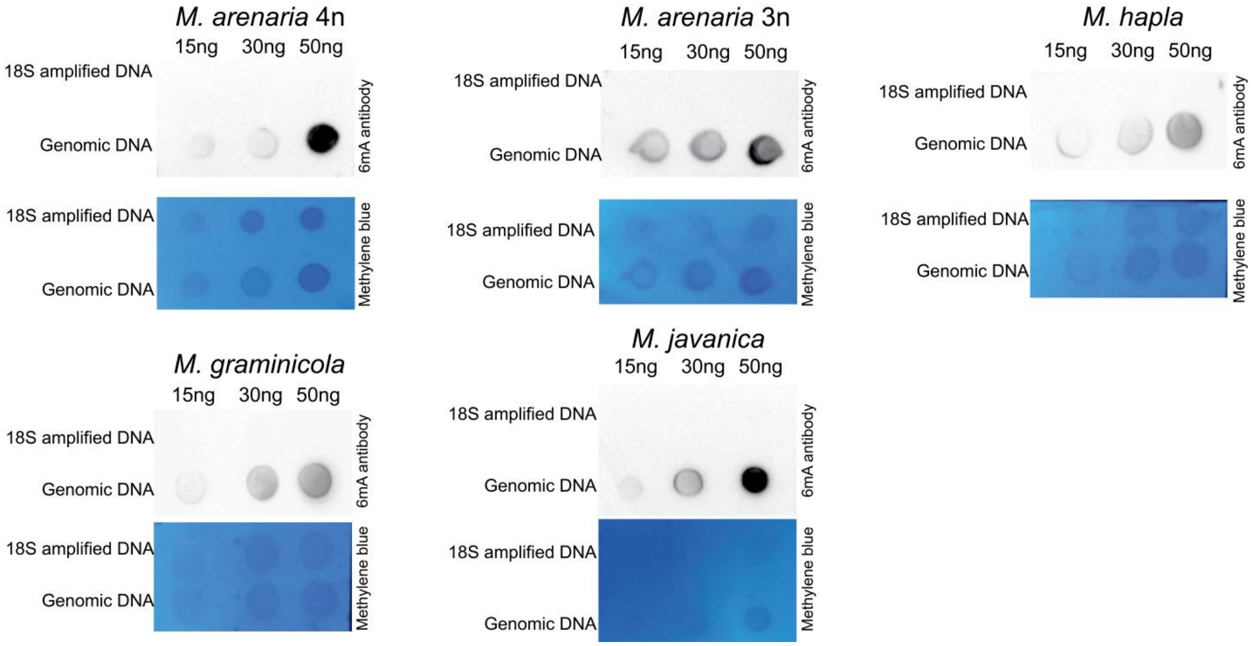


**Fig. S3.**

**The DNA 6mA Dot blot of *M. arenaria* 4n, *M. arenaria* 3n, *M. hapla*, *M. graminicola*, and *M. javanica*.** The amount of DNA used for each Dot is marked below the corresponding Dot. The 6mA antibody come from Abcam (ab151-230). RNA contamination in DNA was eliminated by RNase A treatment and then screening for fragments larger than 200 bp using magnetic beads, and LC-MS/MS confirmed that it did not contain RNA m6A.

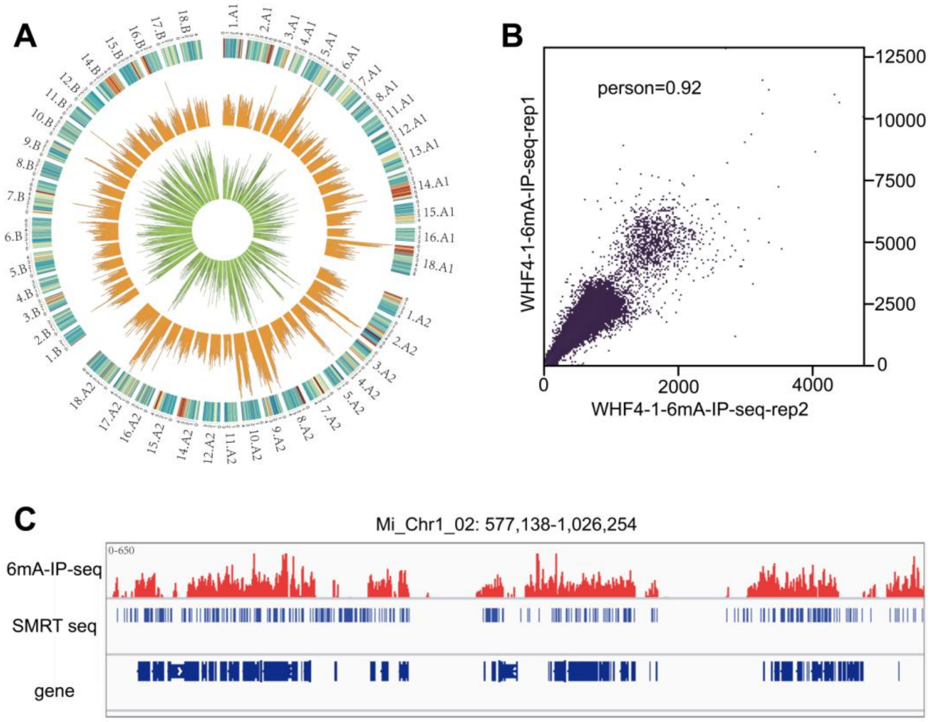


D

| | number of peak | number of peak overlap with SMRT | the proportion of overlapped peaks |
|---|---|---|---|
| 6mAIP-seq rep1 | 17,804 | 9,578 | 53.8% |
| 6mAIP-seq rep2 | 16,924 | 8,947 | 52.9% |

**Fig. S4.**

**Concordance of 6mA-IP-seq with SMRT sequencing in *M. incognita* WHF4-1.** (*A*) Circos plot of 6mA sites from 6mA-IP-seq (green) and SMRT sequencing (orange), outer ring are the expression level and the genome size of *M. incognita*. (*B*) Pearson correlation coefficient between two biological replicates for 6mA-IP-seq. (*C*) Local genome distribution of 6mA peaks identified by 6mA-IP-seq and SMRT-seq. (*D*) Comparison of the number of 6mA peak overlap identified by 6mAIP-seq and SMRT-seq. The possible reason for the low overlap of the data is that the DNA used in the two experiments came from two independent DNA extractions taken several months apart.

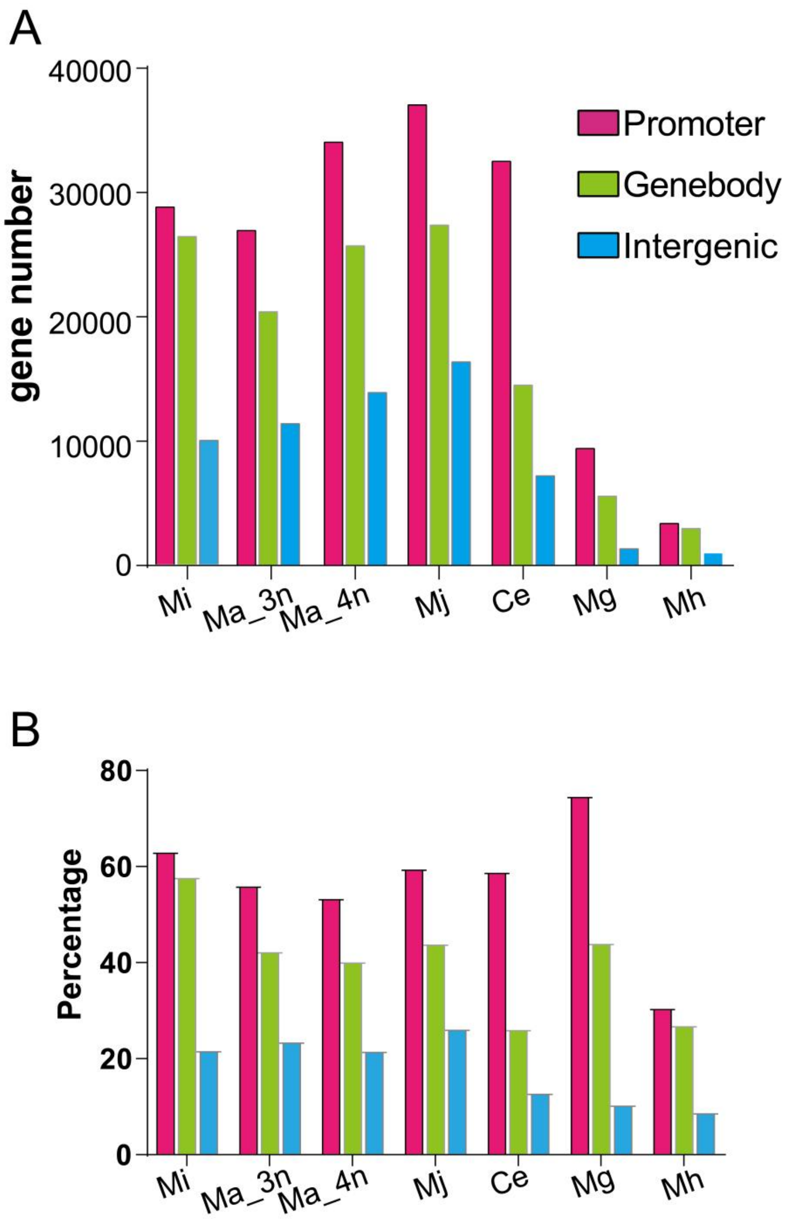


**Fig. S5.**

**Statistics of the distribution characteristics of 6mA methylation in the genome of each species.** (*A*) The gene number of 6mA distribution on different positions of *M. incognita* (Mi), *M. hapla* (Mh), *M. graminicola* (Mg), *M. arenaria* 3n (Ma_3n), *M. arenaria* 4n (Ma_4n), and *M. javanica* (Mj). (*B*) The percentages represent the proportion of genes with 6mA methylation in each genomic region (promoter, gene body, or intergenic) relative to the total number of genes in the genome. Because some genes carry methylation in more than one region, the summed percentages across categories exceed 100%.

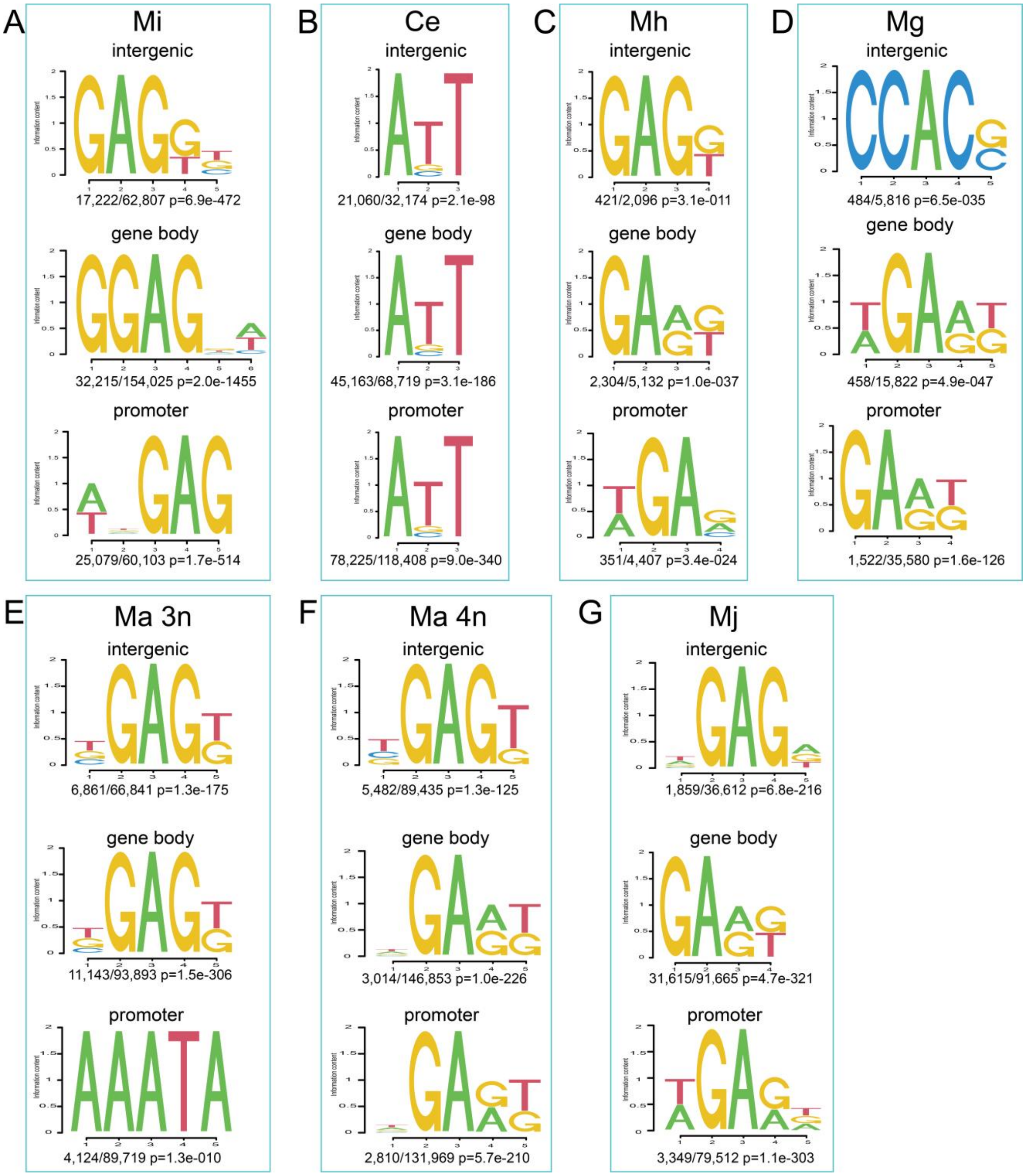


**Fig. S6.**

**Motif analysis of the 6mA site modified on Promoter, Gene body, and Intergenic.** *M. incognita* (Mi), *C. elegans* (Ce), *M. hapla* (Mh), *M. graminicola* (Mg), *M. arenaria* 3n (Ma_3n), *M. arenaria* 4n (Ma_4n), and *M. javanica* (Mj). According to the position of the 6mA site on the genome, the 6mA sites located at promoter, gene body, and intergenic were extracted respectively, and 4 bp sequences upstream and downstream of the site were taken for motif analysis. The motifs of Ce and RKNs are completely different. Most of the motifs of RKNs are relatively similar, the motif of promoter in Ma 3n and the motif of intergenic in Mg are divergence.

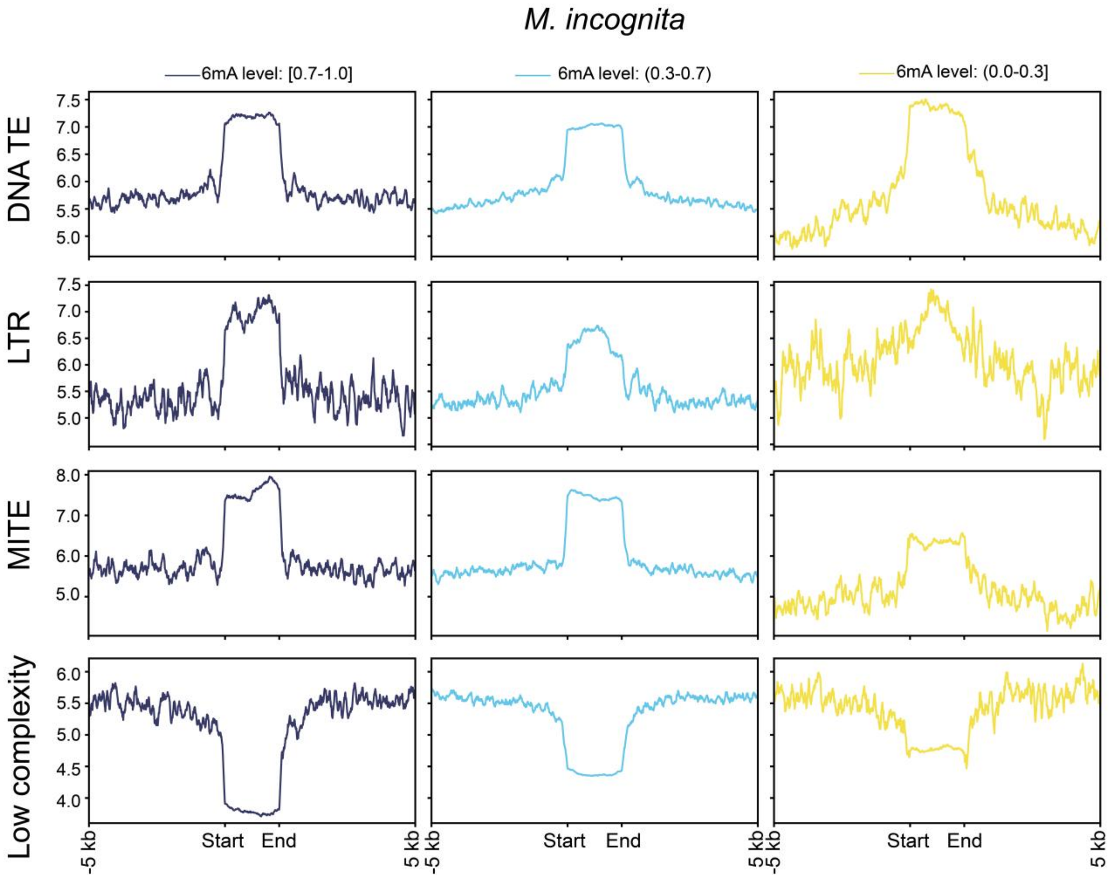


**Fig. S7.**

**The 6mA distribution feature in different types of TE of *M. incognita*.** The DNA TE include DNA/DTA, DNA/DTC, DNA/DTH, DNA/DTM, DNA/DTT, and DNA/Helitron. The DNA TE number count amount to 209,728 in Mi. The LTR include LTR/Copia, LTR/Gypsy, and LTR/unknown. The LTR number count amount to 37,746 in Mi. The MITE include MITE/DTA, MITE/DTC, MITE/DTH, MITE/DTM, and MITE/DTT. The MITE number count amount to 59,704 in Mi. The low complexity includes simple_repeat, and the number count amount to 125,990 in Mi.

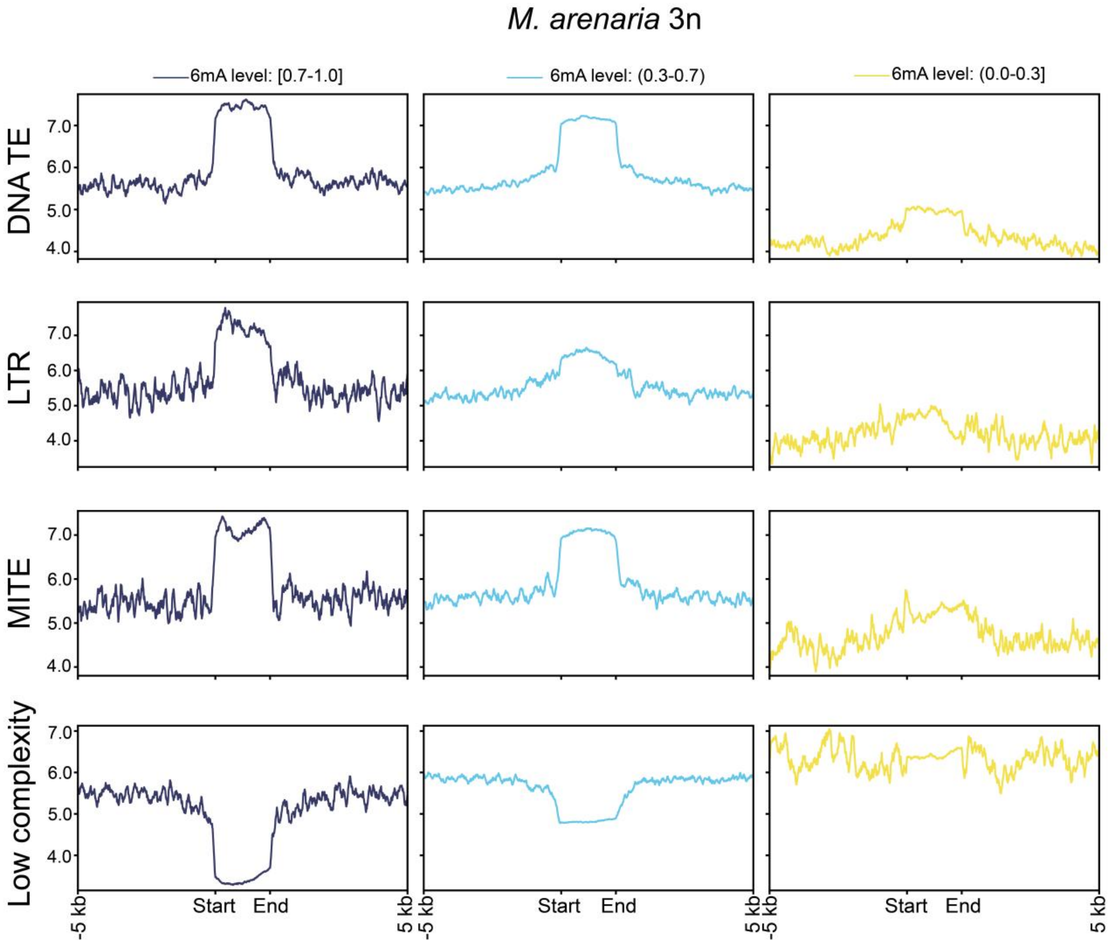


**Fig. S8.**

**The 6mA distribution feature in different types of TE of *M. arenaria* 3n.** The DNA TE include DNA/DTA, DNA/DTC, DNA/DTH, DNA/DTM, DNA/DTT, and DNA/Helitron. The DNA TE number count amount to 215,296 in Ma_3n. The LTR include LTR/Copia, LTR/Gypsy, and LTR/unknown. The LTR number count amount to 45,072 in Ma_3n. The MITE include MITE/DTA, MITE/DTC, MITE/DTH, MITE/DTM, and MITE/DTT. The MITE number count amount to 67,514 in Ma_3n. The low complexity includes simple_repeat, and the number count amount to 130,601 in Ma_3n.

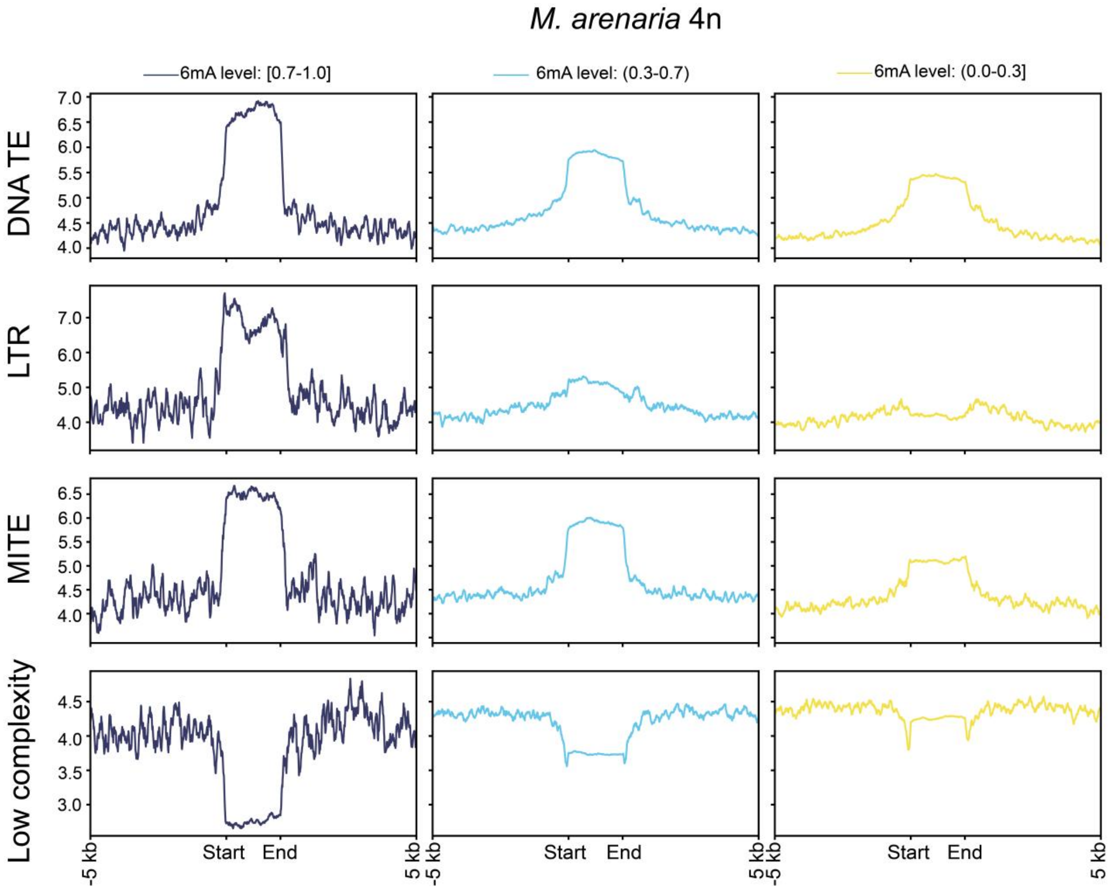


**Fig. S9.**

**The 6mA distribution feature in different types of TE of *M. arenaria* 4n.** The DNA TE include DNA/DTA, DNA/DTC, DNA/DTH, DNA/DTM, DNA/DTT, and DNA/Helitron. The DNA TE number count amount to 671,106 in Ma_4n. The LTR include LTR/Copia, LTR/Gypsy, and LTR/unknown. The LTR number count amount to 135,608 in Ma_4n. The MITE include MITE/DTA, MITE/DTC, MITE/DTH, MITE/DTM, and MITE/DTT. The MITE number count amount to 193,610 in Ma_4n. The low complexity includes simple_repeat, and the number count amount to 171,034 in Ma_4n.

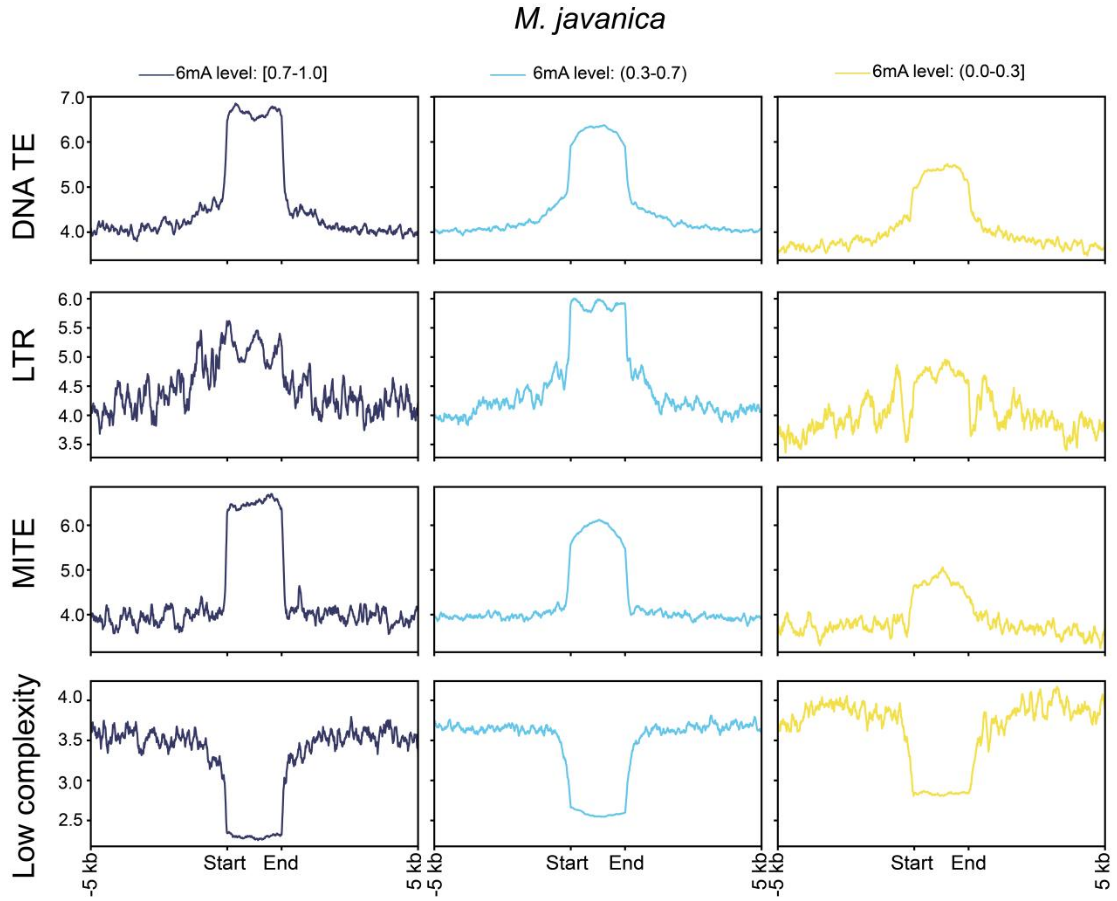


**Fig. S10.**

**The 6mA distribution feature in different types of TE of *M. javanica*.** The DNA TE include DNA/DTA, DNA/DTC, DNA/DTH, DNA/DTM, DNA/DTT, and DNA/Helitron. The DNA TE number count amount to 629,168 in Mj. The LTR include LTR/Copia, LTR/Gypsy, and LTR/unknown. The LTR number count amount to 118,400 in Mj. The MITE include MITE/DTA, MITE/DTC, MITE/DTH, MITE/DTM, and MITE/DTT. The MITE number count amount to 193,180 in Mj. The low complexity includes simple_repeat, and the number count amount to 169,725 in Mj.

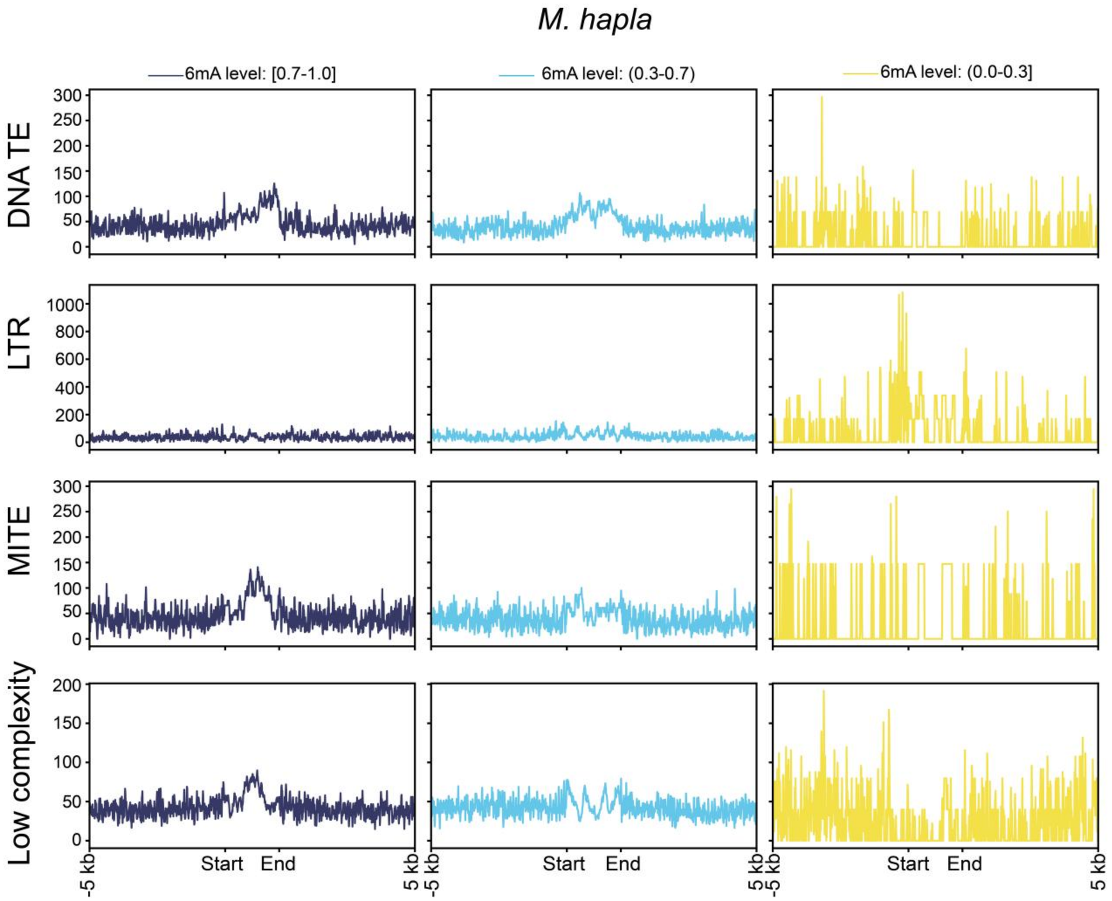


**Fig. S11.**

**The 6mA distribution feature in different types of TE of *M. hapla*.** The DNA TE include DNA/DTA, DNA/DTC, DNA/DTH, DNA/DTM, DNA/DTT, and DNA/Helitron. The DNA TE number count amount to 4,805 in Mh. The LTR include LTR/Copia, LTR/Gypsy, and LTR/unknown. The LTR number count amount to 2,165 in Mh. The MITE include MITE/DTA, MITE/DTC, MITE/DTH, MITE/DTM, and MITE/DTT. The MITE number count amount to 2,187 in Mh. The low complexity includes simple_repeat, and the number count amount to 8,053 in Mh.

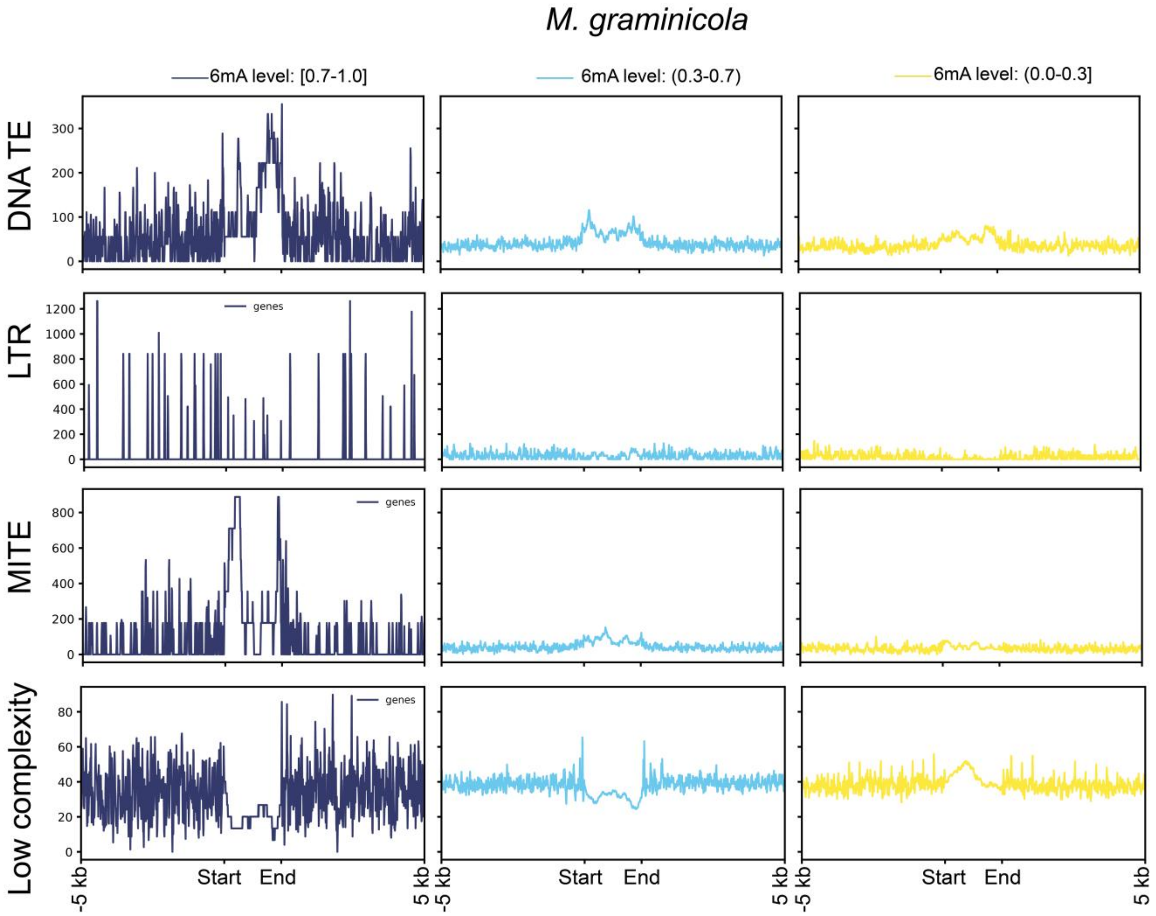


**Fig. S12.**

**The 6mA distribution feature in different types of TE of *M. graminicola*.** The DNA TE include DNA/DTA, DNA/DTC, DNA/DTH, DNA/DTM, DNA/DTT, and DNA/Helitron. The DNA TE number count amount to 8,448 in Mg. The LTR include LTR/Copia, LTR/Gypsy, and LTR/unknown. The LTR number count amount to 556 in Mg. The MITE include MITE/DTA, MITE/DTC, MITE/DTH, MITE/DTM, and MITE/DTT. The MITE number count amount to 2,187 in Mg. The low complexity includes simple_repeat, and the number count amount to 2,640 in Mg.

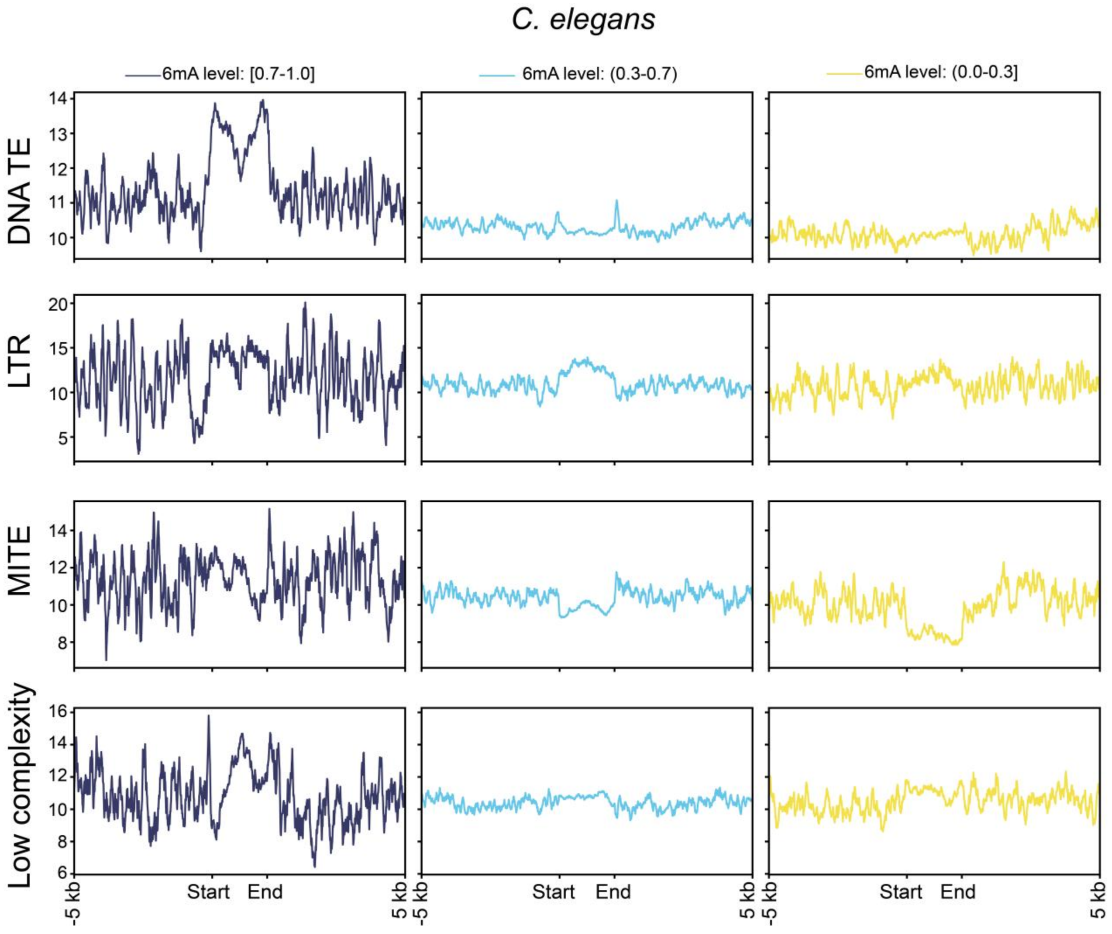


**Fig. S13.**
**The 6mA distribution feature in different types of TE of *C. elegans*.** The DNA TE include DNA/DTA, DNA/DTC, DNA/DTH, DNA/DTM, DNA/DTT, and DNA/Helitron. The DNA TE number count amount to 50,658 in Ce. The LTR include LTR/Copia, LTR/Gypsy, and LTR/unknown. The LTR number count amount to 1,773 in Ce. The MITE include MITE/DTA, MITE/DTC, MITE/DTH, MITE/DTM, and MITE/DTT. The MITE number count amount to 6,435 in Ce. The low complexity includes simple_repeat, and the number count amount to 6,068 in Ce.

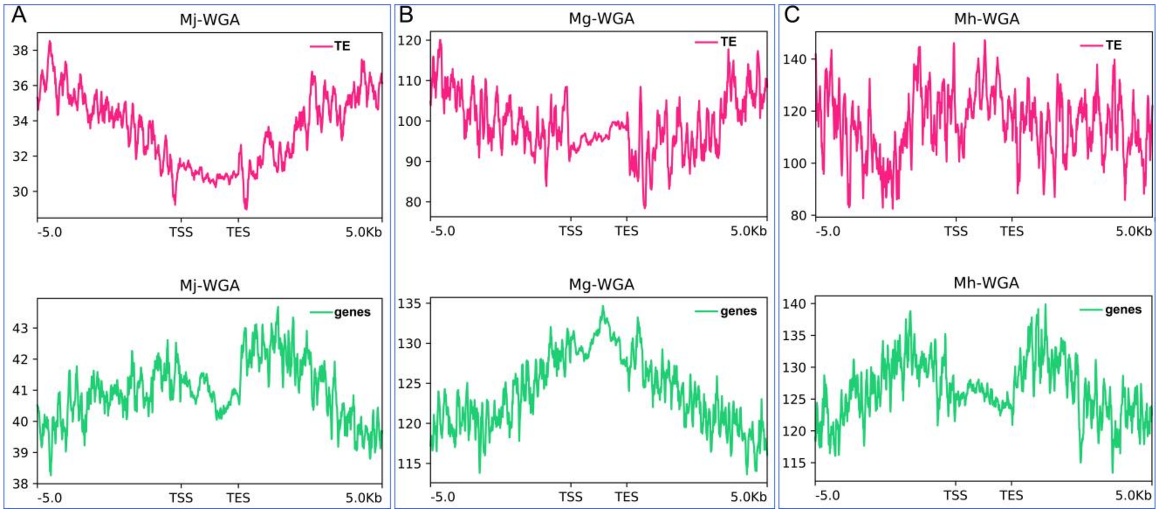


**Fig. S14.**

**The distribution density of false positive 6mA sites in WGA samples in TE and gene regions did not change significantly.** (**a**) The distribution density of 6mA false positive sites in the TE and Gene regions of the *M. javanica* WGA sample did not show the phenomenon of increased density in the TE region and decreased density in the gene region as in the native DNA samples. (**b**) The distribution density of 6mA false positive sites in the TE and gene regions of the *M. graminicola* WGA sample. (**c**) The distribution density of 6mA false positive sites in the TE and gene regions of the *M. hapla* WGA sample.

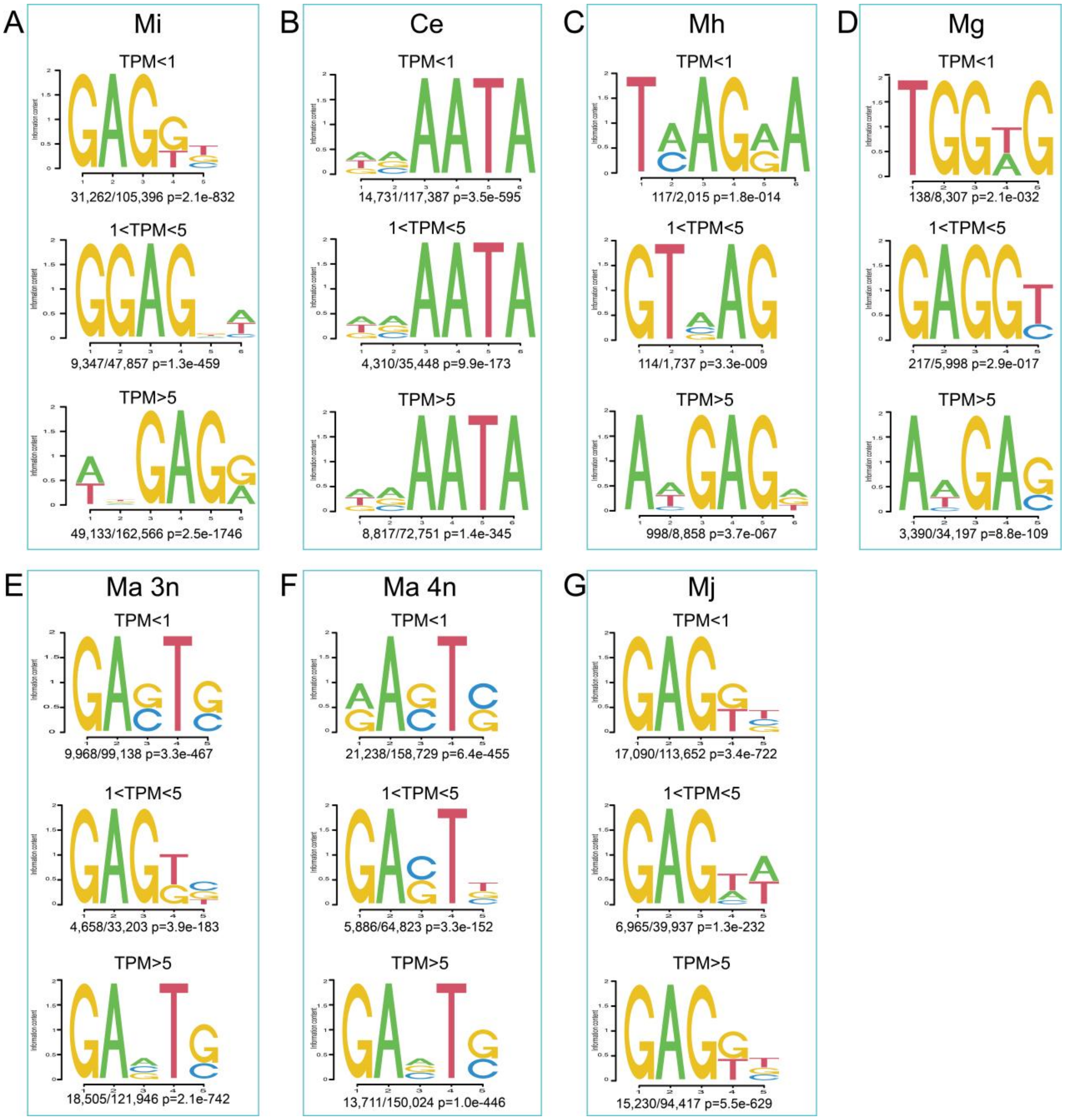


**Fig. S15.**

**Motif analysis for the 4bp up and downstream of the 6mA site modified on gene expression level at TPM <1, 1<TPM<5, and TPM>5.** *M. incognita* (Mi), *C. elegans* (Ce), *M. hapla* (Mh), *M. graminicola* (Mg), *M. arenaria* 3n (Ma_3n), *M. arenaria* 4n (Ma_4n), and *M. javanica* (Mj). The motifs of Ce and RKNs are completely different. The motifs of Ma 3n and Ma 4n are relatively similar. The motifs of Mi, Mj and Mg are relatively similar. The motifs of Mh and other RKNs are divergence.

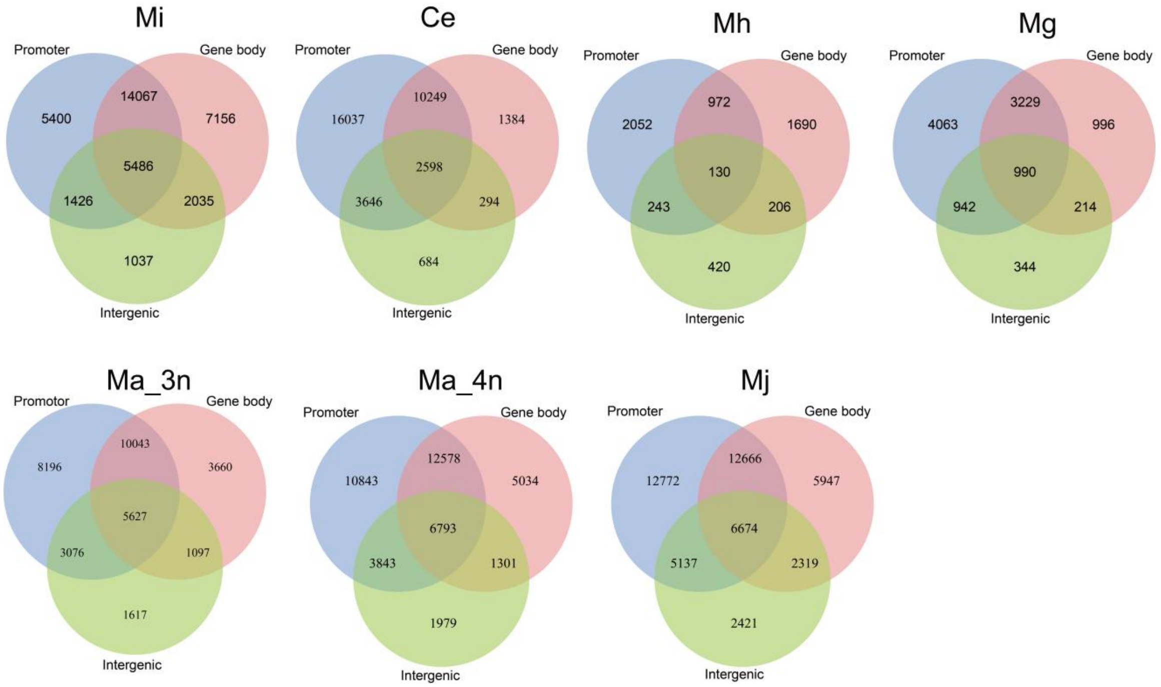


**Fig. S16.**

**Venn diagram showing the overlap between the genes with 6mA modified on promoter, gene body, and intergenic of these nematodes.** *M. incognita* (Mi), *C. elegans* (Ce), *M. hapla* (Mh), *M. graminicola* (Mg), *M. arenaria* 3n (Ma_3n), *M. arenaria* 4n (Ma_4n), and *M. javanica* (Mj).

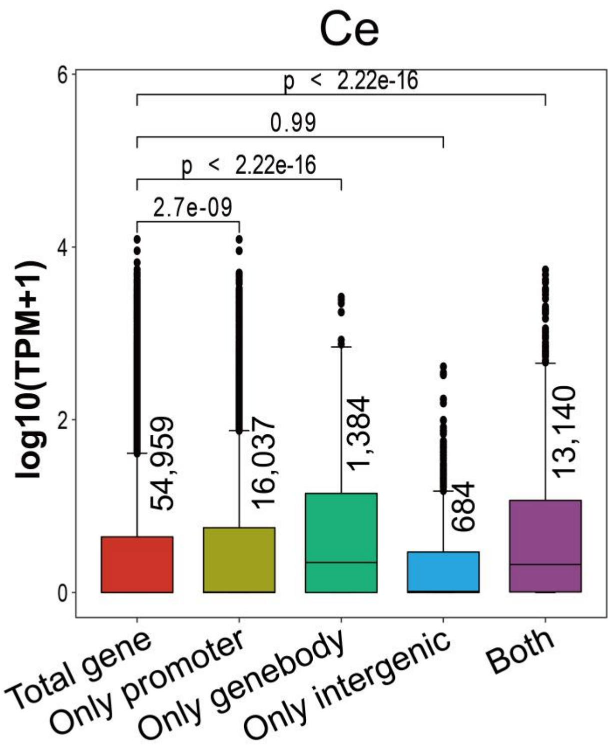


**Fig. S17.**

**The comparing of expression levels between the genes modified by 6mA at different positions and total gene of *C. elegans* (Ce).** N number were showed upon the box. In box plots, the central line represents the median, the box represents the 25% and 75% percentiles, and the whiskers represent 1.5 times the interquartile range beyond the box. *p* values were calculated by two-sided Wilcoxon rank sum tests. In some plots, the median line is present but overlaps with the lower quartile or whisker and may therefore not be visible.

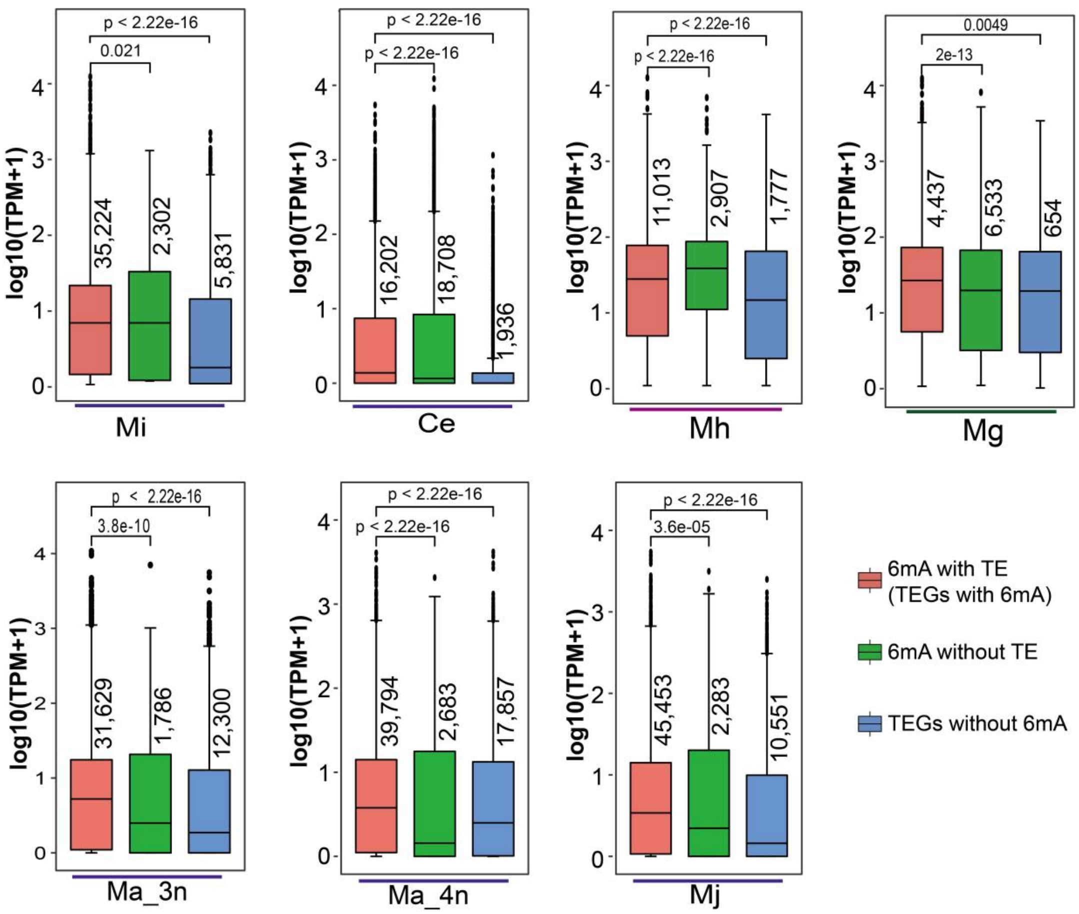


**Fig. S18.**

**The gene expression level comparing between 6mA with TE and 6mA without TE, and between TE with 6mA and TE without 6mA in these nematodes.** The expression levels of genes with both TE and 6mA nearby, and those with only 6mA modification showed different characteristics in different nematodes, while TEGs without 6mA had lower expression levels than TEGs with 6mA in all nematodes. *M. incognita* (Mi), *C. elegans* (Ce), *M. hapla* (Mh), *M. graminicola* (Mg), *M. arenaria* 3n (Ma_3n), *M. arenaria* 4n (Ma_4n), and *M. javanica* (Mj). N number for each box was showed upon the box. In box plots, the central line represents the median, the box represents the 25% and 75% percentiles, and the whiskers represent 1.5 times the interquartile range beyond the box. *p* values were calculated by two-sided Wilcoxon rank sum tests.

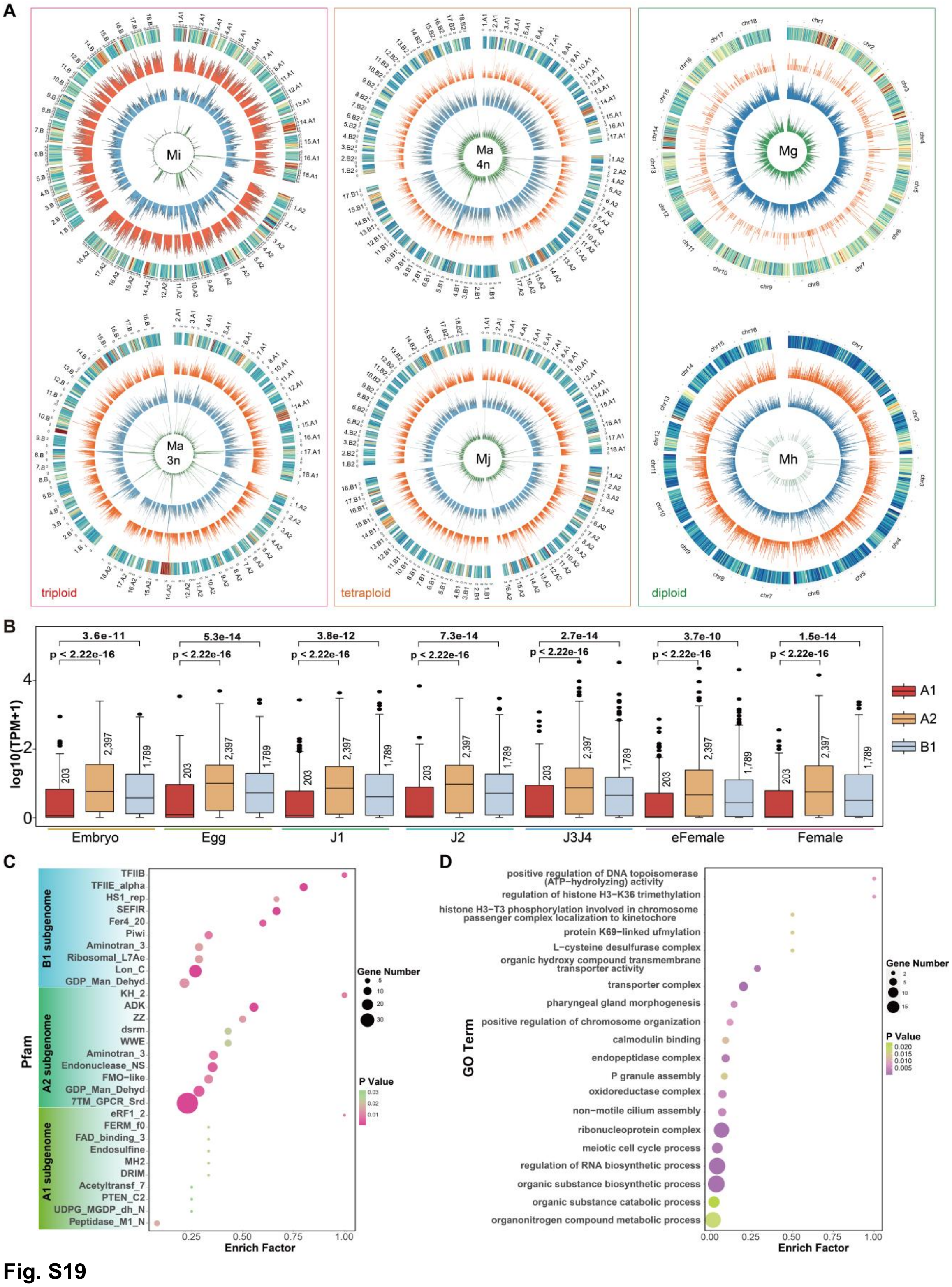


**Fig. S19**

**The lowly methylation 6mA sites play an important role in *M. incognita*.** (*A*) Circos plots of the density distribution of 6mA (in 50kb genome window) across all chromosomes of *M. incognita* (Mi), *M. arenaria* 3n (Ma 3n), *M. arenaria* 4n (Ma 4n), *M. javanica* (Mj), *M. graminicola* (Mg), and *M. hapla* (Mh) in the different 6mA modification level categories. Green, blue, and orange circles represent lowly methylated (0%-30%], moderately methylated (30%–70%), and highly methylated

[70%–100%] 6mA sites, respectively. The circle of heatmap represents the z-score of gene expression levels at the corresponding genomic locations during the egg stage. The redder the color, the higher the expression level; the bluer the color, the lower the expression level. The 6mA distribution feature of *C. elegans* (Ce) is not displayed here because it has been reported. (*B*) Genes associated with high-density-region (6mA sites counting beyond 60 in a 50 kb genome window) of low-methylation-level 6mA sites in the $A_2$ subgenome had higher expression levels at each growth stage than genes at the same location in the $A_1$ and $B_1$ subgenomes. (*C*) The Pfam enrichment analysis of the largest number of low methylation level 6mA sites associated gene from $A_2$ subgenome and the same position of $A_1$ and B subgenome. (*D*) The GO enrichment analysis of the largest number of low methylation level 6mA sites associated gene from $A_2$ subgenome. *p* values were calculated by two-sided Wilcoxon rank sum tests.

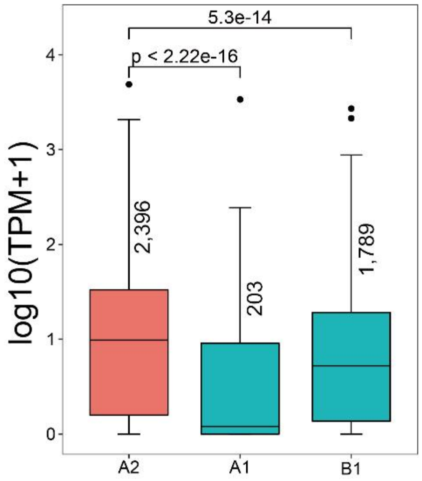


**Fig. S20**
**The gene located in high-density-region (HDR) of low-methylation-level 6mA sites (6mA counts beyond 60 in a 50 kb genome window) in A2 and A1, B1 subgenome were extracted to compared the expression level.** N number were showed upon the box. In box plots, the central line represents the median, the box represents the 25% and 75% percentiles, and the whiskers represent 1.5 times the interquartile range beyond the box. *p* values were calculated by two-sided Wilcoxon rank sum tests.

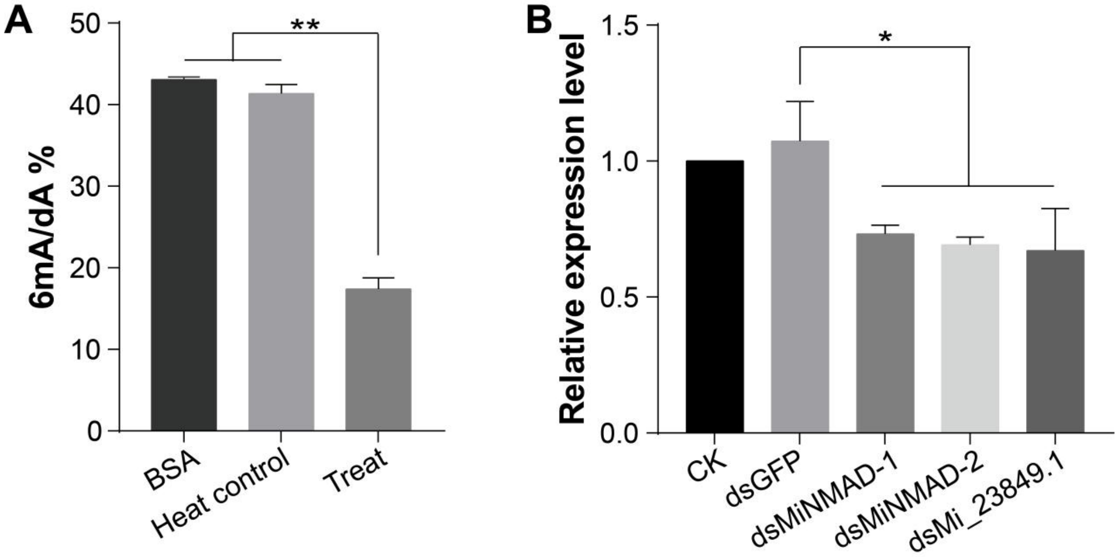


**Fig. S21.**

***M. incognita* (Mi) endogenous demethylase verification.** (*A*) Demethylase activity was measured with the suspension of broken Mi eggs, "BSA" was the negative control, "Treat" was the experimental group, and "Heat control" was the enzyme inactivation control after the suspension of broken eggs was bathed in boiling water for 10 minutes. (*B*) The qRT-PCR results of RNAi efficiency of candidate genes.

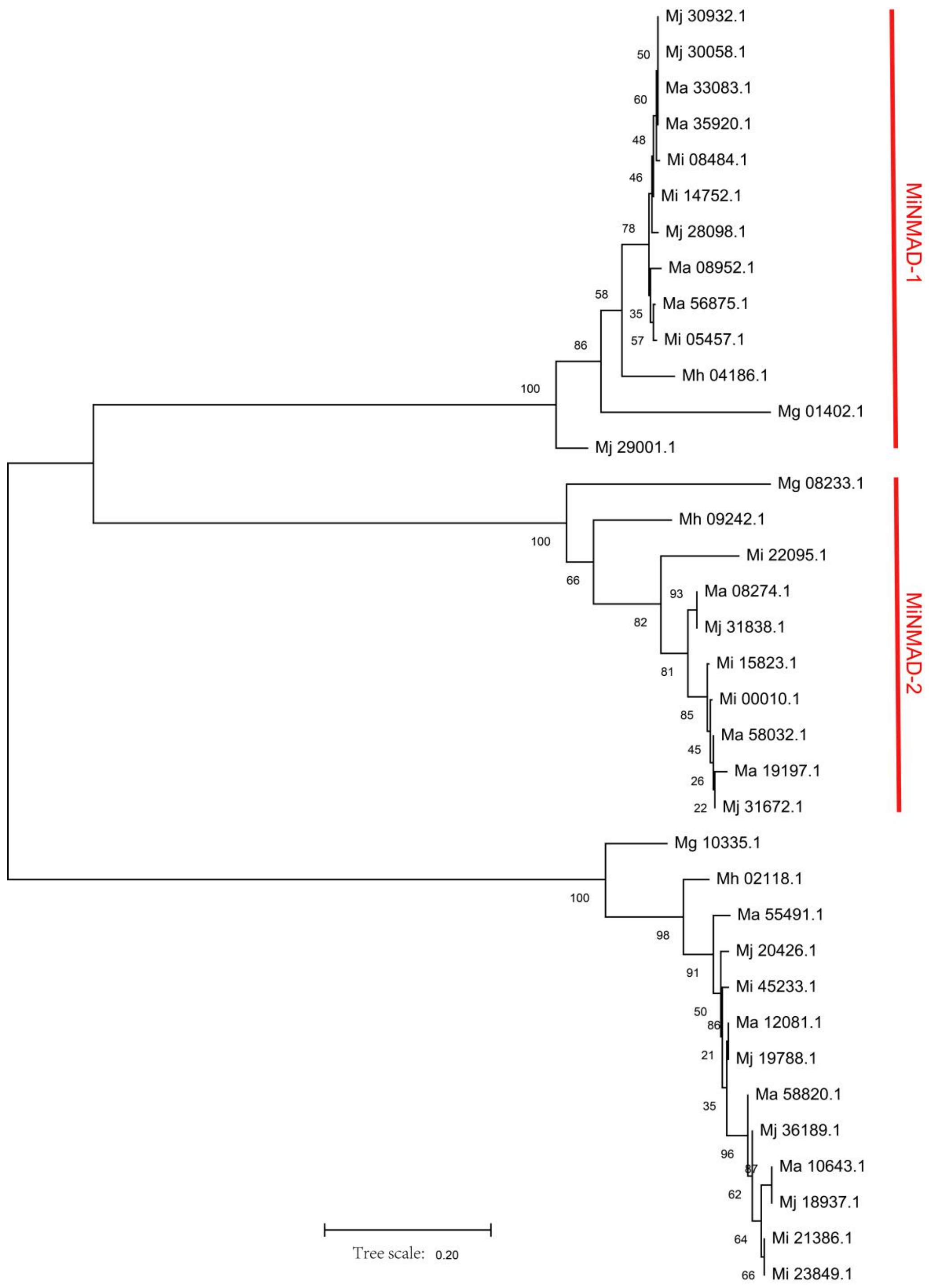


**Fig. S22.**

**The maximum-likelihood tree is construct from the protein sequence of MiNMAD-1/2 and Mi_23849.1 of *M. incognita*, and the relative protein sequence of Ma 4n, Mj, Mg, and Mh.**

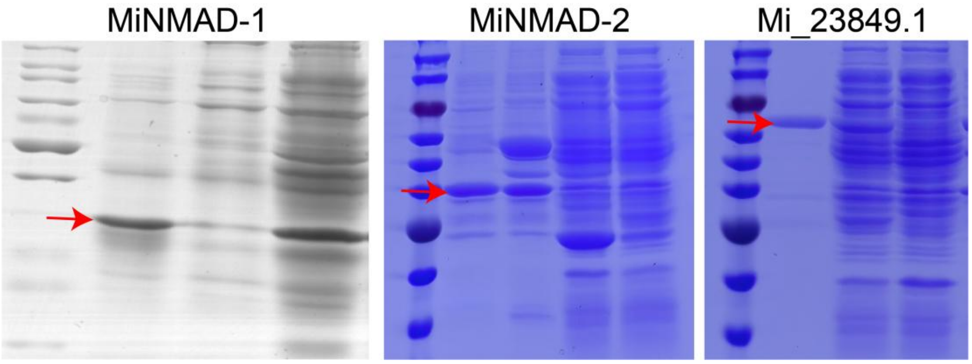


**Fig. S23.**
**The nematode target proteins were expressed in *E. coli* Rosetta and purified, with MiNMAD-1 fused to a SUMO tag, and MiNMAD-2 and Mi_23849 fused to GST tags.**

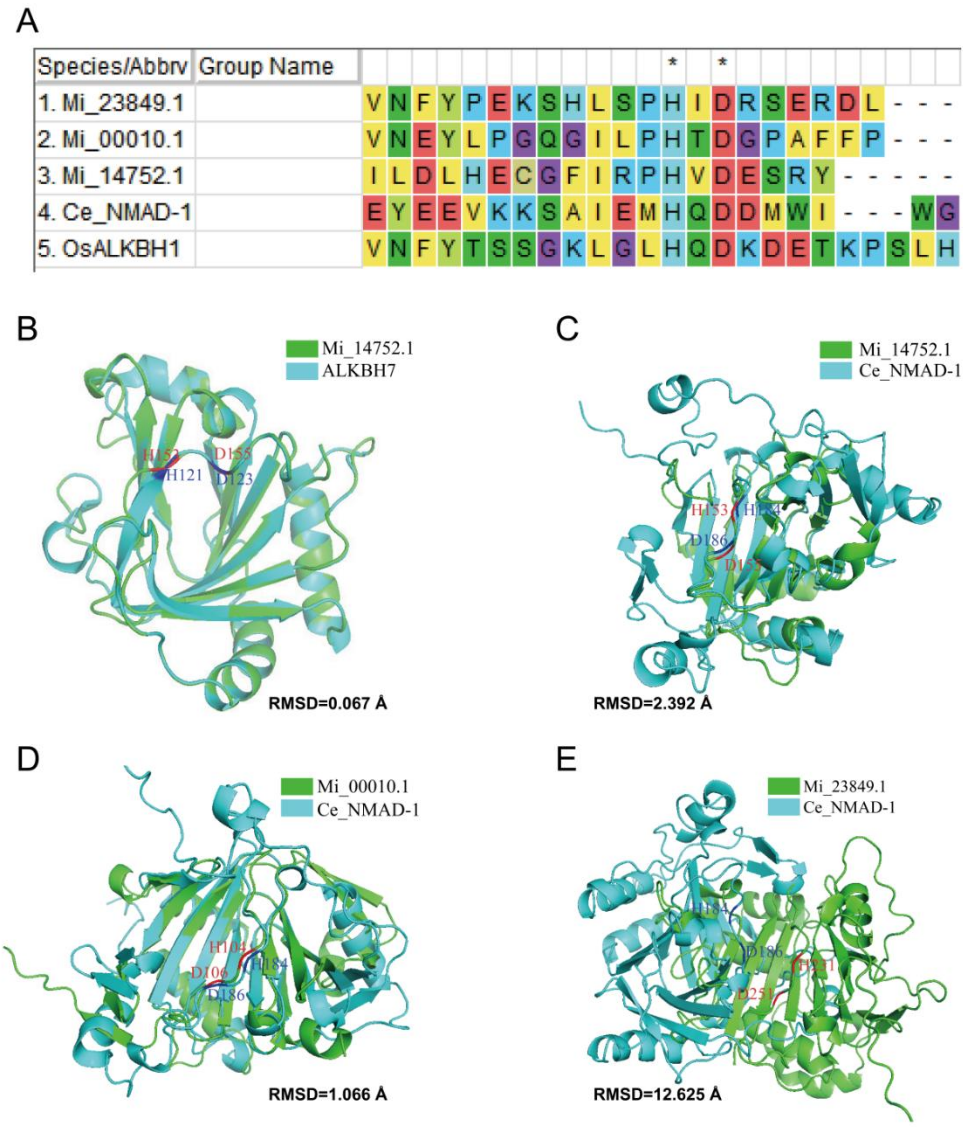


**Fig. S24.**

**Identification of *M. incognita* demethylase catalytic active sites.** (*A*) The candidate demethylase in Mi was compared with the known catalytic domain *C. elegans* (Ce) and *Oryza sativa* L. (Os) 6mA demethylase sequences, and the results showed that these proteins are conserved in the catalytic site. The star * means the reported 6mA catalytic site in Ce and Os. (*B*) The structure of Mi_14752.1 predicted by Alphafold2 was compared with the structure of human ALKBH7 (4QKD) in the Protein Data Bank. (*C*) The comparing of Alphafold2 predicted structure of Mi_14752.1 and Ce_NMAD-1. (*D*) The comparing of Alphafold2 predicted structure of Mi_00010.1 and Ce_NMAD-1. (*E*) The comparing of Alphafold2 predicted structure of Mi_23849.1 and Ce_NMAD-1. The blue text indicates the catalytic site of ALKBH7 and Ce_NMAD-1, and the red text indicates the corresponding amino acid in Mi_14752.1, Mi_00010.1, and Mi_23849.1. The lower the RMSD value, the more similar the structures of the two proteins being compared are.

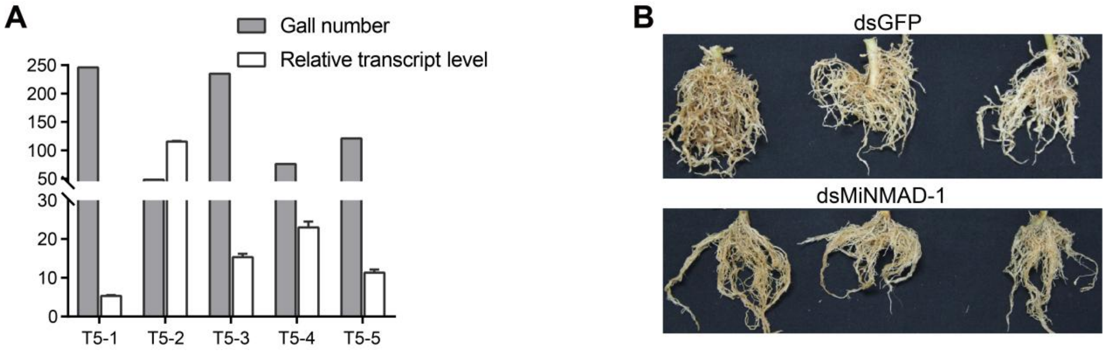


**Fig. S25.**
**The transgenic tobacco with host-induced gene silencing (HIGS) of *minmad-1* exhibits significant resistance to *M. incognita*.** (*A*) The galls number was inversely proportional to *minmad-1* dsRNA expression level, T5-1 to T5-5 represent different transgenic lines expressing *minmad-1* dsRNA. (*B*) Pot experiment show that *minmad-1* dsRNA high expression lines had only a few small galls after 8 weeks of Mi infection, while all roots of the control group were covered with large galls.

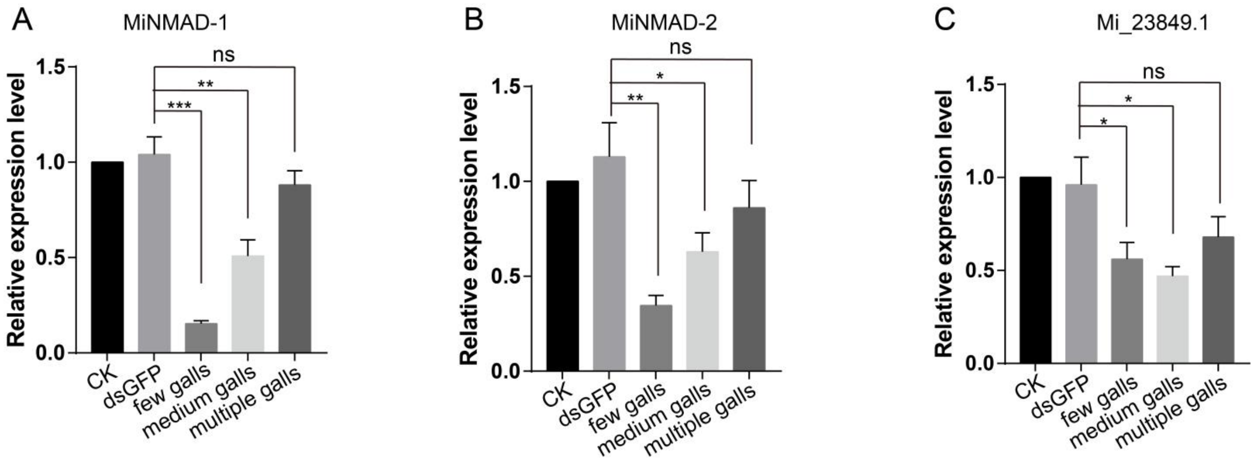


**Fig. S26.**

**Relative expression levels of corresponding genes in *M. incognita* egg after infecting transgenic tobacco.** (*A*) The relative expression level of Mi *minmad-1* gene after infecting tobacco expressing *minmad-1* dsRNA. (*B*) The relative expression level of Mi *minmad-2* gene after infecting tobacco expressing *minmad-2* dsRNA. (*C*) The relative expression level of Mi *Mi_23849.1* gene after infecting tobacco expressing *Mi_23849.1* dsRNA. All results were determined by qRT-PCR. The galls number in the medium and few galls groups of the *minmad-1/2*-HIGS group was significantly less than that of the control group. The galls number in the *Mi_23849*-HIGS group was equivalent to that of the control group. The grouping of medium and few galls relies on experience, and there is no significant analysis between the two group. *p* values were calculated by two-tailed unpaired Student's *t*-test. One star * means $p < 0.05$, two means $p < 0.01$, three means $p < 0.001$.

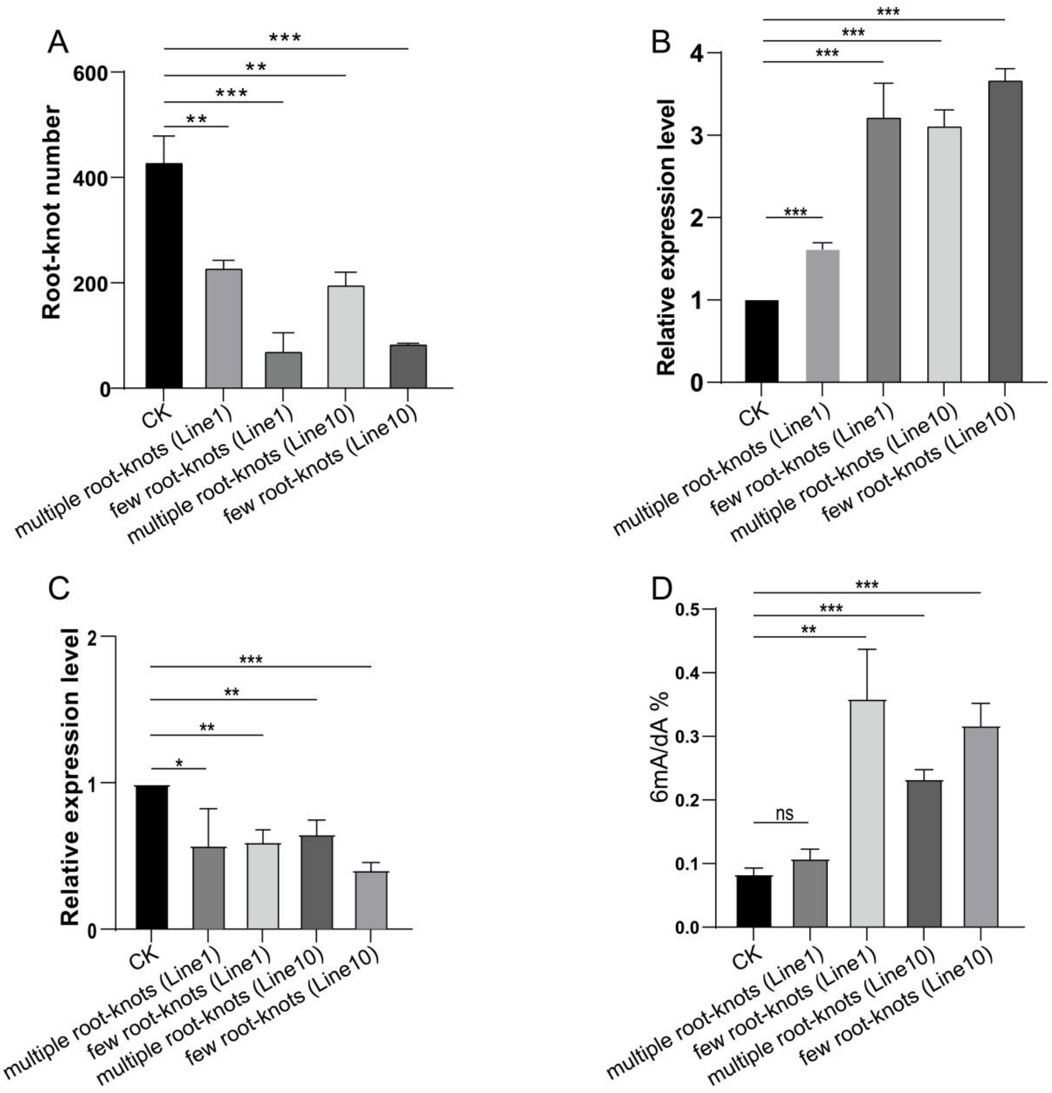


**Fig. S27.**

**Expression of *minmad-1* dsRNA confers resistance to *M. incognita* in transgenic tomato.** (*A*) Number of root galls formed on different transgenic tomato lines expressing *minmad-1* dsRNA compared with control (CK) plants, three weeks after inoculation with 800 *M. incognita* J2s. (*B*) Relative transcript levels of minmad-1 in the roots of different transgenic tomato lines expressing minmad-1 dsRNA, assessed by qRT-PCR. Expression levels are normalized to the tomato reference gene *actin-1* and presented relative to CK plants. (*C*) Relative expression levels of the *M. incognita minmad-1* gene isolated from few and multiple galls transgenic lines. (*D*) The DNA 6mA content of *M. incognita* egg isolated from few and multiple galls transgenic lines. $p$ values were calculated by two-tailed unpaired Student's $t$-test. One star * means $p < 0.05$, two means $p < 0.01$, three means $p < 0.001$.

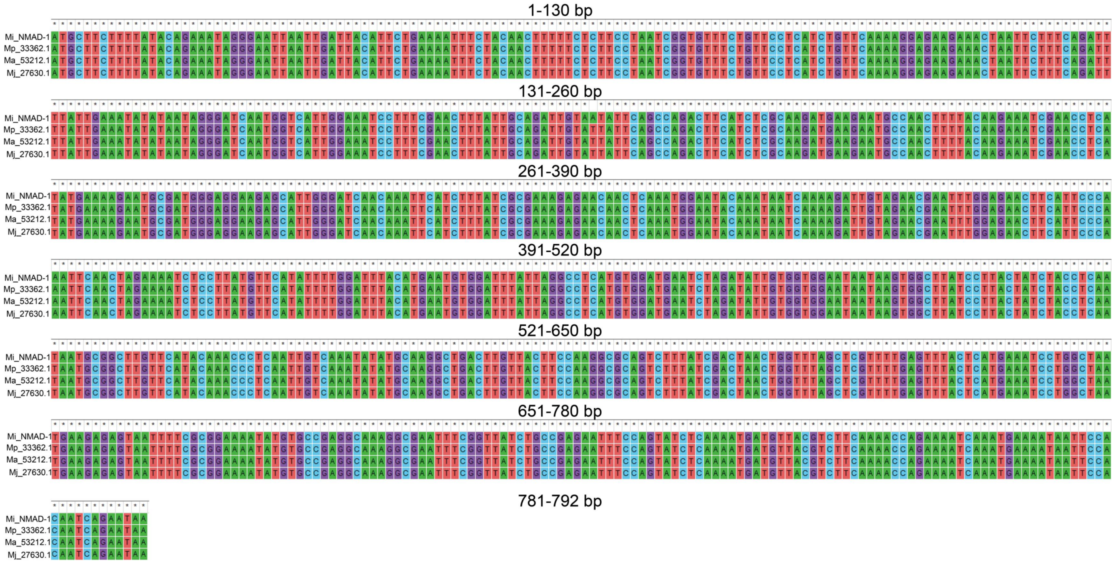


**Fig. S28.**

**The sequences alignment of *minmad-1* among polyploid RKNs (Mi_NMAD-1 for *M. incognita*, Mp_33362.1 for *M. arenaria* 3n, Ma_53212.1 for *M. arenaria* 4n, and Mj_27630.1 for *M. javanica*), the interference fragment is located at 481-787 bp.**

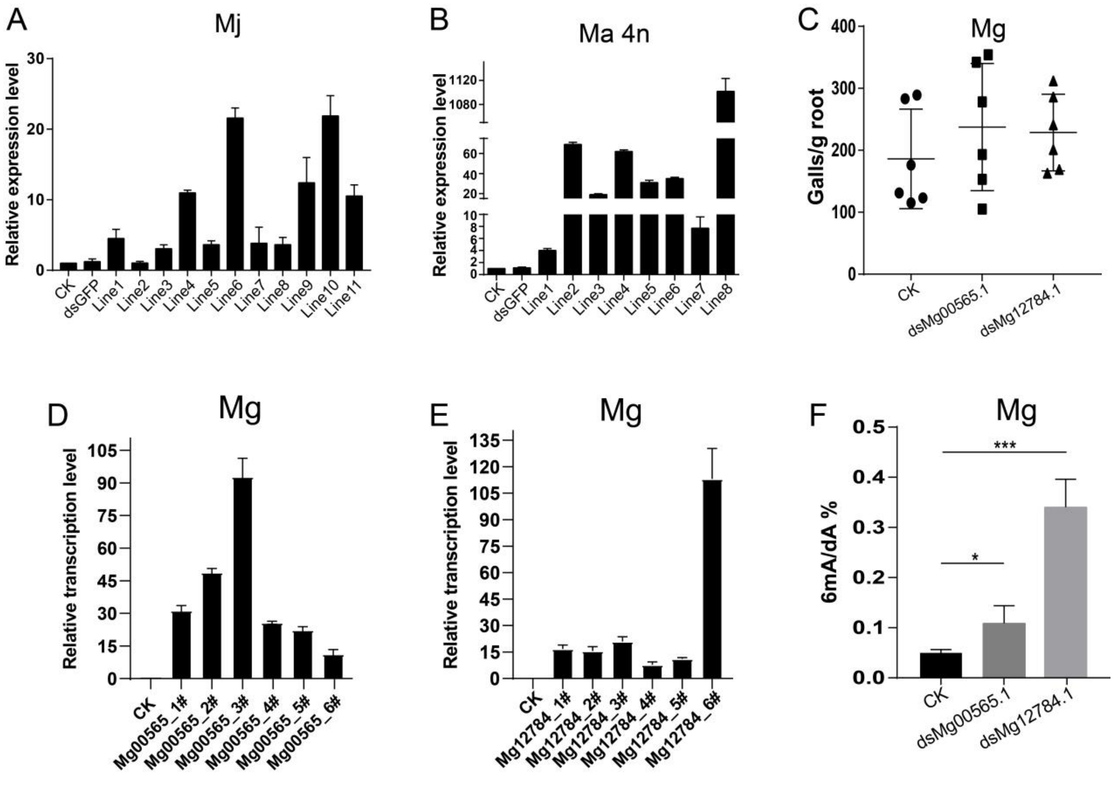


**Fig. S29.**

**Changes in the infection ability and 6mA content of tetraploid and diploid RKNs after knocking down the demethylase transcript level.** (*A*) Relative expression of *minmad-1* gene in transgenic tobacco resistant to Mj. (*B*) Relative expression of *minmad-1* gene in transgenic tobacco resistant to Ma 4n. (*C*) The number of galls of 1000 *M. graminicola* (Mg) J2 infected two months of transgenic rice expressing Mg demethylase gene dsRNA. (*D*, *E*) Relative expression levels of target genes in transgenic rice. (*F*) The content of genomic DNA 6mA of Mg after infecting transgenic rice was significantly higher than that of the control. *p* values were calculated by two-tailed unpaired Student's *t*-test. One star * means $p < 0.05$, two means $p < 0.01$, three means $p < 0.001$.

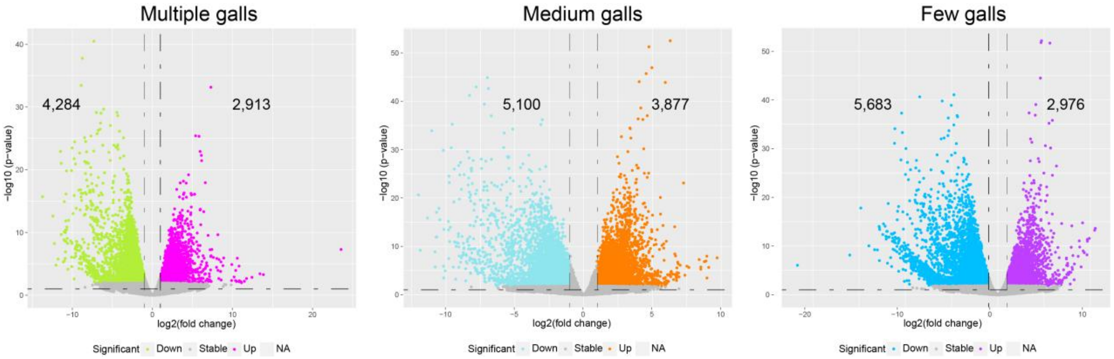


**Fig. S30.**

**The differentially expressed genes (DEGs) after *M. incognita* infected the transgenic tobacco expressing *minmad-1* dsRNA are shown in the volcano plot.** After Mi infected transgenic tobacco expressing *minmad-1* dsRNA, these infected tobaccos were divided into three groups according to the number of galls, and Mi eggs from each group were collected and RNA-seq was performed. Through comparative analysis of the RNA-seq data of each group and the RNA-seq data of the control group, the DEGs of these three groups were finally obtained. For each picture, the left side of the black dotted line represents the down-regulated DEGs, and the right side represents the up-regulated DEGs. The specific DEGs number were marked in the corresponding areas.

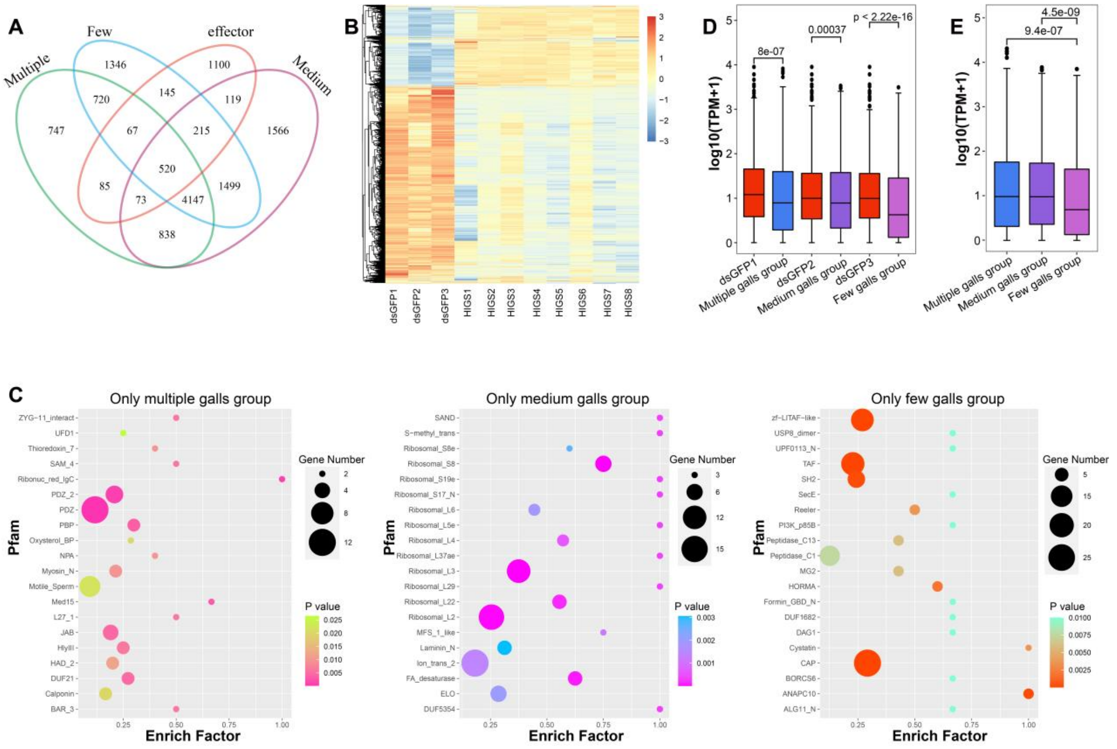


**Fig. S31.**

**The expression characteristic of multiple, medium, and few root-knot group DEGs.** (*A*) Venn diagram showing the relationship between multiple, medium, and few galls group DEGs and secreted protein genes. The unique DEGs for each group should be calculated as the number shown in the Venn diagram plus the portion overlapping specifically with effectors (e.g., 747 + 85 = 832 for the multiple-galls group). (*B*) The analysis of heatmap cluster for the multiple, medium, and few galls group shared DEGs, with two opposite expression clusters for the control and treatment groups. Among them, dsGFP represents the control group in which nematodes infected dsGFP-transgenic tobacco, while HIGS1–8 represent the experimental groups in which nematodes infected ds*minmad-1*-transgenic plants. (*C*) The Pfam enrichment analysis of multiple, medium, and few galls group unique DEGs, a large number of ribosomal proteins were enriched. (*D*) Comparison of the effector expression levels of the dsGFP control group and the experimental group in the multiple, medium and few galls group. (*E*) Comparison of the effector expression levels of multiple, medium and few galls group.

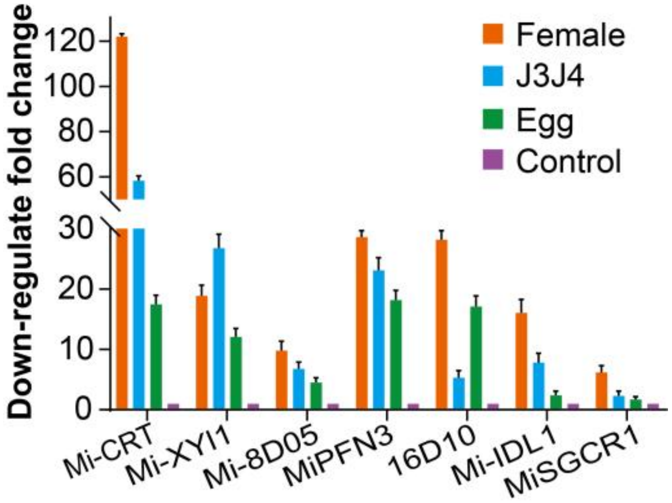


**Fig. S32.**
**The down-regulate fold change of several effectors in *M. incognita* eggs, J3J4 and female after infected with *minmad-1* transgenic tobacco were determined by qRT-PCR.**

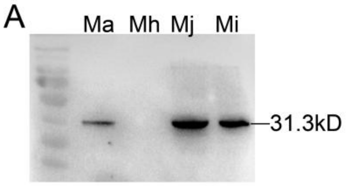


B

| Species | number of PE | PE identified by ChIP-seq |
|---|---|---|
| *M. incognita* | 2,324 | 938 |
| *M. javanica* | 5,324 | 2,820 |
| *M. arenaria* | 6,094 | 3,150 |

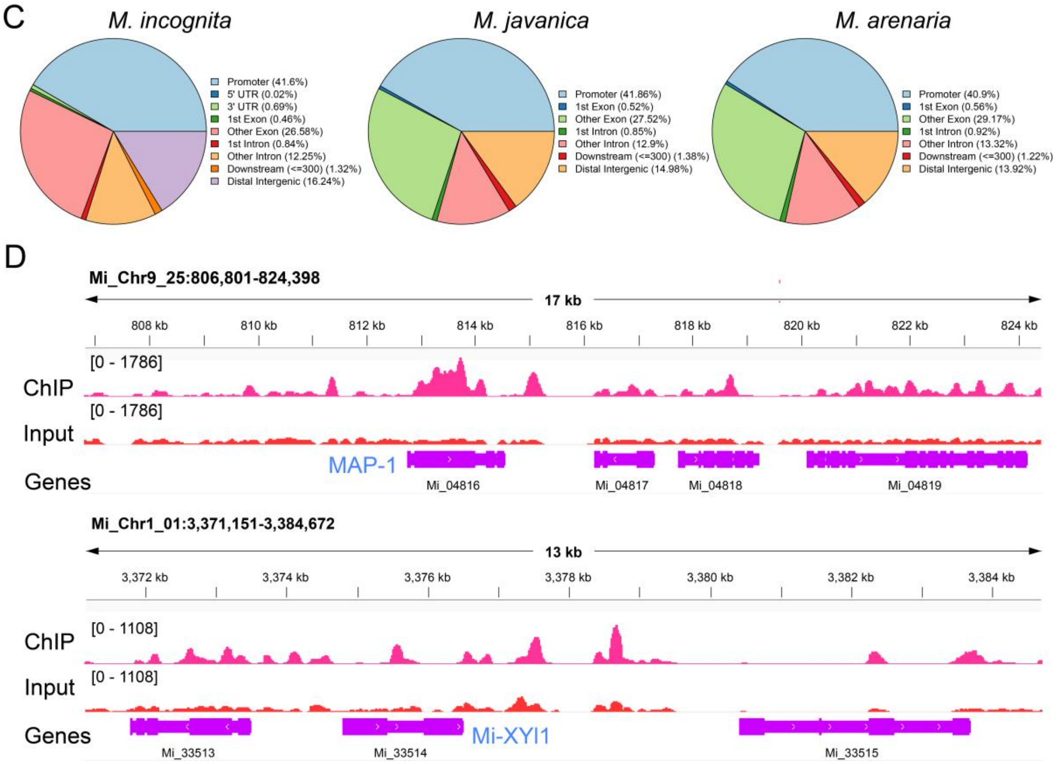


**Fig. S33.**

**ChIP-seq analysis of demethylases in polyploid RKNs reveals direct interactions with effector genes.** (*A*) Western blot analysis showing that the MiNMAD-1 antibody specifically recognizes demethylases from *M. incognita* (Mi), *M. arenaria* (Ma), and *M. javanica* (Mj), but not from *M. hapla* (Mh), which has substantial sequence divergence. (*B*) Number of effector genes identified by ChIP-seq in J2 juveniles of the three polyploid RKN species. PE means putative effector. (*C*) Genomic distribution of ChIP-seq peaks in each corresponding species. (*D*) Representative examples of known effectors identified by ChIP-seq.

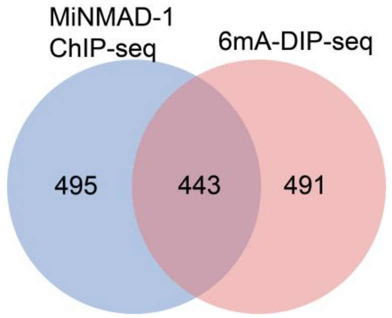


**Fig. S34.**
**Venn diagram of effector genes in *M. incognita* J2 identified by demethylase ChIP-seq and 6mA-DIP-seq, showing that approximately 40% of the genes overlap between the two datasets.**

**Table S1. Nematode materials involved in this study and research overview**

| species | LC-MS/MS | PacBio | Dot blot | 6mA related gene analysis | 6mA-DIP-seq | Identification of demethylas | Functional study of demethylase |
|---|---|---|---|---|---|---|---|
| *M. incognita* | Yes | Yes | Yes | Yes | Yes | Yes | Yes |
| *M. hapla* | Yes | Yes | Yes | Yes | No | Yes | No |
| *M. graminicola* | Yes | Yes | Yes | Yes | No | Yes | No |
| *M. arenaria* 3n | Yes | Yes | Yes | Yes | No | Yes | No |
| *M. arenaria* 4n | Yes | Yes | Yes | Yes | No | Yes | No |
| *M. javanica* | Yes | Yes | Yes | Yes | No | Yes | No |
| *C. elegans* | Yes | Yes[a] | Yes[a] | Yes | No | Yes[a] | No |
| *M. enterolobii* | Yes | No | No | No | No | No | No |
| *H. glycines* X12 | Yes | No | No | No | No | No | No |
| *H. glycines* race5 | Yes | No | No | No | No | No | No |
| *H. glycines* TN10 | Yes | No | No | No | No | No | No |
| *H. glycines* race3 | Yes | No | No | No | No | No | No |
| *D. destructor* | Yes | No | No | No | No | No | No |
| *A. avenae* | Yes | No | No | No | No | No | No |
| *B. xylophilus* | Yes | No | No | No | No | No | No |
| *H. contortus* | Yes | No | No | No | No | No | No |
| *C. nigoni* | Yes | No | No | No | No | No | No |

[a]Greer EL, et al. DNA Methylation on N6-Adenine in *C. elegans*. *Cell*. 2015 May 7;161(4):868-78. GEO accession number GSE66504.

The rest of the unmarked ones are completed by this study. *M. incognita* was selected as a representative for subsequent functional studies.

**Table S2. Summary of the data used in this study**

| Species | PacBio Sequel I | PacBio Revio | Hi-C | RNA-seq | WGA HiFi | 6mA/WGA -DIP-seq | Illumina | RNA-seq of demethylase RNAi |
|---|---|---|---|---|---|---|---|---|
| *M. incognita* | download SRR23517959 | this study SRR31956859 | / | download SRR17162178-80 | this study SRR31954495 | this study SRX25674687-90 | / | this study SRR22190500-22 |
| *C. elegans* | / | / | / | download SRR6125151-3 | / | / | / | / |
| *M. graminicola* | download SRR23517956 | / | / | download SRR17162217/29/30 | this study SRR31954491 | / | / | / |
| *M. arenaria 3n* | download | / | / | download SRR17162224/5/7 | / | / | / | / |
| *M. arenaria 4n* | download SRR23517958 | this study SRR31956858 | / | download SRR17162192-4 | this study SRR31954494 | / | / | / |
| *M. javanica* | download SRR23517957 | this study SRR31956857 | / | Download SRR17162208/9/11 | this study SRR31954493 | / | / | / |
| *M.hapla* | this study SRR22137913 | / | this study SRR22137914 | this study SRR22137906-11 | this study SRR31954492 | / | this study SRR22137912 | / |

Note: Except for *C. elegans*, all downloaded data are from our previous research papers (Dai et al., Nature Comminications, 2023). The meaning of multiple SRA numbers is, for example, SRR17162178-80 means SRR17162178, SRR17162179, SRR17162180; and SRR17162217/29/30 means including SRR17162217, SRR17162229, SRR17162230.

**Table S3. Summary of PacBio reads contamination rate**

| Species | Total reads | Aligned to bacteria | Bacteria ratio | Aligned to fungi | Fungi ratio | Aligned to plant | Plant ratio |
|---|---|---|---|---|---|---|---|
| ***M. incognita*** | 1,709,111 | 1,131 | 0.0662% | 12 | 0.0007% | 1,134 | 0.0664% |
| ***M. javanica*** | 1,954,715 | 8 | 0.0004% | 24 | 0.0012% | 16 | 0.0008% |
| ***M. arenaria* 3n** | 2,015,633 | 841 | 0.0417% | 19 | 0.0009% | 53 | 0.0026% |
| ***M. arenaria* 4n** | 3,037,368 | 44 | 0.0014% | 23 | 0.0008% | 66 | 0.0022% |
| ***M. graminicola*** | 1,009,419 | 249 | 0.0247% | 5 | 0.0005% | 13 | 0.0012% |
| ***M. hapla*** | 1,075,982 | 1,207 | 0.1122% | 22 | 0.0020% | 263 | 0.0244% |

**Table S4. The proportion and sources of contamination in *M. incognita* hifi reads evaluated by 6mASCOPE**

| #total_CCS | mapped_to_goi | contaminants |
|---|---|---|
| 231400 | 231209 (99.9175%) | 191 (0.0825411%) |

| Count | Species |
|---|---|
| 31 | Bosea sp. |
| 14 | Solanum lycopersicum |
| 6 | Heligmosomoides polygyrus |
| 1 | Uncultured bacterium |
| 1 | Trichoderma citrinoviride |
| 1 | Thermus scotoductus |
| 1 | Rhodopseudomonas palustris |
| 1 | Purpureocillium lilacinum |
| 1 | Pseudolabrys taiwanensis |
| 1 | PREDICTED: Zonotrichia |
| 1 | PREDICTED: Tetranychus |
| 1 | PREDICTED: Strongylocentrotus |
| 1 | PREDICTED: Spodoptera |
| 1 | PREDICTED: Paramormyrops |
| 1 | PREDICTED: Ornithorhynchus |
| 1 | PREDICTED: Microtus |
| 1 | PREDICTED: Megachile |
| 1 | PREDICTED: Echinops |
| 1 | PREDICTED: Drosophila |
| 1 | PREDICTED: Atta |
| 1 | Nippostrongylus brasiliensis |
| 1 | Microbacterium sp. |
| 1 | Caenorhabditis sp. |
| 1 | Caenorhabditis elegans |
| 1 | Bradyrhizobium sp. |
| 1 | Bradyrhizobium lablabi |
| 1 | Bradyrhizobium canariense |
| 1 | Bacillus phage |
| 1 | Angiostrongylus costaricensis |
| 1 | Actinoplanes sp. |
| 1 | Actinoplanes friuliensis |

Note: The proportion of reads from bacterial and fungal contamination is 0.018%.

**Table S5. The proportion and sources of contamination in *M. arenaria* 4n hifi reads evaluated by 6mASCOPE**

| #total_CCS | mapped_to_goi | contaminants |
|---|---|---|
| 210593 | 210335 (99.8775%) | 258 (0.122511%) |

| Count | Species |
|---|---|
| 18 | Bosea sp. |
| 15 | Meloidogyne hapla |
| 7 | Meloidogyne paranaensis |
| 5 | Meloidogyne incognita |
| 1 | Sphingobacterium mizutaii |
| 1 | Rhodopseudomonas palustris |
| 1 | Rhabditis tokai |
| 1 | Heterodera glycines |
| 1 | Caenorhabditis sp. |
| 1 | Bradyrhizobium sp. |
| 1 | Bradyrhizobium oligotrophicum |
| 1 | Bradyrhizobium erythrophlei |

Note: The proportion of reads from bacterial and fungal contamination is 0.011%.

**Table S6. The proportion and sources of contamination in *M. javanica* hifi reads evaluated by 6mASCOPE**

| #total_CCS | mapped_to_goi | contaminants |
|---|---|---|
| 259546 | 259266 (99.8921%) | 280 (0.107881%) |

| Count | Species |
|---|---|
| 30 | Meloidogyne incognita |
| 28 | Meloidogyne hapla |
| 22 | Bosea sp. |
| 16 | Meloidogyne paranaensis |
| 6 | PREDICTED: Nicotiana |
| 2 | Rhodopseudomonas palustris |
| 1 | Uncultured bacterium |
| 1 | Tobacco vein-clearing |
| 1 | Sphingomonas indica |
| 1 | Panurginus albopilosus |
| 1 | Nippostrongylus brasiliensis |
| 1 | Nicotiana tabacum |
| 1 | Cutaneotrichosporon oleaginosus |
| 1 | Bradyrhizobium sp. |
| 1 | Bacillus sp. |

Note: The proportion of reads from bacterial and fungal contamination is 0.011%.

**Table S7. Summary of chromosome-level assembly for *M. hapla***

| **Contig** | |
|---|---|
| Total assembly size (bp) | 52,949,056 |
| Number of contigs | 241 |
| Contig N50 (bp) | 342,689 |
| **Scaffolds** | |
| Total assembly size (bp) | 53,139,056 |
| Number of scaffolds | 51 |
| scaffolds N50 (bp) | 2,897,429 |
| Gaps | 190 |
| Length of contigs in scaffolds | 99.18% |
| Chromosome size (bp) | |
| Mh_chr1 | 8,766,289 |
| Mh_chr2 | 4,630,434 |
| Mh_chr3 | 3,898,789 |
| Mh_chr4 | 3,228,195 |
| Mh_chr5 | 3,031,161 |
| Mh_chr6 | 2,897,429 |
| Mh_chr7 | 2,973,021 |
| Mh_chr8 | 2,878,996 |
| Mh_chr9 | 2,838,729 |
| Mh_chr10 | 2,521,355 |
| Mh_chr11 | 2,475,582 |
| Mh_chr12 | 2,163,482 |
| Mh_chr13 | 2,295,975 |
| Mh_chr14 | 2,382,878 |
| Mh_chr15 | 2,237,739 |
| Mh_chr16 | 2,022,336 |

**Table S8. Summary of sequencing data of Mh**

| Species | Library | Total Data | Average length | Sequence depth |
|---|---|---|---|---|
| ***M. hapla*** | Illumina | 11 Gb | 150 bp | ~216× |
| | PacBio | 8.6 Gb | 4308 bp | ~169× |
| | Hi-C | 23 Gb | 150 bp | ~453× |

### Table S9. Summary of 6mA identified by SMRT sequencing

| species | A count in genome | 6mA count | Coverage cutoff | 6mA count after cutoff | Filtered by WGA | 6mA count after WGA | 6mA/A |
|---|---|---|---|---|---|---|---|
| ***M. incognita*** | 74,388,433 | 500,236 | 40× | 315,880 | 221 | 315,659 | 0.424% |
| ***M. arenaria* 3n** | 81,251,221 | 351,719 | 32× | 254,774 | / | 254,774 | 0.314% |
| ***M. arenaria* 4n** | 108,528,702 | 608,756 | 41× | 377,129 | 0 | 377,129 | 0.347% |
| ***M. javanica*** | 107,728,037 | 266,650 | 35× | 252,676 | 2,705 | 249,970 | 0.232% |
| ***M. graminicola*** | 17,532,413 | 49,216 | 50× | 48,659 | 1,569 | 47,090 | 0.269% |
| ***M. hapla*** | 19,265,712 | 15,776 | 15× | 12,858 | 251 | 12,607 | 0.065% |

Note: The cut-off index used in this study refers to the method reported by previous researchers (Stephen J Mondo et al, Nature Genetics, 2017 and Chao Zhou et al, Nature Plants, 2018). In short, the sites with modification score of <20 are first filtered out, and then the sites with lower confidence coverage multiples are removed from the remaining sites based on the coverage distribution.

**Table S10. Comparison of the number of 6mA sites predicted by the same analysis method in WGA-DNA and native-DNA.**

| Sample name | Reads number | Number of 6mA sites | WGA 6mA/native 6mA |
|---|---|---|---|
| *M. incognita* | 1,709,111 | 315,880 | 0.32% |
| *M. incognita* WGA | 95,520 | 999 | |
| *M. graminicola* | 1,009,419 | 48,659 | 13.34% |
| *M. graminicola* WGA | 186,151 | 6,492 | |
| *M. Arenaria* 4n | 3,037,368 | 377,129 | 0.06% |
| *M. arenaria* 4n WGA | 139,620 | 238 | |
| *M. javanica* | 1,954,715 | 252,676 | 5.42% |
| *M. javanica* WGA | 357,670 | 13,694 | |
| *M. hapla* | 1,075,982 | 12,858 | 25% |
| *M. hapla* WGA | 298,794 | 3,214 | |

**Table S11. Contamination ratio of WGA samples evaluated by 6mASCOPE.**

| | | |
|---|---|---|
| | *M. incognita* | |
| #total_CCS | mapped_to_goi | contaminants |
| 74807 | 72375 (96.749%) | 2432 (3.25103%) |
| | *M. arenaria* | |
| #total_CCS | mapped_to_goi | contaminants |
| 131571 | 131420 (99.8852%) | 151 (0.114767%) |
| | *M. javanica* | |
| #total_CCS | mapped_to_goi | contaminants |
| 319915 | 316269 (98.8603%) | 3646 (1.13968%) |
| | *M. hapla* | |
| #total_CCS | mapped_to_goi | contaminants |
| 242355 | 200674 (82.8017%) | 41681 (17.1983%) |
| | *M. graminicola* | |
| #total_CCS | mapped_to_goi | contaminants |
| 147837 | 135870 (91.9053%) | 11967 (8.09473%) |

Note: During 6mASCOPE analysis, possible inter-species chimeric reads were removed for further analysis, so the number of total_CCS reads here is lower than that the real reads number.

**Table S12. Comparison of 6mA peak numbers obtained by 6mA-DIP-seq of WGA samples and native DNA samples**

| Sample name | Reads number | 6mA peak number |
|---|---|---|
| Mi-WGA-6mAIP-rep1 | 21,863,290 | 41 |
| Mi-WGA-6mAIP-rep2 | 18,878,136 | 30 |
| Mi-6mAIP-rep1 | 80,301,822 | 17,804 |
| Mi-6mAIP-rep2 | 40,113,393 | 16,924 |

**Table S13. Statistics of the transposons of five species of RKNs (genome v1 in our previous study) and *C. elegans* ce10**

| Type | Mj | Mi | Ma 3n | Ma 4n | Mh | Mg | Ce |
|---|---|---|---|---|---|---|---|
| DNA/DTH % | 0.18 | 0.33 | 0.21 | 0.24 | 0 | 0 | 0.09 |
| DNA/DTA % | 8.15 | 7.03 | 7.32 | 8.95 | 1.36 | 0.56 | 0.66 |
| DNA/DTT % | 0.3 | 0.26 | 0.14 | 0.31 | 0.02 | 0.01 | 0.03 |
| DNA/DTM % | 8.44 | 6.64 | 7.91 | 8.14 | 0.62 | 0.36 | 10.16 |
| DNA/Helitron % | 0.82 | 1.28 | 0.56 | 0.78 | 0.63 | 0.16 | 2.00 |
| DNA/DTC % | 0.57 | 0.45 | 0.66 | 0.98 | 0.13 | 0.01 | 1.18 |
| **SUM_DNA %** | **18.46** | **15.99** | **16.8** | **19.4** | **2.76** | **1.1** | **14.13** |
| LTR/Gypsy % | 3.4 | 3.42 | 3.4 | 4.3 | 1.92 | 0.37 | 0.4 |
| LTR/unknown % | 3.09 | 2.55 | 2.42 | 3.25 | 0.92 | 0 | 0.45 |
| LTR/Copia % | 0.03 | 0.01 | 0.01 | 0.05 | 0.03 | 0 | 0.01 |
| **SUM_LTR %** | **6.52** | **5.98** | **5.83** | **7.6** | **2.87** | **0.37** | **0.86** |
| MITE/DTT % | 0.01 | 0.01 | 0 | 0 | 0.004 | 0 | 0.001 |
| MITE/DTA % | 2.12 | 2 | 1.91 | 2.1 | 0.36 | 0.07 | 0.65 |
| MITE/DTM % | 1.66 | 1.42 | 1.55 | 1.14 | 0.12 | 0.12 | 0.53 |
| MITE/DTC % | 0.02 | 0.01 | 0.02 | 0.01 | 0.27 | 0.01 | 0.02 |
| MITE/DTH % | 0.09 | 0.01 | 0.09 | 0.07 | 0.08 | 0 | 0 |
| **SUM_MITE %** | **3.9** | **3.45** | **3.57** | **3.32** | **0.72** | **0.2** | **1.21** |
| **SUM %** | **28.88** | **25.42** | **26.2** | **30.32** | **6.35** | **1.67** | **16.2** |
| Low_complexity % | 2.67 | 2.76 | 2.66 | 2.61 | 4.27 | 6.75 | 1.43 |
| **Total %** | **31.55** | **28.18** | **28.86** | **32.93** | **10.62** | **8.42** | **17.63** |

**Table S14. Summary of RNA-seq data on the J3J4 of eight treatments of MiNMAD-1 RNAi**

| RNA-seq sample | Clean paired reads | Aligned concordantly exactly 1 time | Aligned concordantly >1 time | Unaligned reads | SRA number |
|---|---|---|---|---|---|
| **J3J4RNAi_treat1** | 27,851,656 | 12,948,826 (46.49%) | 13,309,674 (47.79%) | 1,571,771 (5.64%) | SRR22190515 |
| **J3J4RNAi_treat2** | 24,966,530 | 17,904,762 (71.72%) | 6,030,962 (24.16%) | 994,333 (3.98%) | SRR22190514 |
| **J3J4RNAi_treat3** | 26,653,509 | 19,535,061 (73.29%) | 6,102,398 (22.90%) | 957,886 (3.59%) | SRR22190513 |
| **J3J4RNAi_treat4** | 20,817,461 | 15,459,866 (74.26%) | 4,620,374 (22.19%) | 694,714 (3.34%) | SRR22190512 |
| **J3J4RNAi_treat5** | 23,345,219 | 15,744,981 (67.44%) | 5,705,538 (24.44%) | 2,371,371 (10.16%) | SRR22190511 |
| **J3J4RNAi_treat6** | 24,993,967 | 18,275,506 (73.12%) | 5,885,822 (23.55%) | 791,036 (3.16%) | SRR22190509 |
| **J3J4RNAi_treat7** | 27,535,906 | 19,239,465 (69.87%) | 7,244,043 (26.31%) | 1,015,747 (3.69%) | SRR22190508 |
| **J3J4RNAi_treat8** | 26,860,532 | 19,452,222 (72.42%) | 6,471,665 (24.09%) | 895,404 (3.33%) | SRR22190507 |
| **Control_1** | 23,695,921 | 17,359,842 (73.26%) | 5,681,057 (23.97%) | 614,436 (2.59%) | SRR17162173 |
| **Control_2** | 23,556,388 | 16,280,721 (69.11%) | 6,421,364 (27.26%) | 792,397 (3.36%) | SRR17162172 |
| **Control_3** | 37,855,109 | 27,164,449 (71.76%) | 9,698,810 (25.62%) | 929,266 (2.45%) | SRR17162171 |

**Table S15. Summary of qRT-PCR primer of the verified effector used in this study**

| effector | Primer F | Primer R |
|---|---|---|
| **Mi-XYl1** | CGTGTCGGAATTGTTGATTTATGT | TTGTGCTGTTAATGCTTCTTGTC |
| **Mi-CRT** | GATGCTCGCTTCTATAGTATTTC | GAGGCCATGAGCTAAATTGATTC |
| **Mi8D05** | TTCCACCACAACAGCCACCTT | GACTGCCAGCAAGACCTCCTC |
| **MiPFN3** | GGAACTGGCCATGTTTCAAAGGC | GTCCATTCGCTGCAGCATTTGC |
| **16D10** | GCCTTTAATGGTTACTTTAATGC | TCAATTATTTCCTCCAGGATTTGG |
| **MiSGCR1** | GGAATCGGTGGCTTTGGT | CTCCTCCGCATCCTCCATA |
| **MiIDL1** | GCTTTTATCTGTCTCAATTGTGG | CGGCCGGGACCTGGAACTTT |
